# Transcriptome and metabolome analyses revealed that narrowband 280 and 310 nm UV-B induce distinctive responses in Arabidopsis

**DOI:** 10.1101/2021.02.12.430709

**Authors:** Tomohiro Tsurumoto, Yasuo Fujikawa, Daisaku Ohta, Atsushi Okazawa

## Abstract

In plants, the UV-B photoreceptor UV RESISTANCE LOCUS8 (UVR8) perceives UV-B and induces UV-B responses including synthesis of UV-B absorbing phenolic compounds such as anthocyanins. UVR8 absorbs a range of UV-B (260–335 nm). However, the responsiveness of plants to each UV-B wavelength has not been intensively studied so far. Here, we performed transcriptome and metabolome analyses of Arabidopsis using UV light emitting diodes (LEDs) with peak wavelengths of 280 and 310 nm to investigate the differences in the wavelength-specific UV-B responses. Irradiation with both UV-LEDs induced gene expression of the transcription factor ELONGATED HYPOCOTYL 5 (HY5), which has a central role in the UVR8 signaling pathway. However, the overall transcriptomic and metabolic responses to 280 and 310 nm UV-LED irradiation were different. Most of the known UV-B-responsive genes, such as salicylic acid, jasmonic acid, and defense-related genes, responded only to 280 nm UV-LED irradiation. Lipids, polyamines and organic acids were the metabolites most affected by 280 nm UV-LED irradiation, whereas the effect of 310 nm UV-LED irradiation on the metabolome was considerably less. Enzymatic genes involved in the phenylpropanoid pathway upstream in anthocyanin biosynthesis were up-regulated only by 280 nm UV-LED irradiation. On the other hand, no enzymatic genes downstream in anthocyanin biosynthesis were induced by the UV-LEDs, but rather, they were down-regulated by 310 nm UV-LED irradiation. These results revealed that the responsivenesses of Arabidopsis to 280 and 310 nm UV-B were significantly different, suggesting that UV-B signaling is mediated by more complex pathways than the current model.

## INTRODUCTION

The sunlight that falls on the earth contains various wavelengths. Plants use the light not only as an energy source, but also as signals for optimizing growth and development. Light is classified according to the wavelengths it contains, and the shortest-wavelength component of the sunlight that reaches the ground is UV. UV is further divided into three bands, UV-A (315–400 nm), UV-B (280–315 nm), and UV-C (100–280 nm). Since UV-C is absorbed by the stratospheric ozone, in the sunlight that reaches the ground, UV-B is the light that possesses the shortest wavelength and accordingly the highest energy. Because nucleic acids and proteins, have absorption peaks near 260 and 280 nm, respectively, UV-B directly causes damage to the molecules. UV-B also produces reactive oxygen species (ROS) that cause damage to DNAs, proteins, and lipids, and induce many processes like programmed cell death (PCD), abiotic stress responses and pathogen defense (Gill and Tuteja, 2010; Hideg *et al*., 2013). Plants have adapted to this harmful UV-B during their evolution, acquiring plant-specific UV-B responses (Jenkins, 2009; Dotto and Casati, 2017; Yin and Ulm, 2017). As a typical UV-B response, the levels of phenolic compounds, such as anthocyanins, that absorb UV-B increase (Rozema *et al*., 1997; Heijde and Ulm, 2012).

The UV-B receptor UV RESISTANCE LOCUS 8 (UVR8) mediates the UV-B response in plants (Rizzini et al., 2011; Heijde and Ulm, 2012; Jenkins, 2017; Yin and Ulm, 2017). UV-B signaling requires monomerization of UVR8, which forms a homodimer in the ground state (Rizzine *et al*., 2011). The absorption spectrum of UVR8 has a peak at 280 nm and extends from at least 250 nm to around 500 nm (Rai *et al*., 2020). Monomerization of UVR8 occurs under UV in the range from 260 to 335 nm (Díaz-Ramos *et al*., 2018; Rai *et al*., 2020). UVR8 monomers interact with CONSTITUTIVE PHOTOMORPHOGENIC1 (COP1), an E3 ubiquitin ligase, and inactivate COP1 activity, resulting in stabilization of ELONGATED HYPOCOTYL5 (HY5), a member of the bZIP transcription factor family, that inhibits hypocotyl growth and lateral root development, and promotes pigment accumulation (Heijde and Ulm, 2012; Gangappa and Botto, 2016; Jenkins, 2017; Yin and Ulm, 2017). HY5 has a central role in UV-B signaling, regulating the expression of about half of UVR8-induced genes, including defense-related and anthocyanin biosynthetic genes (Brown *et al*., 2005; Vandenbussche *et al*., 2018).

Metabolomic analysis of the UV-B response in Arabidopsis showed increased amounts of phenolic compounds, such as sinapoyl malate, and quercetin- and kaempferol-glycosides (Lake *et al*., 2009). Reprogramming of both central and specialized metabolisms in response to UV-B was also shown with increases in sugars, amino acids, and organic acids in the tricarboxylic acid (TCA) cycle, and phenolic compounds, including anthocyanins (Kusano *et al*., 2011). Anthocyanins, which act as UV sunscreen in plants, are synthesized through the biosynthetic pathways of phenylpropanoids and flavonoids. The biosynthesis of phenylpropanoids starts from L-phenylalanine, which is synthesized by arogenate dehydrotase (ADT) (Fraser and Chapple, 2011). *p*-Coumaroyl CoA, a precursor of flavonoids biosynthesis, is produced by L-phenylalanine ammonia-lyase (PAL), cinnamate 4-hydroxylase (C4H) and 4-coumarate-CoA ligase (4CL) in the phenylpropanoid pathway. Chalcone synthase (CHS), chalcone isomerase (CHI), flavonol 3-hydroxylase (F3H) and flavonol 3′-hydroxylase (F3′H) are main enzymes in flavonoid biosynthesis. Dihydroflavonol-4-reductase (DFR) and leucoanthocyanidin dioxygenase/anthocyanidin synthase (LDOX/ANS) are involved in synthesis of the basic skeleton of anthocyanin, and multiple anthocyanin glycosyltransferases (AGTs) modify the sugar moieties of anthocyanins. Expression of these enzymatic genes is regulated by several classes of transcription factors, such as MYB-bHLH-WDR (MBW) complexes (Broun 2005; Xu *et al*., 2014, 2015; Ma and Constabel 2019), and coregulatory Mediator complexes (Bonawitz *et al.*, 2012; Wang *et al*., 2020). Flavonoid biosynthesis is up-regulated under UV-B through regulation of these transcription factors mediated by UVR8 signaling (Jenkins 2014; Ma and Constabel 2019; Qian *et al*., 2020).

Most of the previous studies on UV-B responses used broadband (BB) UV lamps with peaks around 310 nm and ranging from 280 to 380 nm (Brown *et al*., 2005; Lake *et al*., 2009; Kusano *et al*., 2011; Vandenbussche *et al*., 2018), although the maximal absorbance of UVR8 is at 280 nm. To our knowledge, the responsiveness of plants to each UV-B wavelength has not been intensively studied so far. Here, we performed transcriptome and metabolome analyses of Arabidopsis using UV light emitting diodes (LEDs) with peak wavelengths of 280 and 310 nm to investigate the UV-B responses under the narrowband (NB) UV-B light. Our results showed that the responsivenesses of Arabidopsis to 280 and 310 nm UV-B were significantly different.

## RESULTS

### Transcriptomic response to narrowband UV-B

In this study, 280 nm UV-LED with a peak wavelength of 280 nm, a half-width of 10 nm, and a range from 260 to 310 nm and 310 nm UV-LED with a peak wavelength of 310 nm, a half-width of 10 nm, and a range from 290 to 340 nm were used for irradiation (Figure 1). The photon flux density on the irradiated surface was adjusted to 2.5 μmol m^−2^ s^−1^. Arabidopsis grown under long-day conditions (16-h light/8-h dark) for 14 days at 23°C was used for the irradiation experiments. The irradiation conditions were: (i) 45 min irradiation with the 280 nm UV-LED (280-0d); (ii) 45 min irradiation with the 310 nm UV-LED (310-0d); (iii) 45 min irradiation with the 280 nm UV-LED followed by incubation in the dark for two days (280-2d); and (iv) 45 min irradiation with the 310 nm UV-LED followed by incubation in the dark for two days (310-2d). For the control, plants without UV-B irradiation (C-0d) and plants incubated in the dark for two days without UV-B irradiation (C-2d) were used.

**Figure 1.**
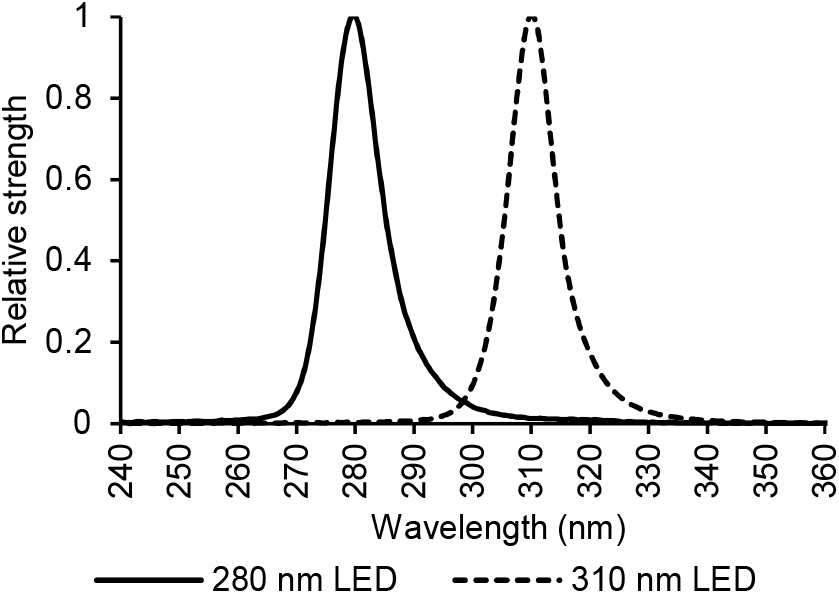
Relative emission spectra of 280 and 310 nm UV light emitting diodes (LEDs) used in the experiments. The 280 nm UV-LED has a peak wavelength of 280 nm, a half-width of 10 nm, and a range from 260 to 310 nm. The 310 nm UV-LED has a peak wavelength of 310 nm, a half-width of 10 nm, and a range from 290 to 340 nm.

To investigate the responses to NB UV-B, the transcriptome in Arabidopsis irradiated with the 280 or 310 nm UV-LED was analyzed using RNA-Seq. It was revealed that the expression of much fewer genes was affected by 310 nm UV-LED irradiation, especially in 310-2d, compared with 280 nm UV-LED irradiation. In comparison with the control, the numbers of differentially expressed genes (DEGs) were 3386, 1277, 3948, and 96 in 280-0d, 310-0d, 280-2d, and 310-2d, respectively (Figure 2a). Moreover, DEGs in Arabidopsis under 280 and 310 nm UV-LED were not closely overlapped as expected in terms of plant UV-B response. The numbers of commonly up-regulated DEGs between 280-0d and 310-0d and between 280-2d and 310-2d were 261 and 15, respectively, and the numbers of commonly down-regulated DEGs between 280-0d and 310-0d and between 280-2d and 310-2d were 487 and 1, respectively (Figure 2b, Tables S1 and S2).

**Figure 2.**
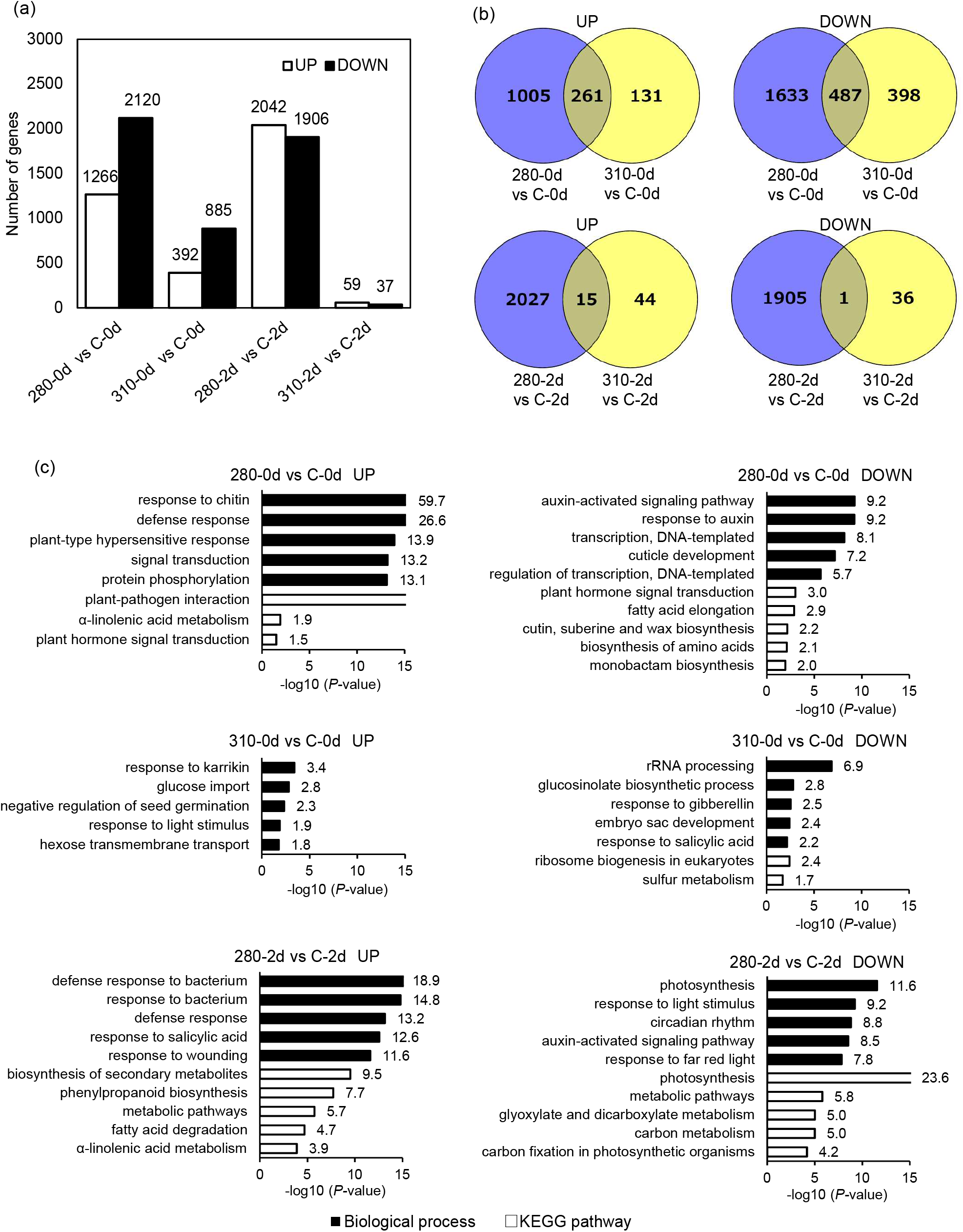
Overall description of transcriptome data. (a) Comparison of up-regulated and down-regulated numbers of DEGs under 280 and 310 nm UV. Welch’s *t*-test was applied for comparison of UV-irradiated samples with control, and a gene with a *P*-value less than 0.05 and a two-fold change was considered as a DEG. (b) Venn diagrams illustrating the shared and unique up-regulated and down-regulated DEGs between samples under 280 and 310 nm UV-B. (c) Gene ontology enrichment analysis of up-regulated and down-regulated DEGs in biological processes and Kyoto Encyclopedia of Genes and Genomes pathway using DAVID ver. 6.8. Top five terms were selected according to the *P*-values when more than five terms were enriched.

### Gene ontology term enrichment analysis of DEGs

To investigate the underlying functions of the DEGs in the responses to 280 and 310 nm UV-LED irradiation, gene ontology (GO) term enrichment analysis was conducted using the Database for Annotation, Visualization and Integrated Discovery (DAVID), version 6.8 (Huang *et al*., 2009a, 2009b). Enriched terms were totally different between the DEGs under 280 and 310 nm UV-LED irradiation in accordance with their slight overlapping. Biotic or abiotic stress-related terms, such as ‘response to chitin’, ‘defense response’, ‘defense response to bacterium’, and ‘response to salicylic acid’ were enriched in up-regulated DEGs in 280-0d and 280-2d. In contrast, the term ‘response to salicylic acid’ was enriched in down-regulated DEGs in 310-0d, and other stress-related terms were not enriched under 310 nm UV-LED irradiation. Auxin related genes in ‘auxin-activated signaling pathway’ and ‘response to auxin’ were down-regulated under 280 nm UV-LED irradiation. Genes related to photosynthesis and responses to light stimulus and far red light were down-regulated in 280-2d, whereas ‘response to light stimulus’ was enriched in DEGs in 310-0d. There was no enriched term in DEGs in 310-2d (Figure 2c and Table S3).

In the Kyoto Encyclopedia of Genes and Genomes (KEGG) pathways, up-regulated DEGs were most enriched in ‘plant-pathogen interaction’ and ‘biosynthesis of secondary metabolites’ in 280-0d and 280-2d, respectively. The term ‘phenylpropanoid biosynthesis’ was also enriched in DEGs in 280-2d. The terms related to lipid metabolism, such as ‘α-linolenic acid metabolism’, and ‘fatty acid degradation’ were characteristically enriched under 280 nm UV-LED irradiation, whereas ‘fatty acid elongation’ and ‘cutin, suberine and wax biosynthesis’ were down-regulated in 280-0d. Down-regulated DEGs were most enriched in ‘plant hormone signal transduction’, ‘ribosome biogenesis in eukaryotes’, and ‘photosynthesis’ in 280-0d, 310-0d, and 280-2d, respectively. There was no enriched term in up-regulated DEGs in 310-0d and 310-2d, and in down-regulated DEGs in 310-2d (Figure 2c and Table S3).

### Effect of narrowband UV-B on metabolome

The metabolome in 280-2d and 310-2d were analyzed using gas chromatography-mass spectrometry (GC-MS) and liquid chromatography-mass spectrometry (LC-MS) to investigate the effect of NB UV-B on the Arabidopsis metabolism. Overall, fewer metabolic changes were confirmed under 310 nm UV-LED irradiation as transcriptomic responses. GC-MS analysis showed that the amounts of 19 hydrophilic metabolites and seven fatty acids in 280-2d, and only two fatty acids in 310-2d, which were also increased in 280-2d, were increased in comparison with the control, whereas the amounts of two hydrophilic metabolites were decreased in 280-2d (Figure 3a,c and Table S3).

**Figure 3.**
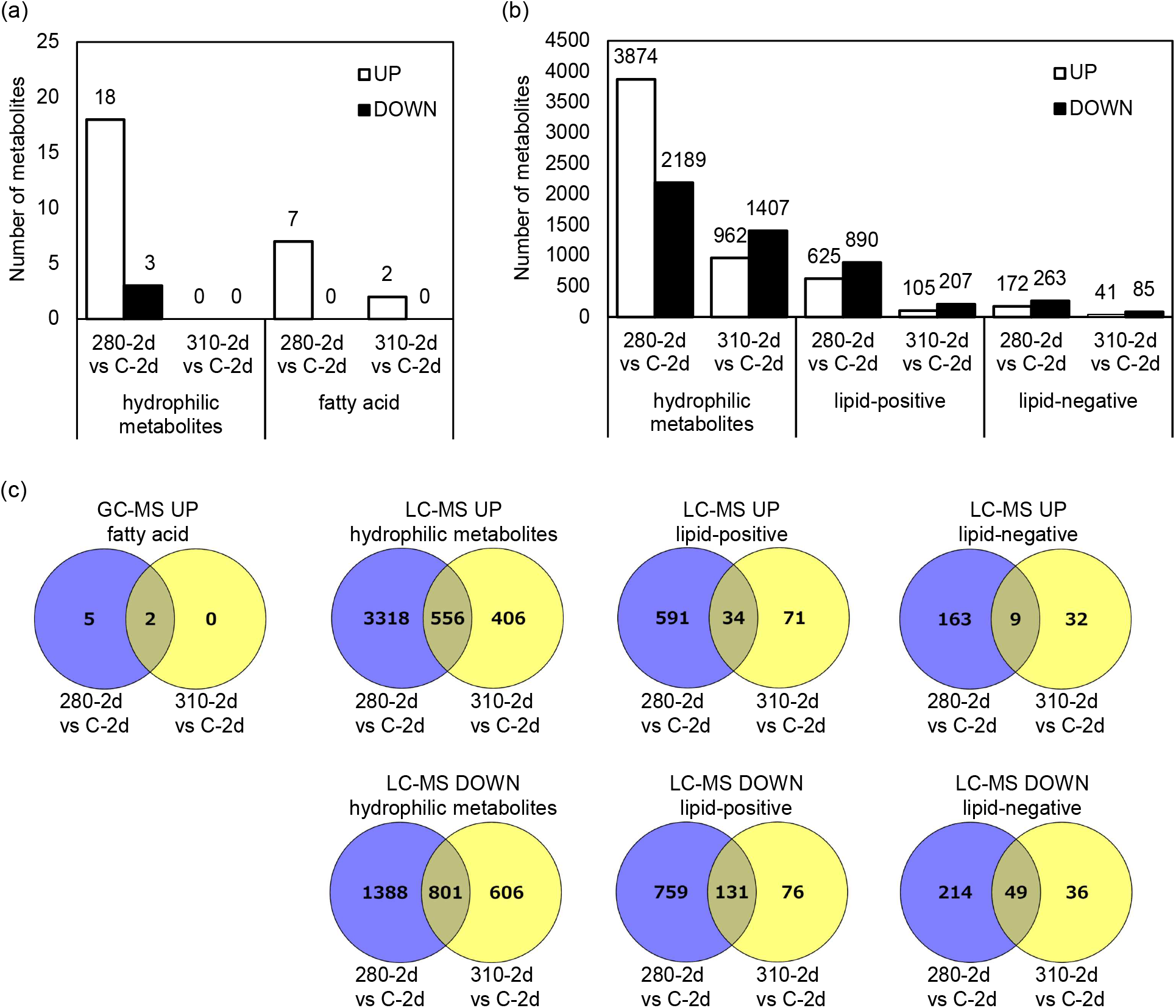
Overall description of metabolome data (a) Comparison of increased and decreased numbers of metabolites under 280 and 310 nm UV annotated by GC-MS. Welch’s *t*-test was applied for comparison of UV-irradiated samples with control, and a metabolite with a *P*-value less than 0.05 was considered significantly changed. (b) Comparison of increased and decreased numbers of metabolites under 280 and 310 nm UV annotated by LC-MS. Welch’s *t*-test was applied for comparison of UV-irradiated samples with control, and metabolite with a *P*-value less than 0.05 was considered significantly changed. lipid-positive, lipids annotated in the positive ion mode; lipid-negative, lipids annotated in the negative ion mode. (c) Venn diagrams illustrating the shared and unique increased and decreased metabolites between samples under 280 and 310 nm UV-B.

In LC-MS lipid analysis, positive and negative ion measurements were performed at the same time. Since some peaks appeared only in either ion mode, the positive and negative ions were separately analyzed. Increased numbers of hydrophilic metabolites were 3874 and 962, those of lipids in the positive ion mode (lipid-positive) were 625 and 105, and those of lipids in the negative ion mode (lipid-negative) were 172 and 41, in 280-2d and 310-2d, respectively. Decreased numbers of hydrophilic metabolites were 2189 and 1407, those of lipid-positive were 890 and 207, and those of lipid-negative were 263 and 85, in 280-2d and 310-2d, respectively (Figure 3b). Again, it was shown that changes in metabolic profiles under 280 and 310 nm UV-LED irradiation were not fully overlapped. The numbers of specifically increased hydrophilic metabolites were 3318 and 406, those of lipid-positive were 591 and 71, and those of lipid-negative were 163 and 32, in 280-2d and 310-2d, respectively, and numbers of specifically decreased hydrophilic metabolites were 1388 and 606, those of lipid-positive were 759 and 76, and those of lipid-negative were 214 and 36, in 280-2d and 310-2d, respectively (Figure 3c).

Among them, annotated metabolites registered in the metabolic pathway in Arabidopsis in KEGG are shown in the metabolic map in Figure 4. For example, metabolites in the TCA cycle and fatty acid biosynthesis, such as dodecanoic acid, decanoic acid, and octanoic acid, were increased more than two-fold in 280-2d in comparison with the control. On the other hand, metabolites in glucosinolate biosynthesis, such as glucoiberverin and glucoerucin, and glutathione metabolism, γ-L-glutamyl-L-cysteine, were decreased more than two-fold in 280-2d. Metabolites in glucosinolate biosynthesis, glucobrassicin, and phenylpropanoid biosynthesis, ferulate, were decreased more than two-fold in 310-2d. Identified lipids were further classified according to LIPID MAPS (Liebisch *et al*., 2020) and the method described in Murphy (2015). It was shown that, in 280-2d, 17 lipids in ceramide (Cer) were characteristically increased, whereas 14 in phosphatidylethanolamine (PE), and 19 in triglyceride (TG) were decreased.

**Figure 4.**
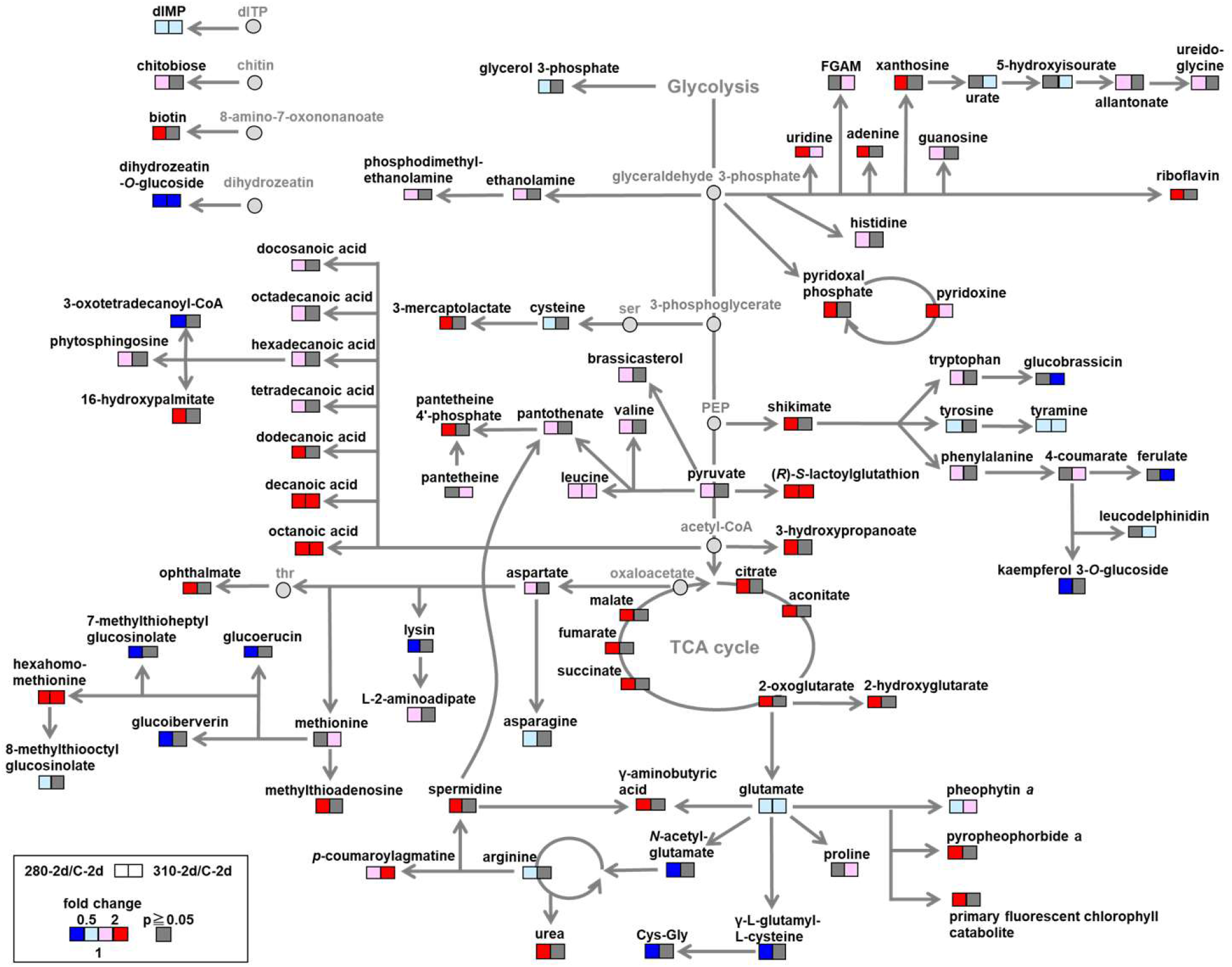
Metabolic responses in Arabidopsis to 280 and 310 nm UV-LED irradiation. Welch’s *t*-test was applied for comparison of UV-irradiated samples with control, and metabolite with a *P*-value less than 0.05 was considered significantly changed. The annotated metabolites are shown in squares and the degrees of changes are visualized by the colors: red, increased by UV-LED irradiation; blue, decreased by UV-LED irradiation; gray, not significantly changed (*P* ≥ 0.05).

### 280 nm specific activation of lipid, polyamine metabolism and TCA cycle

Since the expression of genes related to lipid metabolism (Figure 2c) and the amounts of lipids were specifically increased or decreased under 280 nm UV-LED irradiation (Figure 4 and Table 1), the expression of genes involved in lipid metabolism was extracted from the transcriptomic data. Metabolome analysis showed that, in 280-2d, lipids in Cer were increased, whereas those in PE were decreased (Table 1). Cer is synthesized by Cer synthases catalyzing the condensation of sphingoid bases and coenayme A (CoA)-activated fatty acids (Figure 5a). We assumed that the decreases of lipids in PE were due to their degradation. PE is hydrolyzed by phospholipase D (PLD) (Figure 5b) (Janda *et al*. 2013, Qin and Wang, 2002). Hence, gene expression of Cer synthases and PLDs was focused on. In addition, the relationship between the Cer metabolism and the hypersensitive response (HR)-type PCD has been shown (Ternes *et al*. 2011; Luttgeharm *et al*. 2015). Therefore, expression of HR-type PCD genes was also focused on. Gene expression of a member of Cer synthases, *LOH2* (Figure 5c), and four PLDs, *PLPζ2*, *PLDβ2*, *PLDγ3*, and *PLDγ1* (Figure 5d), were significantly increased in 280-2d together with five PCD markers, *FLAVIN-DEPENDENT MONO-OXYGENASE1* (*FMO1*), *peroxidase C* (*PRXc*), *SENESCENCE-ASSOCIATED GENE 13* (*SAG13*), and *pathogenesis-related 2* and *3* (*PR2*, *PR3*) (Figure 5e). This gene expression profile corresponds well with the metabolome data.

**Figure 5.**
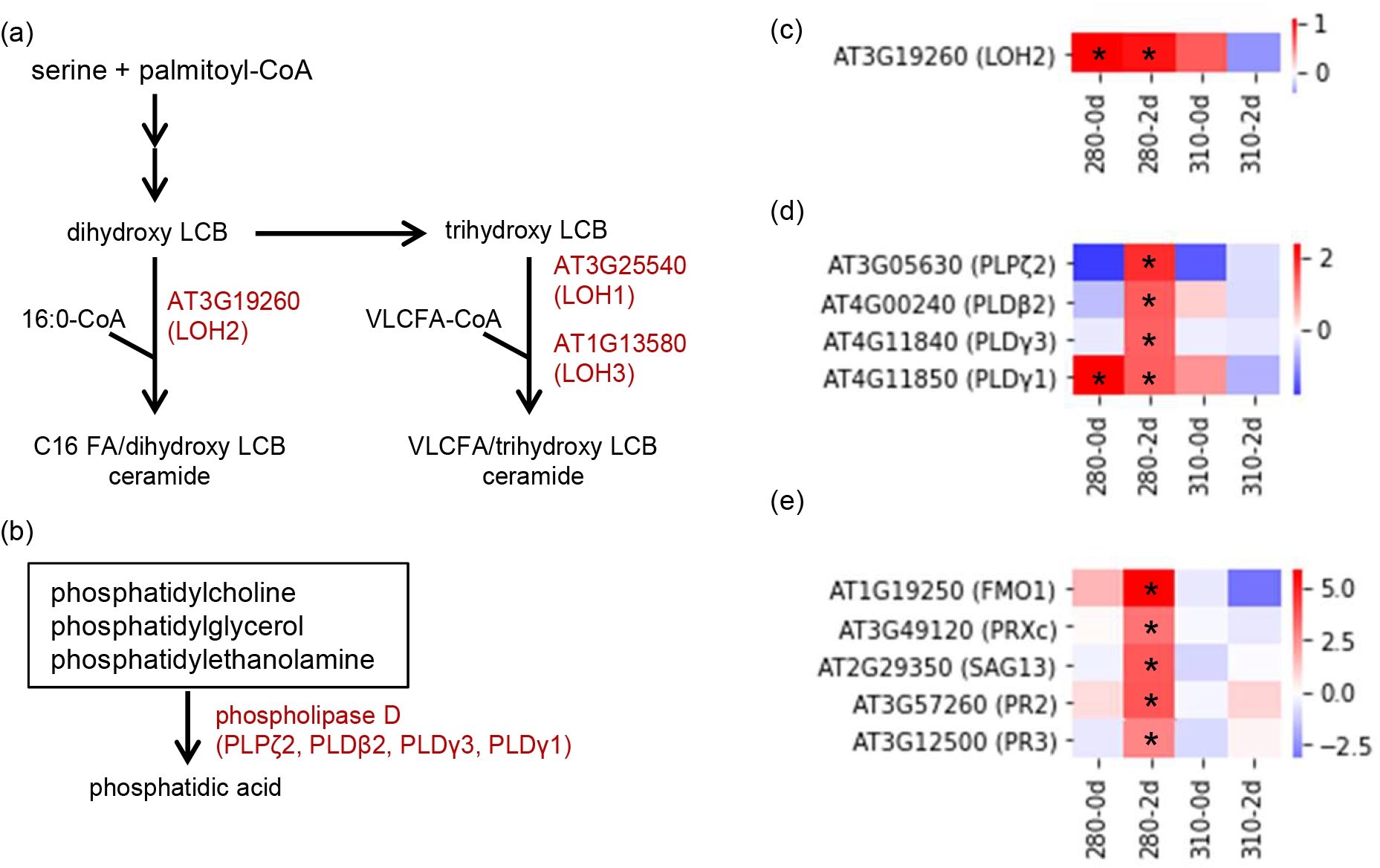
Gene expression profiles of Cer synthase, PLDs, and markers in HR-type PCD under 280 and 310 nm UV-LEDs (a) Metabolic map of Cer biosynthesis. LCB, long-chain base; FA, Fatty acid, VLC, very-long-chain. (b) Metabolic map of PE hydrolysis. (c) Heat map showing the expression level of *LOH2*. (d) Heat map showing the expression level of the *PLDs*. (e) Heat map showing the expression level of HR-type PCD marker genes. Changes in gene expression levels relative to the control are expressed as log_2_ (fold change) values. As shown on the color scale, blue indicates down-regulation and red indicates up-regulation. Asterisks indicate significant differences between the control and UV-LED irradiation using Welch’s *t*-test (\**P* < 0.05).

**Table 1.** Classification of identified lipids whose amounts were significantly changed by UV-LED irradiation. Welch’s *t*-test was performed to determine significant differences (*P* < 0.05) between the control and UV-LED irradiated Arabidopsis.

| Group | Lipid class | Increased |  |  | Decreased |  |  |
| --- | --- | --- | --- | --- | --- | --- | --- |
|  |  | 280-2d<br>specific | 310-2d<br>specific | common | 280-2d<br>specific | 310-2d<br>specific | common |
| Neutral lipids | AcHexChE | 1 | - | - | - | - | - |
|  | AcHexCmE | 6 | - | - | - | - | - |
|  | AcHexSiE | 2 | - | - | - | - | - |
|  | AcHexZyE | - | - | - | 1 | - | - |
|  | CmE | - | - | - | 2 | - | - |
|  | DG | 2 | - | - | - | - | - |
|  | TG | 13 | 1 | 1 | 19 | 4 | 7 |
| Phospholipids | PA | - | - | - | - | - | 1 |
|  | PC | - | - | - | 1 | - | - |
|  | PE | 4 | - | - | 14 | - | 1 |
|  | PG | 3 | 1 | - | 3 | - | - |
|  | PI | 2 | - | - | 3 | - | - |
|  | PIP2 | - | - | - | - | - | 1 |
|  | PS | 2 | 1 | - | 9 | - | - |
| Sphingolipids | Cer | 17 | - | 1 | 3 | - | - |
|  | Hex1Cer | 2 | - | - | 4 | - | - |
| Derivatized lipids | BisMePA | - | - | - | 1 | 2 | 2 |
| Glycoglycerolipid | MGDG | - | - | - | 3 | - | - |
| Fatty acyl and<br>other lipids | Co | 1 | - | - | - | - | - |

AcHexChE, AcylGlcCholesterol ester; AcHexCmE, AcylGlcCampesterol ester; AcHexSiE, AcylGlcSitosterol ester; AcHexZyE, AcylGlcZymosterol ester; CmE, campesterol ester; DG, diglyceride; TG, triglyceride; PA, phosphatidic acid; PC, phosphatidylcholine; PE, phosphatidylethanolamine; PG, phosphatidylglycerol; PI, phosphatidylinositol; PIP2, phosphatidylinositol; PS, phosphatidylserine; Cer, Ceramides; Hex1Cer, simple Glc series; BisMePA, bis-methyl phosphatidic acid; MGDG, monogalactosyldiacylglycerol; Co, coenzyme

Spermidine (Spd) and γ-aminobutyric acid (GABA), which are produced from polyamine biosynthesis and catabolism, respectively, were also specifically increased in 280-2d. (Figure 4 and Table S3). In accordance with the metabolic change, genes involved in polyamine metabolism, polyamine oxidases, *PAO2*, *PAO3*, and *PAO4*, spermidine synthase, *SPDS3*, aldehyde dehyrogenases, *ALDH2B7* and *ALDH3H1*, and glutamic acid decarboxylase, *GAD1*, were up-regulated in 280-2d (Figure 6). Similarly, up-regulation of most of the genes involved in the TCA cycle, citrate synthase *CSY4*, isocitrate dehydrogenase, *IDH6*, the E_1_ subunit of 2-oxogulatarate dehydrogenase complex, *E1-OGDH1*, the E_2_ subunits of 2-oxogulatarate (α-ketoglutarate) dehydrogenase complex, *KGDHE2* and *At4g26910*, dihydrolipoyl dehydrogenase, *LPD2*, succinyl CoA ligase α-subunit, *At5g08300*, succinate dehydroganases, *SDH1-1*, *SDH2-1*, and *SDH2-2*, fumarase, *FUM2*, and mitochondrial malate dehydrogenase, *mMDH2*, in 280-2d were confirmed (Figure 7).

**Figure 6.**
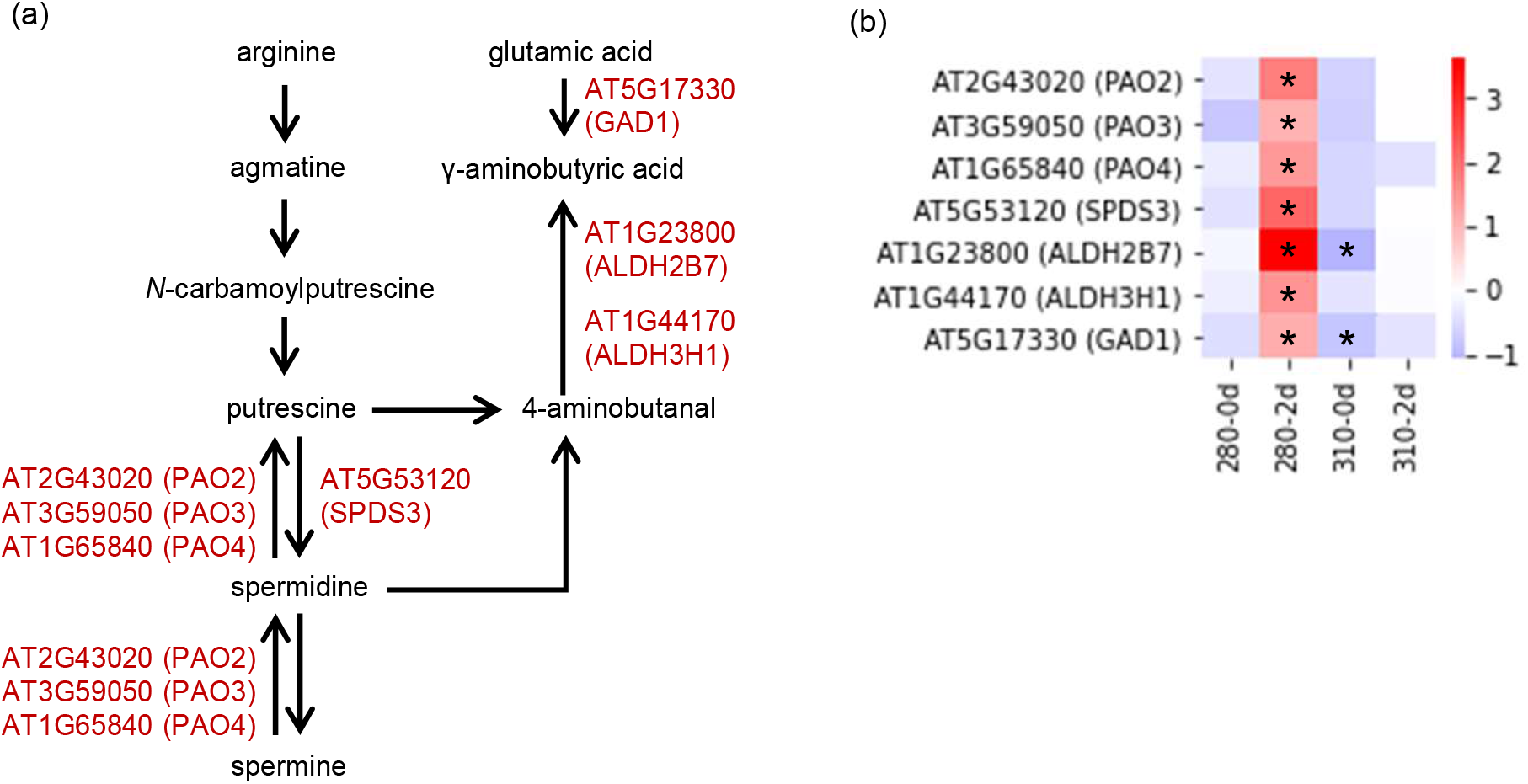
Gene expression profiles of enzymes in polyamine metabolism and GABA biosynthesis under 280 and 310 nm UV-LEDs (a) Metabolic map of polyamine metabolism and GABA biosynthesis. (b) Heat map showing the expression level of enzymatic genes in polyamine metabolism and GABA biosynthesis. Changes in gene expression levels relative to the control are expressed as log_2_ (fold change) values. As shown on the color scale, blue indicates down-regulation and red indicates up-regulation. Asterisks indicate significant differences between the control and UV-LED irradiation using Welch’s *t*-test (\**P* < 0.05).

**Figure 7.**
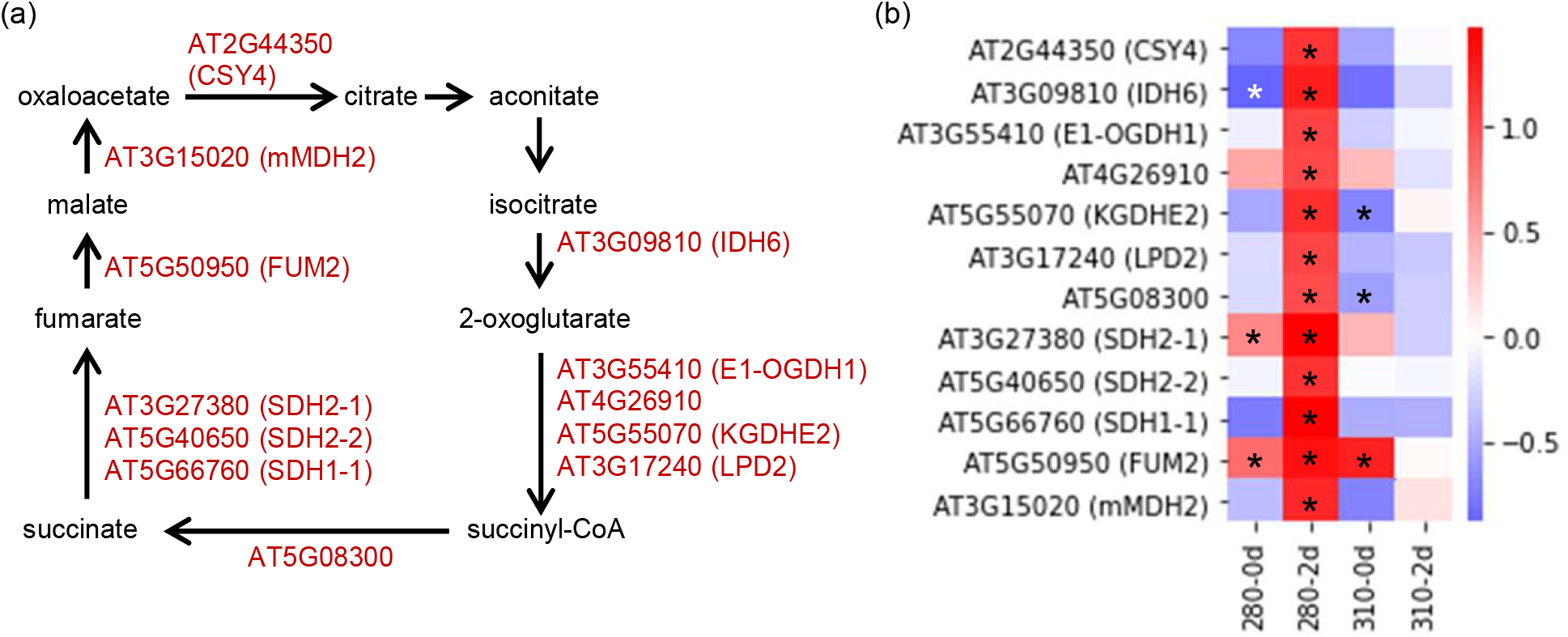
Gene expression profiles of enzymes in TCA cycle under 280 and 310 nm UV-LEDs (a) Metabolic map of TCA cycle. (b) Heat map showing the expression level of enzymatic genes in TCA cycle. Changes in gene expression levels relative to the control are expressed as log_2_ (fold change) values. As shown on the color scale, blue indicates down-regulation and red indicates up-regulation. Asterisks indicate significant differences between the control and UV-LED irradiation using Welch’s *t*-test (\**P* < 0.05).

### Effect of narrowband UV-B on flavonoid biosynthesis

The amounts of shikimic acid and phenylalanine, precursors of flavonoids, were specifically increased in 280-2d (Figure 4). However, contrary to previous studies using BB UV-B lamps, flavonoids were not increased under NB UV-B. Moreover, ferulic acid, leucodelphinidin, and kaempferol 3-glucoside were decreased under NB UV-B instead of increasing (Figure 4). Since a GO term ‘phenylpropanoid biosynthesis’ was enriched in DEGs in 280-2d, we focused on the gene expression of typical enzymes involved in flavonoid biosynthesis, *ADT6*, *PAL1*, *C4H*, *CHS*, *CHI1*, *F3H*, *DFRA*, *LDOX* and *UGT75C1*, which is one of AGTs (Figure 8a,b). We also focused on the gene expression of transcription factors such as HY5, which is a key component in the UVR8 signaling pathway, HYH, a homologue of HY5, MEDs, and members in MBW complexes (MYBs, EGL3, GL3, TTG1, and TT8), which are involved in flavonoid biosynthesis (Figure 8c). Unexpectedly, the expression of genes involved in flavonoid biosynthesis was down-regulated immediately after NB UV-B irradiation (Figure 8b). Then, expression of *ADT6*, *PAL1*, and *C4H*, involved in upstream phenylpropanoid biosynthesis, was specifically increased in 280-2d, whereas expression of *DFR, LDOX*, and *UGT75C1*, involved in downstream anthocyanin biosynthesis was specifically decreased in 310-2d (Figure 8b). A significant increase in gene expression of transcription factors, *HY5*, *ATMYBL2* and *TTG1*, and a decrease of *MED33A* and *MYB4* were observed in both 280-0d and 310-0d (Figure 8c). The expression patterns of other transcription factors, except for *MYB11* and *MYB111*, also showed similar tendencies in both 280-0d and 310-0d. However, after two days incubation in the dark, expression of *MYB13, GL3, MYB4, MYB7* were significantly up-regulated, and *MYBL2* expression was down-regulated only in 280-2d. On the other hand, no expression difference was confirmed in 310-2d as compared with the control C-2d.

**Figure 8.**
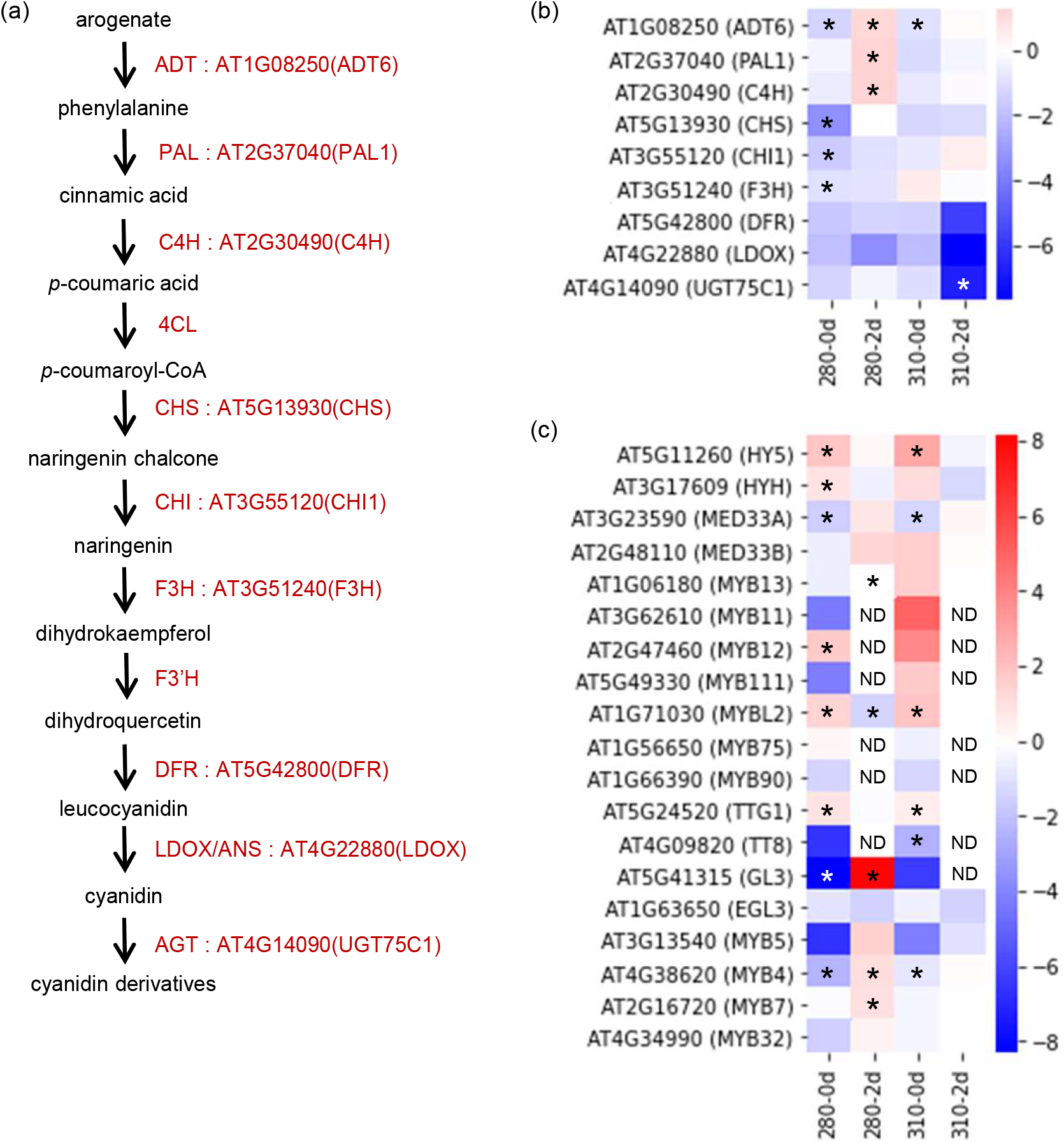
Gene expression profiles of enzymes and transcription factors involved in flavonoid biosynthesis under 280 and 310 nm UV-LEDs (a) Metabolic map of flavonoid GABA biosynthesis. (b) Heat map showing the expression level of enzymatic genes in flavonoid biosynthesis. (c) Heat map showing the expression level of transcription factors in flavonoid biosynthesis. Changes in gene expression levels relative to the control are expressed as log_2_ (fold change) values. As shown on the color scale, blue indicates down-regulation and red indicates up-regulation. Asterisks indicate significant differences between the control and UV-LED irradiation using Welch’s *t*-test (\**P* < 0.05). ND, not detected.

## DISCUSSION

### Effects of 280 and 310 nm UV-LED irradiation on gene expression and metabolic profile are different

Many studies have been conducted on the responses of plants to UV-B, but few studies have been conducted on the effect of each wavelength within the UV-B. Here, it was confirmed that the numbers of DEGs and increased or decreased metabolites were significantly lower under 310 nm UV-LED irradiation than under 280 nm UV-LED irradiation. HY5 regulates the expression of half of UVR8-regulated genes including defense-related and flavonoid biosynthetic ones (Brown *et al*., 2005; Vandenbussche *et al*., 2018). Here, we found that the responses to 280 and 310 nm UV-LED irradiation were inconsistent with each other even though HY5, a key transcription factor in UVR8 signaling, was induced by both UV-LEDs (Figures 2–8). UVR8 is monomerized in the range of 260 to 335 nm with the maximum response at 280 nm (Díaz-Ramos et al., 2018). Under 310 nm irradiation, the monomerization efficiency was lower than under 280 nm irradiation, but *HY5* expression was comparable with that under 280 nm irradiation (Díaz-Ramos *et al*., 2018; Rai *et al*., 2020). Most of the studies on UV-B responses so far used BB UV-B lamps, which have a peak wavelength around 310 nm (Brown *et al*., 2005; Lake *et al*., 2009; Kusano *et al*., 2011; Vandenbussche *et al*., 2018). Since BB UV-B lamps contain a wide range of wavelengths, there is a possibility that the observed phenomena in the previous studies were caused by the interaction of multiple light signaling pathways.

One of the possible reasons for this may be that UVR8 is highly sensitive to 280 nm but less to 310 nm as indicated by *in vitro* monomerization (Díaz-Ramos *et al*., 2018), and the response to 310 nm is mediated by other photoreceptors. For example, in Arabidopsis seeds, phytochrome A (phyA) signaling has been shown to induce germination under a wide range of wavelengths from 300 to 780 nm (Shinomura *et al*., 1996). It is known that phyA also inactivates the COP1/SUPPRESSOR OF PHYA-105 complex, leading to the rapid accumulation of transcription factors such as HY5 (Saijo *et al*., 2003; Sheerin *et al*., 2015). LONG HYPOCOTYL IN FAR-RED1 (HFR1), which acts in phyA signaling, has been shown to be stabilized under UV-B (Fairchild *et al*., 2000; Tavridou *et al*., 2020). In our experiment, only 310 nm UV-LEDs induced the expression of *HFR1* (Table S4). Therefore, phyA signaling may be involved in the induction of *HY5* by 310 nm UV-LED irradiation. It is also possible that other photoreceptors, such as phyB and CRYs, which absorb 310 nm, were activated by 310 nm UV-LED irradiation. Further research using each photoreceptor mutant is required to dissect the different responses under 280 and 310 nm UV-LED irradiation.

### Stress response is induced by 280 nm UV-B

Irradiation of Arabidopsis with UV-B has been reported to induce salicylic acid and jasmonic acid associated and defense-related genes (Surplus *et al*., 1998; A.-H.-Mackerness *et al*., 1999; Vandenbussche *et al.*, 2018). Jasmonic acid is biosynthesized from α-linolenic acid (Vick and Zimmerman, 1984). Here, we confirmed that DEGs in Arabidopsis with 280 nm UV-LED irradiation showed enriched defense responses, including those to biotic and abiotic responses, responses to salicylic acid and jasmonic acid, and α-linolenic acid metabolic pathways. On the other hand, 310 nm UV-LED irradiation did not enrich these terms, but rather down-regulated ‘response to salicylic acid’ (Figure 2 and Table S5). The results indicate that stress responses are induced by short wavelengths in UV-B below 310 nm. Previously, it was shown that UVR8 dependent defense responses against *Botrytis cinerea* were retarded even in jasmonate signaling mutants, *jar1-1* and *P35S:JAZ10.4* (Demkura and Ballaré, 2012). On the other hand, mitogen-activated protein kinase (MAPK) signaling pathway involved in an environmental stress response was shown to be UVR8 independent (González Besteiro *et al*., 2011). Accordingly, the signaling pathway that induced the 280 nm specific stress responses in this study remain to be uncovered.

Irradiation of peach skin with UV-B changed the amounts of certain lipids (Santin *et al*., 2018). NB UV-B LED irradiation of Arabidopsis also changed the amounts of certain lipids, and 280 nm UV-LED irradiation induced a characteristic change of the lipid profile as an increase in Cer and decreases in PE and TG (Table 1). Cer is synthesized by ceramide synthase, LOH1, LOH2, and LOH3 in Arabidopsis (Markham *et al*., 2011). Overexpression of *LOHs* affected Arabidopsis growth, and HR-type PCD marker genes have been shown to be up-regulated in the *LOH2* overexpressing strain of Arabidopsis (Luttgeharm *et al*. 2015). In this study, the expression of *LOH2* and HR type PCD marker genes, *FMO*, *PRXc*, *SAG13*, *PR2*, and *PR3*, were increased by 280 nm-UV LED irradiation (Figure 5c,e). PCD was induced in UV-B treated BY-2 tobacco cells (Lytvyn *et al*., 2010). Our results might explain one of the mechanisms involved in PCD induction through up-regulation of *LOH2* by UV-B. Multiple photoreceptor signaling pathways including UVR8-dependent and -independent pathways have also been proposed for PCD (Petrov *et al*., 2015). Utilization of NB UV-LED in future molecular genetic studies will promote our understanding of mechanisms in PCD induction by UV-B.

Phosphatidylethanolamine is a group of phospholipids, which is the major constituent of cell membranes. Phosphatidylethanolamine is known to be hydrolyzed by phospholipase D (PLD) (Janda *et al*. 2013; Takák, T., *et al*., 2019). Our result showed that the gene expression of enzymes hydrolyzing PE, *PLPζ2*, *PLDβ2*, *PLDγ3*, and *PLDγ1*, were increased by 280 nm UV-LED irradiation (Figure 5d). This might be the reason for the specific decrease in PE by 280 nm UV-LED irradiation. Phosphatidic acid produced by hydrolysis of PE plays an important role in the stress response of plants (Yao and Xue, 2018). Decreased PE and up-regulation of the hydrolyzing enzyme gene are also stress responses specifically induced by 280 nm UV-LED irradiation. The decreases of lipids were also found in TG (Table 1). Triglyceride is used as an energy source through glycolysis and the TCA cycle (Thompson *et al*., 1998). The increase of organic acids in the TCA cycle discussed below and fatty acid may be due to the degradation of TG (Figure 4).

Polyamines and GABA are induced in the responses to various stresses, including UV, and exhibit cytoprotective effects (Rhee *et al*., 2007; Kinnersley and Turano, 2000; Kusano *et al*., 2011; Chen *et al*., 2019; Jiao and Gu, 2019). Here, we also found that 280 nm UV-LED irradiation specifically increased the polyamine, SPD, and GABA (Figure 4 and Table S3). Homeostasis of polyamines is regulated by the dynamic balance of biosynthesis and catabolism. Biosynthesis is catalyzed by arginine decarboxylase (ADC), agmatine iminohydrolase (AIH), *N*-carbamoylputrescine amidohydrolase (CPA), spermidine synthase (SPDS), spermine synthase (SPMS) and thermospermine synthase (ACL5). Catabolism involves two types of enzymes. One is copper-dependent diamine oxidase (DAO) and the other is flavin adenine diamine (FAD)-dependent PAO (Wang *et al*., 2019; Yu *et al*., 2019). As a result of transcriptome analysis, expression of catabolic genes, *PAOs*, was induced by 280 nm UV-LED irradiation (Figure 4b). It has been shown that polyamines of higher molecular weight, including spermine and Spd, are subjected to PAO-mediated catabolism when their levels increase beyond a threshold (Zhang *et al*. 2015). We found that *ADC1* was also up-regulated under 280 nm UV-LED irradiation (data not shown), so there is a possibility that polyamine biosynthesis was activated under 280nm UV-LED irradiation, and the polyamine exceeded the threshold for induction of *PAO* expression. Elucidation of precise mechanisms of Spd increase needs further investigation. 4-Aminobutanal generated by the terminal catabolism of polyamines is then converted to GABA by ALDHs (Zarei *et al*., 2015, 2016). GABA is also synthesized from glutamic acid by glutamate decarboxylase (GAD) (Baum *et al*., 1996; Kinnersley and Turano, 2000). As a result of transcriptome analysis, both GAD and ALDH were increased (Figure 6b). Accordingly, it is considered that 280 nm UV-B is a signal to increase GABA production, however, the involvement of UVR8 signaling in the regulation of polyamine and GABA metabolism remains to be elucidated.

### Metabolites in the TCA cycle increase by 280 nm UV-B

It was shown that UV-B irradiation increased the amounts of metabolites in the first half of the TCA cycle and expression of most of the enzymatic genes in the TCA cycle (Kusano *et al.*, 2011). Regarding metabolites, not only citrate, aconitate, and 2-oxoglutarate in the first half of the TCA cycle, but also succinate, fumarate, and malate in the second half increased under 280 nm UV-LED irradiation (Figure 4). As well as the metabolome, the transcriptome in 280-2d showed specific changes (Figure 7b). On the other hand, our results showed that 310 nm UV-LED irradiation did not affect the metabolites in the TCA cycle. The difference in the effects of 280 and 310 nm on the central metabolism was also confirmed using NB UV-B LEDs.

### Narrowband UV-B irradiation does not induce flavonoid biosynthesis

UV-B stimulates the expression of genes encoding enzymes involved in the anthocyanin biosynthetic pathway in Arabidopsis (Tohge *et al.*, 2011). UV-B is sensed by the photoreceptor UVR8, which activates the transcription factor HY5; then, HY5 up-regulates the expression of the transcription factors that control the biosynthesis of flavonoids and anthocyanins (Brown *et al*., 2005; Stracke *et al*., 2010; Tohge *et al.*, 2011; Shin *et al*., 2013). In addition, it has been shown that UVR8 binds directly to the transcription factor MYB13 regulating flavonoid biosynthesis in a UV-B-dependent manner (Qian *et al.*, 2020). Among the members in MBW complexes, at least *MYB12*, *MYB111*, *MYB75*, *MYB13* and *GL3* have been shown to be induced by UV-B (Brown *et al*., 2005; Stracke *et al*., 2010; Tohge *et al*., 2011; Yan *et al*., 2012; Qian *et al*., 2020). In this study *HY5*, *MYBL2*, and *TTG1* were significantly up-regulated, and *HYH* and *MYB12* also tended to be up-regulated by both 280 and 310 nm UV-LED irradiation. *MYB4*, a repressor of flavonoid biosynthesis, was also commonly down-regulated under irradiation by both UV-LEDs (Figure 8c). However, other transcription factors were not affected by NB UV-LED irradiation. MYBL2 is also a repressor of flavonoid biosynthesis, which suppresses the expression of *F3H*, *DFR*, *LDOX*, *GL3*, *TT8* and *MYB75*, and is negatively regulated by strong light (Dubos *et al*., 2008). The AtGenExpress global stress expression data show that *MYBL2* expression is induced by abiotic stresses including UV-B (Kilian *et al*., 2007; Winter *et al*., 2007). Here, we confirmed the up-regulation of *MYBL2* by NB UV-B irradiation (Figure 8c). The expression of *F3H*, *DFR*, *LDOX*, *GL3*, and *TT8*, showed a decreasing tendency, which is considered to be due to the influence of MYBL2 (Figure 8b,c). MYB4 suppresses the expression of *C4H* (Jin *et al*. 2000). In our experiments, expression of *MYB4* was reduced by both 280 and 310 nm UV-LED irradiation, but expression of *C4H*, together with *ADT6* and *PAL*, was increased only by 280 nm irradiation (Figure 8b,c). There was no significant difference in the expression of *MYB13*, *MYB11* and *MYB111*, but interestingly they showed the opposite behavior under 280 and 310 nm UV-LED irradiation (Figure 8c). After all, NB UV-LED irradiation could induce only upstream phenylpropanoid biosynthesis but not downstream flavonoid and anthocyanin biosynthesis. The phenomena reported in previous studies with BB UV-B lamps may have been induced by multiple signal transductions, as shown by the inconsistent gene expression profiles of members in MBW complexes under 280 and 310 nm UV-LED irradiation in this study.

In conclusion, our study revealed that the responsivenesses of Arabidopsis to 280 and 310 nm UV-B were significantly different, and Arabidopsis distinguished 280 and 310 nm UV-B. It is considered that the phenomena confirmed in the previous experiments using BB UV-B lamps were induced by the multiple signal transductions generated by several wavelengths in UV-B. Our results obtained with NB UV-B LEDs indicate that UV-B signal transduction is mediated through more complex pathways than the current model in which HY5 plays a central role. Utilization of NB UV-LEDs will lead to new insights into the plant UV-B responses.

## EXPERIMENTAL PROCEDURES

### Plant materials and growth conditions

Seeds of *Arabidopsis thaliana* Columbia (Col-0) were sterilized with 0.5% sodium hypochlorite and 0.02% (w/v) Triton-X100 and cultured on half strength Murashige and Skoog medium (pH 6.0) containing 100 mg l^−1^ *myo*-inositol, 0.1 mg l^−1^ thiamin hydrochloride, 0.5 mg l^−1^ nicotinic acid, 0.5 mg l^−1^ pyridoxine, 2 mg l^−1^ glycine, 1% sucrose and 0.8% agar in a growth chamber at 23°C with a 16-h light/8-h dark photoperiod for 14 days.

### UV-B treatment

Fourteen-day-old Arabidopsis seedlings were kept under LED light with a peak wavelength of 280 nm and a half-bandwidth of 10 nm (NCSU234BU280, Nichia, Tokushima, Japan) or LED light with a peak wavelength of 310 nm and a half-bandwidth of 10 nm (NCSU234BU310, Nichia) for 45 min at 2.5 μmol m^−2^ s^−1^, and were kept at 23°C for two days in the dark.

### RNA-Sequencing

RNA extraction and transcriptome analysis were conducted at Takara Bio, Shiga, Japan. Briefly, total RNA was extracted from approximately 100 mg fresh weight (FW) of Arabidopsis shoots using NucleoSpin RNA Plant (Takara Bio) according to the manufacturer’s instructions. Three biological replicates for each treatment were used for analysis. RNA amplification was done using SMART-Seq v4 Ultra Low Input RNA Kit for Sequencing (Illumina, San Diego, CA, USA). A DNA library was prepared using the Nextera XT DNA Sample Preparation Kit (Illumina). RNA-Seq was performed using NovaSeq system (Illumina), and the obtained nucleotide sequences were mapped to the Arabidopsis genome sequence (TAIR 10.46) using STAR version 2.6.0c (Dobin *et al.*, 2013). Then, gene expression levels were estimated as transcripts per million (TPM) using Genedata Profiler Genome version 13.0.11 (Genedata, Basel, Switzerland).

### Identification and enrichment analysis of DEGs

A statistical comparison of TPM was performed using Microsoft Excel to select DEGs between control and UV-LED irradiated samples. The data were presented as means (*n* = 3), and Welch’s *t*-test was applied to detect significant differences. A gene with a *P*-value less than 0.05 and a difference more than two-fold was considered as a DEG. To identify the most significant gene sets associated with GO and KEGG pathways, enrichment analysis of DEGs was performed using DAVID (http://david.abcc.ncifcrf.gov) (Huang et al, 2009a; 2009b).

### Profiling of hydrophilic metabolites using GC-MS

Metabolite extraction and metabolome analysis were conducted at Kazusa DNA Research Institute, Chiba, Japan. Briefly, 50 mg FW of Arabidopsis shoots were extracted with 75–80% methanol, loaded on a MonoSpin C18 column (GL Science, Tokyo, Japan), and eluted with 70% methanol. Methoxyamine and pyridine were added to the obtained fraction for methoximation, and then *N*-methyl-*N*-(trimethylsilyl)trifluoroacetamide was added for trimethylsilylation. Three biological replicates for each treatment were used for analysis. The analysis was performed using a gas chromatograph–quadrupole mass spectrometer, QP2010 Ultra (Shimadzu, Kyoto, Japan), with an auto sampler AOC-5000 Plus (Shimadzu). Chromatographic separation was achieved using a DB-5 column (inner diameter, 0.25 mm × 30 m and film thickness, 1.00 μm, Agilent Technologies, Wilmingston, NC, USA). The carrier gas was helium at a flow rate of 1.1 ml min^−1^. The injection temperature was 280°C, and the injection volume was 0.5 μl. The temperature program was isothermal for 4 min at 100°C, and was then raised at a rate of 4°C min^−1^ to 320°C and held for 8 min. The transfer line temperature, ion source temperature and scan speed were set to 280°C, 200°C and 2000 unit s^−1^, respectively. Data acquisition was performed in the mass range of 45 to 600 *m/z*. The obtained data was analyzed using the GCMSsolution software (Shimazu) and the GC/MS metabolic component database Ver.2 (Shimadzu).

### Profiling of fatty acids using GC-MS

Metabolite extraction and metabolome analysis were conducted at Kazusa DNA Research Institute. Briefly, 50 mg FW of Arabidopsis shoots were extracted with 650 μl of methanol/methyl *tert*-butyl ether, 3:10, 125 μl of ultrapure water was added, and the methyl *tert*-butyl ether fraction was collected. After adding 10% boron trifluoride methanol to the obtained fraction for methyl esterification, ultrapure water and hexane were added, and then a hexane fraction was analyzed. Three biological replicates for each treatment were used for analysis. The analysis was performed using a QP2010 Ultra (Shimadzu) with an auto sampler AOC-5000 Plus (Shimadzu). Chromatographic separation was achieved using a DB-5ms column (inner diameter, 0.25 mm × 30 m and film thickness, 0.25 μm, Agilent Technologies). The carrier gas was helium at a flow rate of 1.1 ml min^−1^. The injection temperature was 280°C, and the injection volume was 0.5 μl. The temperature program was isothermal for 2 min at 40°C, and was then raised at a rate of 6°C min^−1^ to 320°C and held for 5 min. The transfer line temperature, ion source temperature and scan speed were set to 280°C, 200°C and 2500 unit sec^−1^, respectively. Data acquisition was performed in the mass range of 45 to 500 *m/z*. The analysis of the obtained data is same as described above.

### Profiling of hydrophilic metabolites using LC-MS

Metabolite extraction and metabolome analysis were conducted at Kazusa DNA Research Institute. Briefly, 100 mg FW of Arabidopsis shoots were extracted with 75% methanol, loaded on a MonoSpin C18 column (GL Science), and eluted with 75% methanol. Three biological replicates for each treatment were used for analysis. The analysis was performed using a high-performance liquid chromatography (HPLC) Ultimate 3000 RSLC (Thermo Fisher Scientific, Waltham, MA, USA) coupled with a high-resolution mass spectrometer Q Exactive (Thermo Fisher Scientific) with electrospray ionization (ESI) in the positive mode. Chromatographic separation was achieved using an Inert Sustain AQ-C18 column (2.1 mm × 150 mm, 3 μm-particle, GL Science). The column was kept at 40°C, and the flow rate was 0.2 ml min^−1^. The mobile phase solutions were water with 0.1% formic acid (eluent A) and acetonitrile (eluent B) and were implemented in the following gradient: 0–3 min, 2% B; and 3–30 min, 2–98% B. The injection volume was 2 μl. Mass spectrometry conditions were as follows: the scan range was set at *m/z* 80–1200. The full scan resolution was 70,000. The MS/MS scan resolution was 17,500. The obtained data was analyzed using a ProteoWizard (http://proteowizard.sourceforge.net) and a PowerGetBatch (Kazusa DNA Research Inst.). Then, the KEGG database (http://www.genome.jp/kegg/) was used to annotate the metabolites.

### Profiling of lipids using LC-MS

Metabolite extraction and metabolome analysis were conducted at Kazusa DNA Research Institute. Briefly, 100 mg FW of Arabidopsis shoots were extracted with 650 μl of methanol/methyl *tert*-butyl ether, 3:10, 125 μl of ultrapure water was added, and the methyl *tert*-butyl ether fraction was collected. Three biological replicates for each treatment were used for analysis. The analysis was performed using an Ultimate 3000 RSLC (Thermo Fisher Scientific) coupled with a Q Exactive (Thermo Fisher Scientific) with ESI in the positive or negative mode. Chromatographic separation was achieved using a SunShell C18 column (2.1 mm × 150 mm, 2.6 μm-particle, ChromaNik Technologies, Osaka, Japan). The column was kept at 40°C, and the flow rate was 0.2 ml min^−1^. The mobile phase solutions were acetonitrile/water (60:40 v/v) (eluent A) and 2-propanol/acetonitrile (90:10 v/v) (eluent B), both containing 0.1% formic acid and 10 mM ammonium formate and were implemented in the following gradients: 0–10 min, 30–35% B; 10–20 min, 35–55% B; 20–35 min, 55–65% B; 35–45 min, 65–100% B; and 45–50 min, 100% B. The injection volume was 2 μl. Mass spectrometry conditions were the same as described above. The obtained data was analyzed using a ProteoWizard, a PowerGetBatch (Kazusa DNA Research Inst.) and a Lipid Search (Mitsui Knowledge Industry Co., Ltd., Tokyo, Japan). Lipid classification was performed by Lipid Search according to the LIPID MAPS (https://www.lipidmaps.org/, Liebisch *et al*., 2020) and the method of Murphy (2015) described previously (Yamada et al., 2013), and the MS/MS spectrum was compared with the lipid spectrum registered in the Lipid Search database. Some of the detected peaks had the same retention time in the positive and negative modes. In that case, the one with the larger peak area was selected.

### Statistical analysis for metabolite profiling

Differences between the relative quantity of metabolites were evaluated using Welch’s *t*-test. The data were presented as the means (*n* = 3), and *P*-values less than 0.05 were considered statistically significant.

## DATA AVAILABILITY STATEMENT

RNA-seq data were deposited in the DDBJ Sequence Read Archive (https://www.ddbj.nig.ac.jp/dra/index.html), under accession number DRA011512. Metabolomics data were submitted to MetaboLights (https://www.ebi.ac.uk/metabolights/, Haug *et al*., 2013), under study ID MTBLS2461.

## ACKNOWLEDGMENTS

This research is supported in part by the JSBBA Fund for Industry-Academia-Government Collaboration (to A.O.) and Nichia Corporation (to T.T. and Y.F.).

## SUPPORTING INFORMATION

Additional Supporting Information may be found in the online version of this article.

**Table S1**. Common DEGs in 280-0d and 310-0d.

**Table S2**. Common DEGs in 280-2d and 310-2d.

**Table S3**. Enriched terms of KEGG pathway and gene ontology (GO) biological process for sample-specific DEGs

**Table S4**. List of metabolites whose amounts were significantly changed by UV-LED irradiation.

**Table S5**. DEGs specifically up-regulated in 310-0d.

## REFERENCES

A.-H.-Mackerness, S., Surplus, S.L., Blake, P., John, C.F., Buchanan-Wollaston, V., Jordan, B.R. and Thomas, B. (1999) Ultraviolet-B-induced stress and change in gene expression in *Arabidopsis thaliana*: role of signalling pathways controlled by jasmonic acid, ethylene and reactive oxygen species. Plant Cell Environ. 22, 1413–1423.

Baum, G., Lev-Yadun, S., Fridmann, Y., Arazi, T., Katsnelson, H., Zik, M. and Fromm, H. (1996) Calmodulin binding to gluatamate decarboxylase is required for regulation of gluatamate and GABA metabolism and normal development in plants. EMBO J. 15, 2988–2996.

Bonawitz, N.D., Soltau, W.L., Blatchley, M.R., Powers, B.L., Hurlock, A.K., Seals, L.A., Weng, J.-K., Stout, J. and Chapple, C. (2012) REF4 and RFR1, subunits of the transcriptional coregulatory complex mediator, are required for phenylpropanoid homeostasis in *Arabidopsis*. J. Biol. Chem. 287, 5434–5445.

Broun, P. (2005) Transcriptional control of flavonoid biosynthesis: a complex network of conserved regulators involved in multiple aspects of differentiation in *Arabidopsis*. Curr. Opin. Plant Biol. 8, 272–279.

Brown, B.A., Cloix, C., Jiang, G.H., Kaiserli, E., Herzyk, P., Kliebenstein, D.J. and Jenkins, G.I. (2005) A UV-B-specific signaling component orchestrates plant UV protection. Proc. Natl. Acad. Sci. USA, 102, 18225–18230.

Chen, D., Shao, Q, Yin, L., Younis, A. and Zheng, B. (2019) Polyamine function in plants: metabolism, regulation on development, and roles in abiotic stress responses. Front. Plant Sci. 9, 1945.

Demkura, P.V. and Ballaré, C.L. (2012) UVR8 mediates UV-B induced *Arabidopsis* defense responses against *Botrytis cinerea* by controlling sinapate accumulation. Mol. Plant 5, 642–652.

Díaz-Ramos, L.A., O’Hara, A., Kanagarajan, S., Farkas, D., Strid, Å. and Jenkins, G.I. (2018) Difference in the action spectra for UVR8 monomerisation and *HY5* transcript accumulation in Arabidopsis. Photochem. Photobiol. Sci. 17, 1108–1117.

Dobin, A., Davis, C.A., Schlesinger, F., Drenkow, J., Zaleski, C., Jha, S., Batut, P., Chaisson, M. and Gingeras, T.R. (2013) STAR: ultrafast universal RNA-seq aligner. Bioinformatics, 29, 15–21.

Dotto, M. and Casati, P. (2017) Developmental reprogramming by UV-B radiation in plants. Plant Sci. 264, 96–101.

Dubos, C., Le Gourrierec, J., Baudry, A., Huep, G., Lanet, E., Debeaujon, I., Routaboul, J.-M., Alboresi, A., Weisshaar, B. and Lepiniec, L. (2008) MYBL2 is a new regulator of flavonoid biosynthesis in *Arabidopsis thaliana*. Plant J. 55, 940–953.

Fairchild, C.D., Schumaker, M.A. and Quail, P.H. (2000) HFR1 encodes an atypical bHLH protein that acts in phytochrome A signal transduction. Genes Dev. 14, 2377–2391.

Fraser, C.M. and Chapple, C. (2011) The phenylpropanoid pathway in Arabidopsis. Arabidopsis Book, 9, e0152.

Gangappa, S.N. and Botto, J.F. (2016) The multifaceted roles of HY5 in plant growth and development. Mol. Plant, 9, 1353–1365.

Gill, S.S. and Tuteja, N. (2010) Reactive oxygen species and antioxidant machinery in abiotic stress tolerance in crop plants. Plant Physiol. Biochem. 48, 909–930.

González Besteiro, M.A., Bartles, S., Albert, A. and Ulm, R. (2011) Arabidopsis MAP kinase phosphatase 1 and its target MAP kinase 3 and 6 antagonistically determine UV-B stress tolerance, independent of the UVR8 photoreceptor pathway. Plant J. 68, 727–737.

Haug, K., Salek, R.M., Conesa, P., Hastings, J., de Matos, P., Rijnbeek, M., Mahendraker, T., Williams, M., Neumann, S., Rocca-Serra, P., Maguire, E., González-Beltrán, A., Sansone, S.-A., Griffin, J.L. and Steinbeck, C. (2013) MetaboLights—an open-access general-purpose repository for metabolomics studies and associated meta-data. Nucleic Acids Res. 41, D781–D786

Heijde, M. and Ulm, R. (2012) UV-B photoreceptor-mediated signalling in plants. Trends Plant Sci. 17, 230–237.

Hideg, É., Jansen, M.A.K. and Strid, A. (2013) UV-B exposure, ROS, and stress: inseparable companions or loosely linked associates? Trends Plant Sci. 18, 107–115.

Huang, D.W., Sherman, B.T. and Lempicki, R.A. (2009a) Systematic and integrative analysis of large gene lists using DAVID bioinformatics resources. Nat. Protoc. 4, 44–57.

Huang, D.W., Sherman, B.T. and Lempicki, R.A. (2009b) Bioinformatics enrichment tools: paths toward the comprehensive functional analysis of large gene lists. Nucleic Acids Res. 37, 1–13.

Janda, M., Planchais, S., Djafi, N., Martinec, J., Burketova, L., Valentova, O., Zachowski, A. and Ruelland, E. (2013) Phosphoglycerolipids are master players in plant hormone signal transduction. Plant Cell Reports, 32, 839–851.

Jenkins, G.I. (2009) Signal transduction in responses to UV-B radiation. Annu. Rev. Plant Biol. 60, 407–431.

Jenkins, G.I. (2014) The UV-B photoreceptor UVR8: from structure to physiology. Plant Cell 26, 21–37.

Jenkins, G.I. (2017) Photomorphogenic responses to ultraviolet-B light. Plant Cell Environ. 40, 2544–2557.

Jiao, C. and Gu, Z. (2019) iTRAQ-based proteomic analysis reveals changes in response to UV-B treatment in soybean sprouts. Food Chem. 275, 467–473.

Jin, H., Cominelli, E., Bailey, P., Parr, A., Mehrtens, F., Jones, J., Tonelli, C., Weisshaar, B. and Martin, C. (2000) Transcriptional repression by AtMYB4 controls production of UV-protecting sunscreens in *Arabidopsis*. EMBO J. 19, 6150–6161.

Kilian, J., Whitehead, D., Horak, J., Wanke, D., Weinl, S., Batistic, O., D’Angelo, C., Bornberg-Bauer, E., Kudla, J. and Harter, K. (2007) The AtGenExpress global stress expression data set: protocols, evaluation and model data analysis of UV-B light, drought and cold stress responses. Plant J. 50, 347–363.

Kinnersley, A.M. and Turano, F.J. (2000) Gamma aminobutyric acid (GABA) and plant responses to stress. Crit. Rev. Plant Sci. 19, 479–509.

Kusano, M., Tohge, T., Fukushima, A., Kobayashi, M., Hayashi, N., Otsuki, H., Kondou, Y., Goto, H., Kawashima, M., Matsuda, F., Niida, R., Matsui, M., Saito, K. and Fernie, A.R. (2011) Metabolomics reveals comprehensive reprogramming involving two independent metabolic responses of Arabidopsis to UV◻B light. Plant J. 67, 354–369.

Lake, J.A., Field, K.J., Davey, M.P., Beerling, D.J. and Lomax, B.H. (2009) Metabolomic and physiological responses reveal multi-phasic acclimation of *Arabidopsis thaliana* to chronic UV radiation. Plant Cell Environ. 32, 1377–1389.

Liebisch, G., Fahy, E., Aoki, J., Dennis, E.A., Durand, T., Ejsing, C.S., Fedorova, M., Feussner, I., Griffiths, W.J., Köfeler, H., Merrill, Jr. A.H., Murphy, R.C., O’Donnell, V.B., Oskolkova, O., Subramaniam, S., Wakelam, M.J.O. and Spener, F. (2020) Update on LIPID MAPS classification, nomenclature, and shorthand notation for MS-derived lipid structures. J. Lipid Res. 61, 1539–1555.

Luttgeharm, K.D., Chen, M., Mehra, A., Cahoon, R.E., Markham, J.E. and Cahoon, E.B. (2015) Overexpression of Arabidopsis ceramide synthases differentially affects growth, sphingolipid metabolism, programmed cell death, and mycotoxin resistance. Plant Physiol. 169, 1108–1117.

Lytvyn, D.I., Yemets, A.I. and Blume, Y.B. (2010) UV-B overexposure induces programmed cell death in a BY-2 tobacco cell line. Environ. Exp. Bot. 68, 51–57.

Ma, D. and Constable, P. (2019) MYB repressors as regulators of phenylpropanoid metabolism in plants. Trends Plant Sci. 24, 275–289.

Markham, J.E., Molino, D., Gissot, L., Bellec, Y., Hématy, K., Marion, J., Belcram, K., Palauqui, J.-C., Satiat-JeuneMa□tre, B. and Faure, J.-D. (2011) Sphingolipids containing very-long-chain fatty acids define a secretory pathway for specific polar plasma membrane protein targeting in *Arabidopsis*. Plant Cell 23, 2362–2378.

Murphy, R.C. (2015) Tandem Mass Spectrometry of Lipids: Molecular Analysis of Complex Lipids. Royal Society of Chemistry.

Petrov, V., Hille, J., Mueller-Roever, B. and Gechev, T.S. (2015) ROS-mediated abiotic stress-induced programmed cell death in plants. Front. Plant Sci. 6, 69.

Qian, C., Chen, Z., Liu, Q., Mao, W., Chen, Y., Tian, W., Liu, Y., Han, J., Ouyang, X. and Huang, X. (2020) Coordinated transcriptional regulation by the UV-B photoreceptor and multiple transcription factors for plant UV-B responses. Mol. Plant, 13, 777–792.

Qin, C. and Wang, X. (2002) The Arabidopsis phospholipase D family. Characterization of a calcium-independent and phosphatidylcholine-selective PLDζ1 with distinct regulatory domains. Plant Physiol. 128, 1057–1068.

Rai, N., O’Hara, A., Farkas, D., Safronov, O., Ratanasopa, K., Wang, F., Lindfors, A.V., Jenkins, G.I., Lehto, T., Salojärvi, J., Brosché, M., Strid, Å., Aphalo, P.J. and Morales, L.O. (2020) The photoreceptor UVR8 mediates the perception of both UV-B and UV-A wavelengths up to 350 nm of sunlight with responsivity moderated by cryptochromes. Plant Cell Environ. 43, 1513–1527.

Rhee, H.J., Kim, E.-J. and Lee, J.K. (2007) Physiological polyamines: simple primordial stress molecules. J. Cell. Mol. Med. 11, 685–703.

Rizzini, L., Favory, J.-J., Cloix, C., Faggionato, D., O’Hara, A., Kaiserli, E., Baumeister, R., Schäfer, E., Nagy, F., Jenkins, G.I. and Ulm, R. (2011) Perception of UV-B by the *Arabidopsis* UVR8 protein. Science, 332, 103–106.

Rozema, J., van de Staaji, J., Björn, L.O. and Caldwell, M. (1997) UV-B as an environmental factor in plant life: stress and regulation. Trends Ecol. Evol. 12, 22–28.

Saijo, Y., Sullivan, J.A., Wang, H., Yang J., Shen, Y., Rubio V., Ma, L., Hoecker, U. and Deng, X.W. (2003) The COP1–SPA1 interaction defines a critical step in phytochrome A-mediated regulation of HY5 activity. Genes Dev. 17, 2642–2647.

Santin, M., Lucini, L., Castagna, A., Chiodelli, G., Hauser, M.-T. and Ranieri, A. (2018) Post-harvest UV-B radiation modulates metabolite profile in peach fruit. Postharvest Biol. Technol. 139, 127–134.

Sheerin, D.J., Menon, C., zur Oven-Krockhaus, S., Enderle, B., Zhu, L., Johnen, P., Schleifenbaum, F., Stierhof, Y.-D., Huq, E. and Hiltbrunner, A. (2015) Light-activated phytochrome A and B interact with members of the SPA family to promote photomorphogenesis in Arabidopsis by reorganizing the COP1/SPA complex. Plant Cell, 27, 189–201.

Shin, D.H., Choi, M., Kim, K. Bang, G., Cho, M., Choi, S.-B., Choi, G. and Park, Y.-I. (2013) HY5 regulates anthocyanin biosynthesis by inducing the transcriptional activation of the MYB75/PAP1 transcription factor in *Arabidopsis*. FEBS Lett. 587, 1543–1547.

Shinomura, T., Nagatani, A. Hanzawa, H., Kubota, M., Watanabe, M. and Furuya, M. (1996) Action spectra for phytochrome A- and B-specific photoinduction of seed germination in *Arabidopsis thaliana*. Proc. Natl. Acad. Sci. USA 93, 8129–8133.

Stracke, R., Favory, J.-J., Gruber, H., Bartelniewoehner, L., Bartels, S., Binkert, M., Funk, M., Weisshaar, B. and Ulm, R. (2010) The *Arabidopsis* bZIP transcription factor HY5 regulates expression of the *PFG1/MYB12* gene in response to light and ultraviolet-B radiation. Plant Cell Environ. 33, 88–103.

Surplus, S.L., Jordan, B.R., Murphy, A.M., Carr, J.P., Thomas, B. and A.-H.-Mackerness, S. (1998) Ultraviolet-B-induced responses in *Arabidopsis thaliana*: role of salicylic acid and reactive oxygen species in the regulation of transcripts encoding photosynthetic and acidic pathogenesis-related proteins. Plant Cell Environ. 21, 685–694.

Takáč, T., Novák, D. and Šamaj, J. (2019) Recent advances in the cellular and developmental biology of phospholipases in plants. Front. Plant Sci. 10, 362.

Tavridou, E., Schmid-Siegert, E., Fankhauser, C. and Ulm, R. (2020) UVR8-mediated inhibition of shade avoidance involves HFR1 stabilization in Arabidopsis. PLoS Genet. 16, e1008797.

Ternes, P., Feussner, K., Werner, S., Lerche, J., Iven, T., Heilmann, I., Riezman, H. and Feussner, I. (2011) Disruption of the ceramide synthase LOH1 causes spontaneous cell death in *Arabidopsis thaliana*. New Phytol. 192, 841–854.

Thompson, J.E., Froese, C.D., Madey, E., Smith, M.D. and Hong, Y. (1998) Lipid metabolism during plant senescence. Prog. Lipid Res., 37, 119–141.

Tohge, T., Kusano, M., Fukushima, A., Saito, K. and Fernie, A. R. (2011) Transcriptional and metabolic programs following exposure of plants to UV-B irradiation. Plant Signal. Behav. 6, 1987–1992.

Vandenbussche, F., Yu, N., Li, W., Vanhaelewyn, L., Hamshou, M., Van Der Straeten, D. and Smagghe, G. (2018) An ultraviolet B condition that affects growth and defense in *Arabidopsis*. Plant Sci. 268, 54–63.

Vick, B.A. and Zimmerman, D.C. (1984) Biosynthesis of jasmonic acid by several plant species. Plant Physiol. 75, 458–461.

Wang, W., Paschalidis, K., Feng, J.-C., Song, J. and Liu, J.-H. (2019) Polyamine catabolism in plants: a universal process with diverse functions. Front. Plant Sci. 10, 561.

Wang, X.-C., Wu, J., Guan, M.-L., Zhao, C.-H., Geng, P. and Zhao, Q. (2020) *Arabidopsis* MYB4 plays dual roles in flavonoid biosynthesis. Plant J. 101, 637–652.

Winter, D., Vinegar, B., Nahal, H., Ammar, R., Wilson, G.V. and Provart, N.J. (2007) An “electronic Fluorescent Pictograph” browser for exploring and analyzing large-scale biological data sets. PLoS One, 8, e718.

Xu, W., Grain, D., Bobet, S., Gourrierec, J.L., Thévenin, J., Kelemen, Z., Lepiniec, L. and Dubos, C. (2014) Complexity and robustness of the flavonoid transcriptional regulatory network revealed by comprehensive analyses of MYB-bHLH-WDR complexes and their targets in Arabidopsis seed. New Phytol. 202, 132–144.

Xu, W., Dubos, C. and Lepiniec L. (2015) Transcriptional control of flavonoid biosynthesis by MYB-bHLH-WDR complexes. Trends Plant Sci. 20, 176–185.

Yamada, T., Uchikata, T., Sakamoto, S., Yokoi, Y., Fukusaki, E. and Bamba, T. (2013) Development of a lipid profiling system using reverse-phase liquid chromatography coupled to high-resolution mass spectrometry with rapid polarity switching and an automated lipid identification software. J. Chormatogr. A, 1292, 211–218.

Yan, A., Pan, J., An, L., Gan, Y. and Feng, H. (2012) The responses of trichome mutants to enhanced ultraviolet-B radiation in *Arabidopsis thaliana*. J. Photochem. Photobiol. B Biol. 113, 29–35.

Yao, H.-Y. and Xue, H.-W. (2018) Phosphatidic acid plays key roles regulating plant development and stress responses. J. Integr. Plant Biol. 60, 851–863.

Yin, R. and Ulm, R. (2017) How plants cope with UV-B: from perception to response. Curr. Opin. Plant Biol. 37, 42–48.

Yu, Z., Jia, D. and Liu, T. (2019) Polyamine oxidases play various roles in plant development and abiotic stress tolerance. Plants, 8, 184.

Zarei, A., Trobacher, C.P. and Shelp, B.J. (2015) NAD^+^-aminoaldehyde dehydrogenase candidates for 4-aminobutyrate (GABA) and β-alanine production during terminal oxidation of polyamines in apple fruit. FEBS Lett. 589, 2695–2700.

Zarei, A., Trobacher, C.P. and Shelp, B.J. (2016) Arabidopsis aldehyde dehydrogenase 10 family members confer salt tolerance through putrescine-derived 4-aminobutyrate (GABA) production. Sci. Rep. 6, 35115.

Zhang, Q., Wang, M., Hu, J., Wang, W., Fu, X. and Liu, J.-H. (2015) PtrABF of *Poncirus trifoliata* functions in dehydration tolerance by reducing stomatal density and maintaining reactive oxygen species homeostasis. J. Exp. Bot. 66, 5911–5927.

