## Supplemental Table S1 for "Transcriptome and metabolome analyses revealed that narrowband 280 and 310 nm UV-B induce distinctive responses in Arabidopsis"

Common DEGs in 280-0d and 310-0d

| ID | Gene name | Log2 FC | | *P* -value | | Description |
|---|---|---|---|---|---|---|
| | | 280-0d | 310-0d | 280-0d | 310-0d | |
| **'280-0d and 310-0d'_DEGs up-regulated** | | | | | | |
| AT1G01570 | - | 1.45 | 1.41 | 0.044 | 0.044 | F22L4.11 protein [Source:UniProtKB/TrEMBL;Acc:Q9LMM4] |
| AT1G02360 | - | 2.94 | 3.69 | 0.034 | 0.011 | Chitinase family protein [Source:TAIR;Acc:AT1G02360] |
| AT1G03610 | - | 1.32 | 1.66 | 0.002 | 0.001 | Plant/protein (DUF789) [Source:UniProtKB/TrEMBL;Acc:Q8LF98] |
| AT1G05675 | UGT74E1 | 6.56 | 3.80 | 0.020 | 0.036 | Glycosyltransferase (Fragment) [Source:UniProtKB/TrEMBL;Acc:A0A1W6AJW6] |
| AT1G05805 | BHLH128 | 1.52 | 1.99 | 0.010 | 0.005 | Transcription factor bHLH128 [Source:UniProtKB/Swiss-Prot;Acc:Q8H102] |
| AT1G07128 | - | 3.70 | 2.38 | 0.013 | 0.028 | Potential natural antisense gene, locus overlaps with AT1G07130 [Source:TAIR;Acc:AT1G07128] |
| AT1G07130 | STN1 | 3.28 | 1.85 | 0.000 | 0.002 | CST complex subunit STN1 [Source:UniProtKB/Swiss-Prot;Acc:Q9LMK5] |
| AT1G07150 | MAPKKK13 | 2.65 | 3.75 | 0.007 | 0.010 | F10K1.14 protein [Source:UniProtKB/TrEMBL;Acc:Q9LMK8] |
| AT1G07240 | UGT71C5 | 1.44 | 1.48 | 0.009 | 0.019 | UDP-glycosyltransferase 71C5 [Source:UniProtKB/Swiss-Prot;Acc:Q9FE68] |
| AT1G08500 | ENODL18 | 2.50 | 1.96 | 0.013 | 0.047 | Early nodulin-like protein 18 [Source:UniProtKB/TrEMBL;Acc:O82083] |
| AT1G08550 | VDE1 | 1.22 | 1.51 | 0.010 | 0.002 | NPQ1 [Source:UniProtKB/TrEMBL;Acc:A0A384K8V4] |
| AT1G08930 | ERD6 | 3.33 | 2.31 | 0.001 | 0.019 | ERD6 [Source:UniProtKB/TrEMBL;Acc:A0A178W9Q9] |
| AT1G09250 | BHLH149 | 1.53 | 2.32 | 0.001 | 0.003 | Transcription factor bHLH149 [Source:UniProtKB/Swiss-Prot;Acc:O80482] |
| AT1G09940 | HEMA2 | 2.68 | 1.02 | 0.001 | 0.050 | Glutamyl-tRNA reductase 2, chloroplastic [Source:UniProtKB/Swiss-Prot;Acc:P49294] |
| AT1G10770 | - | 11.91 | 11.20 | 0.001 | 0.008 | At1g10770 [Source:UniProtKB/TrEMBL;Acc:Q9SAC5] |
| AT1G11080 | scpl31 | 1.66 | 1.39 | 0.031 | 0.010 | Carboxypeptidase [Source:UniProtKB/TrEMBL;Acc:F4I7C9] |
| AT1G12090 | ELP | 1.71 | 2.15 | 0.002 | 0.001 | ELP [Source:UniProtKB/TrEMBL;Acc:A0A178W1A4] |
| AT1G13670 | - | 1.44 | 2.62 | 0.005 | 0.008 | At1g13670 [Source:UniProtKB/TrEMBL;Acc:Q9LMY0] |
| AT1G13930 | - | 2.32 | 2.55 | 0.000 | 0.007 | At1g13930/F16A14.27 [Source:UniProtKB/TrEMBL;Acc:Q9XI93] |
| AT1G15800 | - | 2.24 | 2.71 | 0.012 | 0.008 | At1g15800 [Source:UniProtKB/TrEMBL;Acc:Q6NL17] |
| AT1G15890 | - | 2.20 | 1.63 | 0.000 | 0.007 | Probable disease resistance protein At1g15890 [Source:UniProtKB/Swiss-Prot;Acc:Q9LMP6] |
| AT1G16110 | WAKL6 | 2.47 | 1.76 | 0.002 | 0.040 | Wall-associated receptor kinase-like 6 [Source:UniProtKB/Swiss-Prot;Acc:Q8GXQ3] |
| AT1G17030 | - | 3.45 | 2.84 | 0.030 | 0.046 | Uncharacterized protein At1g17030/F20D23_27 [Source:UniProtKB/TrEMBL;Acc:Q8GWU4] |
| AT1G17420 | LOX3 | 2.68 | 1.39 | 0.004 | 0.033 | Lipoxygenase 3, chloroplastic [Source:UniProtKB/Swiss-Prot;Acc:Q9LNR3] |
| AT1G18265 | - | 2.17 | 2.05 | 0.008 | 0.000 | Protein FLOURY 1-like [Source:UniProtKB/Swiss-Prot;Acc:Q9LM24] |
| AT1G18330 | RVE7 | 2.51 | 3.91 | 0.001 | 0.000 | Protein REVEILLE 7 [Source:UniProtKB/Swiss-Prot;Acc:B3H5A8] |
| AT1G18390 | - | 2.77 | 2.01 | 0.021 | 0.043 | Protein kinase superfamily protein [Source:TAIR;Acc:AT1G18390] |
| AT1G19320 | - | 11.05 | 10.90 | 0.031 | 0.031 | At1g19320 [Source:UniProtKB/TrEMBL;Acc:Q9LN66] |
| AT1G20030 | - | 3.14 | 2.46 | 0.003 | 0.026 | Pathogenesis-related thaumatin superfamily protein [Source:UniProtKB/TrEMBL;Acc:Q9LNT0] |
| AT1G21100 | IGMT1 | 3.37 | 3.02 | 0.000 | 0.047 | Indole glucosinolate O-methyltransferase 1 [Source:UniProtKB/Swiss-Prot;Acc:Q9LPU5] |
| AT1G21110 | IGMT3 | 5.88 | 3.29 | 0.014 | 0.035 | Indole glucosinolate O-methyltransferase 3 [Source:UniProtKB/Swiss-Prot;Acc:Q9LPU6] |
| AT1G21120 | - | 6.43 | 3.11 | 0.000 | 0.027 | O-methyltransferase family protein [Source:TAIR;Acc:AT1G21120] |

| | | | | | | |
|---|---|---|---|---|---|---|
| AT1G21550 | CML44 | 4.77 | 1.77 | 0.000 | 0.011 | Probable calcium-binding protein CML44 [Source:UniProtKB/Swiss-Prot;Acc:Q9LPK5] |
| AT1G22570 | NPF5.15 | 3.18 | 3.62 | 0.012 | 0.005 | Protein NRT1/ PTR FAMILY 5.15 [Source:UniProtKB/Swiss-Prot;Acc:Q9SK99] |
| AT1G22770 | GI | 1.88 | 2.64 | 0.016 | 0.006 | Protein GIGANTEA [Source:UniProtKB/Swiss-Prot;Acc:Q9SQI2] |
| AT1G23550 | SRO2 | 1.99 | 1.48 | 0.024 | 0.046 | Probable inactive poly [ADP-ribose] polymerase SRO2 [Source:UniProtKB/Swiss-Prot;Acc:Q9ZUD9] |
| AT1G24147 | - | 4.23 | 2.09 | 0.000 | 0.001 | Transmembrane protein [Source:UniProtKB/TrEMBL;Acc:Q1G3U1] |
| AT1G25400 | - | 3.43 | 1.16 | 0.000 | 0.021 | At1g25400 [Source:UniProtKB/TrEMBL;Acc:Q9C6L0] |
| AT1G27930 | - | 1.09 | 1.31 | 0.000 | 0.001 | Probable methyltransferase At1g27930 [Source:UniProtKB/Swiss-Prot;Acc:Q9C7F9] |
| AT1G30190 | - | 4.48 | 1.79 | 0.000 | 0.019 | Cotton fiber protein [Source:UniProtKB/TrEMBL;Acc:Q9C6Z9] |
| AT1G30720 | - | 2.32 | 2.15 | 0.043 | 0.017 | Berberine bridge enzyme-like 10 [Source:UniProtKB/Swiss-Prot;Acc:Q9SA87] |
| AT1G33590 | - | 2.55 | 1.21 | 0.000 | 0.041 | Leucine-rich repeat (LRR) family protein [Source:TAIR;Acc:AT1G33590] |
| AT1G33600 | - | 3.69 | 1.87 | 0.000 | 0.011 | Leucine-rich repeat (LRR) family protein [Source:UniProtKB/TrEMBL;Acc:Q9FW48] |
| AT1G33610 | - | 2.42 | 1.73 | 0.005 | 0.005 | Leucine-rich repeat (LRR) family protein [Source:UniProtKB/TrEMBL;Acc:F4HR91] |
| AT1G34780 | APRL4 | 1.00 | 1.06 | 0.004 | 0.009 | 5'-adenylylsulfate reductase-like 4 [Source:UniProtKB/Swiss-Prot;Acc:Q9SA00] |
| AT1G53430 | - | 1.88 | 1.27 | 0.002 | 0.015 | Probable LRR receptor-like serine/threonine-protein kinase At1g53430 [Source:UniProtKB/Swiss-Prot;Acc:C0LGG8] |
| AT1G53870 | - | 1.06 | 1.32 | 0.027 | 0.001 | Protein LURP-one-related 3 [Source:UniProtKB/Swiss-Prot;Acc:Q67XV7] |
| AT1G53890 | - | 1.03 | 1.32 | 0.024 | 0.001 | Protein of unknown function (DUF567) [Source:TAIR;Acc:AT1G53890] |
| AT1G56250 | VBF | 16.13 | 8.49 | 0.000 | 0.036 | F-box protein VBF [Source:UniProtKB/Swiss-Prot;Acc:Q9C7K0] |
| AT1G56510 | ADR2 | 2.38 | 1.12 | 0.000 | 0.004 | Disease resistance protein ADR2 [Source:UniProtKB/Swiss-Prot;Acc:Q9C7X0] |
| AT1G59850 | TOR1L5 | 2.65 | 2.16 | 0.002 | 0.001 | TORTIFOLIA1-like protein 5 [Source:UniProtKB/Swiss-Prot;Acc:Q9XIE4] |
| AT1G62300 | WRKY6 | 1.20 | 1.07 | 0.016 | 0.023 | Uncharacterized protein At1g62300 (Fragment) [Source:UniProtKB/TrEMBL;Acc:C0SV11] |
| AT1G62510 | - | 1.27 | 1.59 | 0.013 | 0.009 | Bifunctional inhibitor/lipid-transfer protein/seed storage 2S albumin superfamily protein [Source:UniProtKB/TrEMBL;Acc:Q9SXE6] |
| AT1G63740 | - | 1.25 | 1.05 | 0.011 | 0.007 | Disease resistance protein (TIR-NBS-LRR class) family [Source:UniProtKB/TrEMBL;Acc:Q9CAD9] |
| AT1G65390 | PP2A5 | 5.25 | 2.14 | 0.007 | 0.046 | Protein PHLOEM PROTEIN 2-LIKE A5 [Source:UniProtKB/Swiss-Prot;Acc:Q9C5Q9] |
| AT1G65800 | SD16 | 1.03 | 1.03 | 0.004 | 0.037 | Receptor-like serine/threonine-protein kinase SD1-6 [Source:UniProtKB/Swiss-Prot;Acc:Q9S972] |
| AT1G66540 | - | 1.41 | 2.22 | 0.018 | 0.012 | At1g66540 [Source:UniProtKB/TrEMBL;Acc:A2RVN3] |
| AT1G66725 | MIR163 | 4.37 | 3.58 | 0.000 | 0.000 | MIR163; miRNA [Source:TAIR;Acc:AT1G66725] |
| AT1G69640 | SBH1 | 1.20 | 1.24 | 0.004 | 0.005 | Sphinganine C4-monooxygenase 1 [Source:UniProtKB/Swiss-Prot;Acc:Q8VYI1] |
| AT1G69830 | AMY3 | 2.50 | 2.44 | 0.006 | 0.004 | Alpha-amylase 3, chloroplastic [Source:UniProtKB/Swiss-Prot;Acc:Q94A41] |
| AT1G69890 | - | 4.20 | 2.08 | 0.000 | 0.036 | Actin cross-linking protein (DUF569) [Source:UniProtKB/TrEMBL;Acc:Q9CAS2] |
| AT1G70820 | - | 3.71 | 3.73 | 0.001 | 0.000 | At1g70820 [Source:UniProtKB/TrEMBL;Acc:Q9SSL0] |
| AT1G71030 | ATMYBL2 | 1.40 | 1.95 | 0.018 | 0.006 | At1g71030/F23N20_2 [Source:UniProtKB/TrEMBL;Acc:Q9C9A5] |
| AT1G71697 | CK1 | 1.60 | 1.03 | 0.001 | 0.017 | Probable choline kinase 1 [Source:UniProtKB/Swiss-Prot;Acc:Q9M9H6] |
| AT1G72060 | - | 2.40 | 1.58 | 0.006 | 0.033 | Serine-type endopeptidase inhibitor [Source:UniProtKB/TrEMBL;Acc:Q9C7G9] |
| AT1G72940 | - | 3.82 | 1.59 | 0.000 | 0.002 | At1g72940/F3N23_14 [Source:UniProtKB/TrEMBL;Acc:Q9SSN2] |
| AT1G73480 | - | 1.19 | 1.84 | 0.021 | 0.017 | Alpha/beta-Hydrolases superfamily protein [Source:UniProtKB/TrEMBL;Acc:Q94AM5] |
| AT1G73750 | - | 1.76 | 2.13 | 0.012 | 0.002 | Uncharacterised conserved protein UCP031088, alpha/beta hydrolase [Source:TAIR;Acc:AT1G73750] |
| AT1G75380 | BBD1 | 1.09 | 1.42 | 0.011 | 0.003 | Bifunctional nuclease 1 [Source:UniProtKB/Swiss-Prot;Acc:Q9FWS6] |
| AT1G75810 | - | 1.81 | 1.61 | 0.002 | 0.002 | At1g75810 [Source:UniProtKB/TrEMBL;Acc:Q9LQT3] |

| | | | | | | |
|---|---|---|---|---|---|---|
| AT1G76590 | - | 1.81 | 2.35 | 0.002 | 0.044 | At1g76590 [Source:UniProtKB/TrEMBL;Acc:Q2HIW3] |
| AT1G76650 | CML38 | 5.51 | 1.17 | 0.000 | 0.008 | CML38 [Source:UniProtKB/TrEMBL;Acc:A0A178WMC5] |
| AT1G76955 | - | 2.36 | 1.36 | 0.009 | 0.006 | At1g76955 [Source:UniProtKB/TrEMBL;Acc:Q0V819] |
| AT1G78460 | - | 1.79 | 1.76 | 0.000 | 0.015 | AT1G78460 protein [Source:UniProtKB/TrEMBL;Acc:Q9SYN5] |
| AT1G78600 | LZF1 | 3.94 | 4.61 | 0.000 | 0.000 | Light-regulated zinc finger protein 1 [Source:UniProtKB/TrEMBL;Acc:F4IBS4] |
| AT1G80610 | - | 1.27 | 1.07 | 0.006 | 0.002 | At1g80610/T21F11_6 [Source:UniProtKB/TrEMBL;Acc:Q8VYI7] |
| AT1G80920 | ATJ8 | 1.10 | 1.41 | 0.024 | 0.006 | Chaperone protein dnaJ 8, chloroplastic [Source:UniProtKB/Swiss-Prot;Acc:Q9SAG8] |
| AT2G03310 | - | 1.49 | 1.76 | 0.021 | 0.020 | unknown protein; Ha. [Source:TAIR;Acc:AT2G03310] |
| AT2G03440 | NRP1 | 1.44 | 1.16 | 0.001 | 0.000 | Nodulin-related protein 1 [Source:UniProtKB/Swiss-Prot;Acc:Q9ZQ80] |
| AT2G03530 | UPS2 | 2.69 | 1.39 | 0.004 | 0.008 | ureide permease 2 [Source:TAIR;Acc:AT2G03530] |
| AT2G08650 | - | 2.29 | 1.85 | 0.009 | 0.018 | - |
| AT2G15020 | - | 1.14 | 2.04 | 0.026 | 0.026 | At2g15020 [Source:UniProtKB/TrEMBL;Acc:Q9ZUK9] |
| AT2G15042 | - | 2.10 | 2.33 | 0.002 | 0.048 | Leucine-rich repeat (LRR) family protein [Source:TAIR;Acc:AT2G15042] |
| AT2G15880 | PEX3 | 3.95 | 4.37 | 0.000 | 0.018 | Pollen-specific leucine-rich repeat extensin-like protein 3 [Source:UniProtKB/Swiss-Prot;Acc:Q9XIL9] |
| AT2G15890 | MEE14 | 3.55 | 3.80 | 0.000 | 0.022 | CCG-binding protein 1 [Source:UniProtKB/Swiss-Prot;Acc:Q9XIM0] |
| AT2G15970 | COR413PM1 | 1.13 | 1.42 | 0.003 | 0.030 | WCOR413-like protein [Source:UniProtKB/TrEMBL;Acc:A0A178VY40] |
| AT2G17780 | MCA2 | 1.23 | 1.04 | 0.004 | 0.002 | Protein MID1-COMPLEMENTING ACTIVITY 2 [Source:UniProtKB/Swiss-Prot;Acc:Q3EBY6] |
| AT2G18350 | ZHD6 | 2.10 | 1.47 | 0.001 | 0.043 | Zinc-finger homeodomain protein 6 [Source:UniProtKB/Swiss-Prot;Acc:Q9ZPW7] |
| AT2G18969 | - | 1.51 | 1.73 | 0.031 | 0.016 | BEST Arabidopsis thaliana protein match is: sequence-specific DNA binding transcription factors;transcription regulators (TAIR:AT4G30180.1); Ha. [Source:TAIR;Acc:AT2G18969] |
| AT2G22830 | SQE2 | 2.26 | 1.90 | 0.020 | 0.037 | Squalene epoxidase 2, mitochondrial [Source:UniProtKB/Swiss-Prot;Acc:O81000] |
| AT2G22980 | SCPL13 | 1.19 | 1.69 | 0.019 | 0.026 | Serine carboxypeptidase-like 13 [Source:UniProtKB/TrEMBL;Acc:A8MQS0] |
| AT2G27690 | CYP94C1 | 1.51 | 1.29 | 0.042 | 0.020 | Cytochrome P450 94C1 [Source:UniProtKB/Swiss-Prot;Acc:Q9ZUX1] |
| AT2G29170 | - | 2.91 | 2.21 | 0.000 | 0.000 | NAD(P)-binding Rossmann-fold superfamily protein [Source:TAIR;Acc:AT2G29170] |
| AT2G29310 | - | 2.05 | 1.96 | 0.002 | 0.008 | Tropinone reductase homolog At2g29310 [Source:UniProtKB/Swiss-Prot;Acc:Q9ZW14] |
| AT2G29670 | - | 1.82 | 2.30 | 0.002 | 0.000 | At2g29670/T27A16.23 [Source:UniProtKB/TrEMBL;Acc:O82388] |
| AT2G30040 | MAPKKK14 | 1.42 | 2.66 | 0.001 | 0.038 | Mitogen-activated protein kinase kinase kinase 14 [Source:UniProtKB/TrEMBL;Acc:O64741] |
| AT2G30600 | - | 1.02 | 2.13 | 0.032 | 0.002 | BTB/POZ domain-containing protein [Source:UniProtKB/TrEMBL;Acc:F4INV5] |
| AT2G32390 | GLR3.5 | 2.03 | 2.57 | 0.023 | 0.008 | Glutamate receptor 3.5 [Source:UniProtKB/Swiss-Prot;Acc:Q9SW97] |
| AT2G34430 | LHB1B1 | 1.59 | 1.49 | 0.001 | 0.031 | Chlorophyll a-b binding protein, chloroplastic [Source:UniProtKB/TrEMBL;Acc:Q39142] |
| AT2G35290 | - | 3.00 | 1.90 | 0.016 | 0.044 | At2g35290 [Source:UniProtKB/TrEMBL;Acc:Q8GWV0] |
| AT2G35650 | CSLA7 | 1.27 | 1.82 | 0.000 | 0.003 | Glucomannan 4-beta-mannosyltransferase 7 [Source:UniProtKB/Swiss-Prot;Acc:Q9ZQN8] |
| AT2G36750 | UGT73C1 | 3.13 | 2.94 | 0.007 | 0.035 | UDP-glycosyltransferase 73C1 [Source:UniProtKB/Swiss-Prot;Acc:Q9ZQ99] |
| AT2G39660 | BIK1 | 2.92 | 1.26 | 0.008 | 0.035 | BIK1 [Source:UniProtKB/TrEMBL;Acc:A0A178VYW0] |
| AT2G40000 | HSPRO2 | 2.18 | 1.35 | 0.001 | 0.010 | Nematode resistance protein-like HSPRO2 [Source:UniProtKB/Swiss-Prot;Acc:O04203] |
| AT2G40840 | DPE2 | 1.57 | 1.41 | 0.003 | 0.005 | 4-alpha-glucanotransferase DPE2 [Source:UniProtKB/Swiss-Prot;Acc:Q8RXD9] |
| AT2G41100 | CML12 | 3.89 | 2.05 | 0.000 | 0.041 | Calmodulin-like protein 12 [Source:UniProtKB/Swiss-Prot;Acc:P25071] |
| AT2G42540 | COR15A | 3.24 | 2.40 | 0.001 | 0.012 | Protein COLD-REGULATED 15A, chloroplastic [Source:UniProtKB/Swiss-Prot;Acc:Q42512] |
| AT2G45560 | CYP76C1 | 1.80 | 1.85 | 0.005 | 0.000 | CYP76C1 [Source:UniProtKB/TrEMBL;Acc:A0A178VLJ2] |

| | | | | | | |
|---|---|---|---|---|---|---|
| AT2G46170 | RTNLB5 | 1.54 | 1.31 | 0.004 | 0.005 | Reticulon-like protein B5 [Source:UniProtKB/Swiss-Prot;Acc:O82352] |
| AT2G46670 | - | 1.34 | 2.20 | 0.031 | 0.003 | CCT motif family protein (Fragment) [Source:UniProtKB/TrEMBL;Acc:C0SV91] |
| AT2G46790 | APRR9 | 1.35 | 2.29 | 0.004 | 0.005 | Two-component response regulator-like APRR9 [Source:UniProtKB/Swiss-Prot;Acc:Q8L500] |
| AT2G47270 | UPB1 | 3.11 | 1.66 | 0.001 | 0.005 | UPB1 [Source:UniProtKB/TrEMBL;Acc:A0A178VV95] |
| AT2G47890 | COL13 | 2.59 | 2.69 | 0.004 | 0.018 | Zinc finger protein CONSTANS-LIKE 13 [Source:UniProtKB/Swiss-Prot;Acc:O82256] |
| AT3G01205 | - | 1.24 | 1.89 | 0.020 | 0.009 | - |
| AT3G01310 | - | 1.40 | 1.64 | 0.001 | 0.046 | Phosphoglycerate mutase-like family protein [Source:UniProtKB/TrEMBL;Acc:F4J8C7] |
| AT3G05800 | BHLH150 | 1.31 | 2.56 | 0.026 | 0.004 | Transcription factor bHLH150 [Source:UniProtKB/Swiss-Prot;Acc:Q9M9L6] |
| AT3G07195 | - | 4.14 | 2.16 | 0.004 | 0.013 | RPM1-interacting protein 4 (RIN4) family protein [Source:UniProtKB/TrEMBL;Acc:A0A1I9LSZ8] |
| AT3G10020 | - | 2.45 | 2.01 | 0.009 | 0.033 | AT3g10020/T22K18_16 [Source:UniProtKB/TrEMBL;Acc:Q9SR67] |
| AT3G10113 | RVE7L | 2.65 | 4.00 | 0.001 | 0.000 | Protein REVEILLE 7-like [Source:UniProtKB/Swiss-Prot;Acc:F4J2J6] |
| AT3G11580 | - | 1.45 | 1.39 | 0.002 | 0.003 | AP2/B3-like transcriptional factor family protein [Source:UniProtKB/TrEMBL;Acc:A0A1I9LT60] |
| AT3G13061 | - | 1.63 | 1.18 | 0.007 | 0.024 | other RNA [Source:TAIR;Acc:AT3G13061] |
| AT3G15540 | IAA19 | 1.17 | 1.03 | 0.025 | 0.023 | Auxin-responsive protein [Source:UniProtKB/TrEMBL;Acc:Q2VWA2] |
| AT3G15760 | - | 2.69 | 1.69 | 0.001 | 0.002 | At3g15760 [Source:UniProtKB/TrEMBL;Acc:Q9LW03] |
| AT3G15840 | PIFI | 1.50 | 1.41 | 0.001 | 0.039 | Post-illumination chlorophyll fluorescence increase [Source:UniProtKB/TrEMBL;Acc:Q9LVZ5] |
| AT3G20600 | NDR1 | 2.80 | 1.01 | 0.002 | 0.033 | Protein NDR1 [Source:UniProtKB/Swiss-Prot;Acc:O48915] |
| AT3G21150 | BBX32 | 4.13 | 3.21 | 0.001 | 0.046 | B-box zinc finger protein 32 [Source:UniProtKB/Swiss-Prot;Acc:Q9LJB7] |
| AT3G21890 | MIP1B | 2.70 | 3.76 | 0.001 | 0.024 | BBX31 [Source:UniProtKB/TrEMBL;Acc:A0A178VEW7] |
| AT3G22060 | CRRSP38 | 2.44 | 1.63 | 0.000 | 0.013 | Cysteine-rich repeat secretory protein 38 [Source:UniProtKB/Swiss-Prot;Acc:Q9LRJ9] |
| AT3G22070 | - | 1.61 | 1.13 | 0.040 | 0.036 | Proline-rich family protein [Source:UniProtKB/TrEMBL;Acc:Q9LRJ8] |
| AT3G22370 | AOX1A | 1.74 | 1.04 | 0.011 | 0.040 | Ubiquinol oxidase 1a, mitochondrial [Source:UniProtKB/Swiss-Prot;Acc:Q39219] |
| AT3G22961 | - | 2.02 | 1.58 | 0.012 | 0.033 | Paired amphipathic helix (PAH2) superfamily protein [Source:UniProtKB/TrEMBL;Acc:Q9LIJ9] |
| AT3G23030 | IAA2 | 1.09 | 1.21 | 0.002 | 0.020 | Auxin-responsive protein [Source:UniProtKB/TrEMBL;Acc:A0A1I9LQ54] |
| AT3G23170 | - | 2.71 | 1.63 | 0.004 | 0.006 | At3g23170 [Source:UniProtKB/TrEMBL;Acc:Q9LTD3] |
| AT3G25770 | AOC2 | 1.71 | 1.23 | 0.002 | 0.010 | AOC2 [Source:UniProtKB/TrEMBL;Acc:A0A178VKE4] |
| AT3G26180 | CYP71B20 | 1.18 | 1.11 | 0.003 | 0.010 | Cytochrome P450 71B20 [Source:UniProtKB/Swiss-Prot;Acc:Q9LTM3] |
| AT3G26580 | - | 1.20 | 1.20 | 0.012 | 0.048 | Orf03 protein [Source:UniProtKB/TrEMBL;Acc:Q38955] |
| AT3G26740 | CCL | 2.49 | 2.59 | 0.000 | 0.024 | Light-regulated protein 1, chloroplastic [Source:UniProtKB/Swiss-Prot;Acc:Q96500] |
| AT3G29320 | PHS1 | 1.97 | 1.91 | 0.003 | 0.001 | Alpha-glucan phosphorylase 1 [Source:UniProtKB/Swiss-Prot;Acc:Q9LIB2] |
| AT3G44400 | - | 3.34 | 1.52 | 0.001 | 0.010 | Disease resistance protein (TIR-NBS-LRR class) family [Source:UniProtKB/TrEMBL;Acc:Q9M285] |
| AT3G44450 | BIC2 | 3.90 | 3.80 | 0.040 | 0.033 | Protein BIC2 [Source:UniProtKB/Swiss-Prot;Acc:Q9M280] |
| AT3G45860 | CRK4 | 2.50 | 1.97 | 0.018 | 0.027 | Cysteine-rich receptor-like protein kinase 4 [Source:UniProtKB/Swiss-Prot;Acc:Q9LZU4] |
| AT3G46970 | PHS2 | 1.92 | 1.42 | 0.012 | 0.022 | Alpha-glucan phosphorylase 2, cytosolic [Source:UniProtKB/Swiss-Prot;Acc:Q9SD76] |
| AT3G47340 | ASN1 | 1.99 | 2.96 | 0.016 | 0.008 | DIN6 [Source:UniProtKB/TrEMBL;Acc:A0A178VBT4] |
| AT3G47820 | PUB39 | 3.72 | 2.84 | 0.010 | 0.020 | RING-type E3 ubiquitin transferase [Source:UniProtKB/TrEMBL;Acc:A0A178V8W3] |
| AT3G47830 | - | 2.51 | 2.07 | 0.015 | 0.030 | Putative DNA glycosylase At3g47830 [Source:UniProtKB/Swiss-Prot;Acc:F4JCQ3] |
| AT3G48640 | - | 6.21 | 1.42 | 0.002 | 0.040 | Transmembrane protein [Source:UniProtKB/TrEMBL;Acc:Q9SMN5] |

| | | | | | | |
|---|---|---|---|---|---|---|
| AT3G49780 | PSK3 | 1.80 | 1.36 | 0.005 | 0.005 | Phytosulfokines 3 [Source:UniProtKB/Swiss-Prot;Acc:Q9M2Y0] |
| AT3G50950 | RPP13L4 | 1.71 | 1.03 | 0.004 | 0.006 | Disease resistance RPP13-like protein 4 [Source:UniProtKB/Swiss-Prot;Acc:Q38834] |
| AT3G51330 | - | 1.02 | 1.03 | 0.009 | 0.038 | Eukaryotic aspartyl protease family protein [Source:UniProtKB/TrEMBL;Acc:Q84WU7] |
| AT3G51450 | SSL7 | 2.19 | 1.21 | 0.001 | 0.008 | Protein STRICTOSIDINE SYNTHASE-LIKE 7 [Source:UniProtKB/Swiss-Prot;Acc:Q9SD04] |
| AT3G52450 | PUB22 | 1.31 | 1.56 | 0.014 | 0.009 | RING-type E3 ubiquitin transferase [Source:UniProtKB/TrEMBL;Acc:A0A178VML4] |
| AT3G52740 | BIC1 | 1.69 | 2.23 | 0.002 | 0.043 | Protein BIC1 [Source:UniProtKB/Swiss-Prot;Acc:Q9LXJ1] |
| AT3G52748 | - | 3.12 | 1.79 | 0.003 | 0.033 | other RNA [Source:TAIR;Acc:AT3G52748] |
| AT3G53232 | RTFL1 | 2.45 | 2.32 | 0.045 | 0.041 | At3g53232 [Source:UniProtKB/TrEMBL;Acc:Q8LBB5] |
| AT3G53800 | Fes1B | 4.12 | 3.76 | 0.001 | 0.000 | Fes1B [Source:UniProtKB/TrEMBL;Acc:Q9M346] |
| AT3G54065 | - | 3.45 | 3.01 | 0.020 | 0.043 | LOW protein: ankyrin repeat protein [Source:UniProtKB/TrEMBL;Acc:A0A1I9LQJ5] |
| AT3G56090 | FER3 | 1.06 | 1.47 | 0.007 | 0.001 | Ferritin-3, chloroplastic [Source:UniProtKB/Swiss-Prot;Acc:Q9LYN2] |
| AT3G57700 | - | 12.35 | 11.15 | 0.003 | 0.008 | Protein kinase superfamily protein [Source:UniProtKB/TrEMBL;Acc:Q9SVY4] |
| AT3G60200 | - | 2.06 | 1.60 | 0.009 | 0.009 | Uncharacterized protein At3g60200 [Source:UniProtKB/TrEMBL;Acc:Q9M1C3] |
| AT3G61060 | AtPP2-A13 | 1.65 | 2.33 | 0.029 | 0.021 | Phloem protein 2-A13 [Source:UniProtKB/TrEMBL;Acc:F4JD33] |
| AT3G61280 | - | 2.15 | 1.49 | 0.008 | 0.008 | Glycosyltransferase [Source:UniProtKB/TrEMBL;Acc:Q5Q0C1] |
| AT3G61580 | SLD1 | 1.08 | 1.52 | 0.002 | 0.000 | Delta(8)-fatty-acid desaturase 1 [Source:UniProtKB/Swiss-Prot;Acc:Q9ZRP7] |
| AT3G62550 | - | 2.68 | 2.55 | 0.008 | 0.033 | Adenine nucleotide alpha hydrolases-like superfamily protein [Source:UniProtKB/TrEMBL;Acc:Q93W91] |
| AT4G04330 | RBCX1 | 4.03 | 3.86 | 0.000 | 0.013 | Chaperonin-like RBCX protein 1, chloroplastic [Source:UniProtKB/Swiss-Prot;Acc:Q94AU9] |
| AT4G05150 | - | 1.45 | 1.48 | 0.000 | 0.050 | AT4g05150/C17L7_70 [Source:UniProtKB/TrEMBL;Acc:Q940N7] |
| AT4G05330 | AGD13 | 2.24 | 2.30 | 0.022 | 0.014 | AGD13 [Source:UniProtKB/TrEMBL;Acc:A0A178V602] |
| AT4G08785 (2 | - | 2.77 | 2.54 | 0.012 | 0.004 | - |
| AT4G11000 | - | 4.29 | 2.59 | 0.004 | 0.039 | Ankyrin repeat family protein [Source:TAIR;Acc:AT4G11000] |
| AT4G11360 | RHA1B | 1.92 | 1.18 | 0.002 | 0.004 | E3 ubiquitin-protein ligase RHA1B [Source:UniProtKB/Swiss-Prot;Acc:Q9SUS5] |
| AT4G11521 | CRK34 | 3.50 | 2.02 | 0.005 | 0.013 | Putative cysteine-rich receptor-like protein kinase 34 [Source:UniProtKB/Swiss-Prot;Acc:Q8LPI0] |
| AT4G11600 | GPX6 | 1.51 | 1.22 | 0.014 | 0.038 | Probable phospholipid hydroperoxide glutathione peroxidase 6, mitochondrial [Source:UniProtKB/Swiss-Prot;Acc:O48646] |
| AT4G12320 | CYP706A6 | 1.60 | 1.84 | 0.003 | 0.009 | At4g12320 [Source:UniProtKB/TrEMBL;Acc:Q66GJ1] |
| AT4G13505 | - | 1.88 | 1.25 | 0.000 | 0.021 | other RNA [Source:TAIR;Acc:AT4G13505] |
| AT4G15248 | MIP1A | 2.48 | 2.52 | 0.002 | 0.017 | B-box domain protein 30 [Source:UniProtKB/Swiss-Prot;Acc:Q1G3I2] |
| AT4G15690 | GRXS5 | 2.92 | 1.79 | 0.001 | 0.031 | Monothiol glutaredoxin-S5 [Source:UniProtKB/Swiss-Prot;Acc:O23420] |
| AT4G15700 | GRXS3 | 3.10 | 1.86 | 0.004 | 0.020 | Monothiol glutaredoxin-S3 [Source:UniProtKB/Swiss-Prot;Acc:O23421] |
| AT4G15975 | - | 5.56 | 1.79 | 0.000 | 0.027 | RING/U-box superfamily protein [Source:TAIR;Acc:AT4G15975] |
| AT4G16860 | RPP4 | 2.66 | 2.25 | 0.000 | 0.009 | Disease resistance protein RPP4 [Source:UniProtKB/Swiss-Prot;Acc:F4JNA9] |
| AT4G16880 | - | 3.28 | 2.51 | 0.000 | 0.001 | Leucine-rich repeat (LRR) family protein [Source:UniProtKB/TrEMBL;Acc:F4JNB0] |
| AT4G16890 | SNC1 | 1.28 | 1.28 | 0.001 | 0.001 | disease resistance protein (TIR-NBS-LRR class), putative [Source:TAIR;Acc:AT4G16890] |
| AT4G16920 | - | 1.62 | 1.70 | 0.000 | 0.000 | Disease resistance protein (TIR-NBS-LRR class) family [Source:TAIR;Acc:AT4G16920] |
| AT4G16950 | RPP5 | 1.78 | 1.72 | 0.001 | 0.000 | Disease resistance protein RPP5 [Source:UniProtKB/Swiss-Prot;Acc:F4JNB7] |
| AT4G16957 | - | 2.58 | 1.16 | 0.007 | 0.023 | SUPPRESSOR OF, CONSTITUTIVE protein [Source:UniProtKB/TrEMBL;Acc:A0A1P8B3D3] |
| AT4G17100 | - | 1.27 | 1.28 | 0.012 | 0.029 | CONTAINS InterPro DOMAIN/s: Endoribonuclease XendoU (InterPro:IPR018998); Ha. [Source:TAIR;Acc:AT4G17100] |

| | | | | | | |
|---|---|---|---|---|---|---|
| AT4G17250 | - | 1.94 | 1.10 | 0.003 | 0.025 | AT4g17250/dl4660w [Source:UniProtKB/TrEMBL;Acc:Q93ZA8] |
| AT4G18010 | IP5P2 | 2.12 | 1.16 | 0.002 | 0.008 | Type I inositol polyphosphate 5-phosphatase 2 [Source:UniProtKB/Swiss-Prot;Acc:Q9FUR2] |
| AT4G18205 | PUP22 | 3.89 | 2.13 | 0.004 | 0.033 | Probable purine permease 22 [Source:UniProtKB/Swiss-Prot;Acc:Q8RY74] |
| AT4G18422 | - | 15.52 | 11.46 | 0.000 | 0.002 | unknown protein; FUNCTIONS IN: molecular_function unknown; INVOLVED IN: biological_process unknown; LOCATED IN: cellular_component unknown; Ha. [Source:TAIR;Acc:AT4G18422] |
| AT4G20830 | - | 2.95 | 1.53 | 0.009 | 0.030 | Berberine bridge enzyme-like 19 [Source:UniProtKB/Swiss-Prot;Acc:Q9SVG4] |
| AT4G20860 | FAD-OXR | 2.14 | 1.17 | 0.006 | 0.030 | Berberine bridge enzyme-like 22 [Source:UniProtKB/Swiss-Prot;Acc:Q9SUC6] |
| AT4G21200 | GA2OX8 | 2.89 | 2.92 | 0.017 | 0.017 | Gibberellin 2-beta-dioxygenase 8 [Source:UniProtKB/Swiss-Prot;Acc:O49561] |
| AT4G21380 | SD18 | 2.25 | 2.42 | 0.002 | 0.006 | Receptor-like serine/threonine-protein kinase SD1-8 [Source:UniProtKB/Swiss-Prot;Acc:O81905] |
| AT4G22980 | - | 1.51 | 1.14 | 0.001 | 0.001 | Molybdenum cofactor sulfurase-like protein [Source:UniProtKB/TrEMBL;Acc:O82746] |
| AT4G23170 | CRK9 | 2.55 | 1.31 | 0.000 | 0.038 | Putative cysteine-rich receptor-like protein kinase 9 [Source:UniProtKB/Swiss-Prot;Acc:O65469] |
| AT4G23180 | CRK10 | 2.22 | 1.54 | 0.001 | 0.048 | Cysteine-rich receptor-like protein kinase 10 [Source:UniProtKB/Swiss-Prot;Acc:Q8GYA4] |
| AT4G23220 | CRK14 | 3.58 | 2.53 | 0.017 | 0.028 | Cysteine-rich receptor-like protein kinase 14 [Source:UniProtKB/Swiss-Prot;Acc:Q8H199] |
| AT4G24230 | ACBP3 | 1.57 | 1.45 | 0.012 | 0.012 | Acyl-CoA-binding domain 3 [Source:UniProtKB/TrEMBL;Acc:B3H4G6] |
| AT4G25110 | AMC2 | 1.31 | 1.16 | 0.041 | 0.049 | Metacaspase-2 [Source:UniProtKB/Swiss-Prot;Acc:Q7XJE5] |
| AT4G25480 | DREB1A | 9.30 | 10.60 | 0.049 | 0.024 | Dehydration-responsive element-binding protein 1A [Source:UniProtKB/Swiss-Prot;Acc:Q9M0L0] |
| AT4G26130 | - | 1.01 | 1.25 | 0.002 | 0.002 | AT4g26130/F20B18_240 [Source:UniProtKB/TrEMBL;Acc:Q9SZI4] |
| AT4G26370 | - | 1.33 | 1.38 | 0.001 | 0.020 | Antitermination NusB domain-containing protein [Source:UniProtKB/TrEMBL;Acc:Q93XY7] |
| AT4G26530 | FBA5 | 2.58 | 2.71 | 0.003 | 0.003 | Fructose-bisphosphate aldolase [Source:UniProtKB/TrEMBL;Acc:A0A178V385] |
| AT4G27290 | - | 4.19 | 3.90 | 0.015 | 0.028 | Serine/threonine-protein kinase [Source:UniProtKB/TrEMBL;Acc:A0A178V0J5] |
| AT4G27310 | - | 1.32 | 1.29 | 0.000 | 0.021 | BBX28 [Source:UniProtKB/TrEMBL;Acc:A0A178V4D3] |
| AT4G28140 | ERF054 | 11.47 | 10.05 | 0.001 | 0.027 | Ethylene-responsive transcription factor ERF054 [Source:UniProtKB/Swiss-Prot;Acc:Q9M0J3] |
| AT4G28350 | LECRK72 | 3.35 | 1.73 | 0.000 | 0.006 | Probable L-type lectin-domain containing receptor kinase VII.2 [Source:UniProtKB/Swiss-Prot;Acc:O49445] |
| AT4G30280 | XTH18 | 4.94 | 1.95 | 0.007 | 0.049 | Probable xyloglucan endotransglucosylase/hydrolase protein 18 [Source:UniProtKB/Swiss-Prot;Acc:Q9M0D2] |
| AT4G33300 | ADR1-L1 | 2.29 | 1.18 | 0.001 | 0.010 | Probable disease resistance protein At4g33300 [Source:UniProtKB/Swiss-Prot;Acc:Q9SZA7] |
| AT4G34138 | UGT73B1 | 1.36 | 1.96 | 0.002 | 0.002 | Glycosyltransferase (Fragment) [Source:UniProtKB/TrEMBL;Acc:W8PUI6] |
| AT4G34150 | - | 3.00 | 1.02 | 0.000 | 0.024 | AT4g34150/F28A23_90 [Source:UniProtKB/TrEMBL;Acc:Q945K9] |
| AT4G34380 | - | 12.19 | 7.70 | 0.001 | 0.037 | At4g34380 [Source:UniProtKB/TrEMBL;Acc:Q9SZ03] |
| AT4G34550 | - | 4.58 | 5.70 | 0.017 | 0.008 | At4g34550 [Source:UniProtKB/TrEMBL;Acc:O65680] |
| AT4G37220 | - | 2.45 | 2.62 | 0.004 | 0.002 | Cold-regulated 413 plasma membrane protein 4 [Source:UniProtKB/Swiss-Prot;Acc:O23164] |
| AT4G40065 | - | 2.83 | 2.86 | 0.031 | 0.043 | other RNA [Source:TAIR;Acc:AT4G40065] |
| AT5G01600 | FER1 | 1.61 | 1.51 | 0.023 | 0.020 | Ferritin-1, chloroplastic [Source:UniProtKB/Swiss-Prot;Acc:Q39101] |
| AT5G02160 | - | 1.07 | 1.44 | 0.003 | 0.041 | AT5g02160 [Source:UniProtKB/TrEMBL;Acc:Q9FPH2] |
| AT5G02810 | APRR7 | 1.23 | 1.82 | 0.001 | 0.005 | Two-component response regulator-like APRR7 [Source:UniProtKB/Swiss-Prot;Acc:Q93WK5] |
| AT5G03240 | UBQ3 | 1.12 | 1.32 | 0.002 | 0.010 | Ubiquitin 4 [Source:UniProtKB/TrEMBL;Acc:A0A1P8BGQ7] |
| AT5G03470 | B'ALPHA | 1.00 | 1.56 | 0.008 | 0.043 | Serine/threonine protein phosphatase 2A regulatory subunit [Source:UniProtKB/TrEMBL;Acc:A0A178UNX1] |
| AT5G05410 | DREB2A | 3.13 | 3.70 | 0.005 | 0.002 | Dehydration-responsive element-binding protein 2A [Source:UniProtKB/Swiss-Prot;Acc:O82132] |
| AT5G06690 | WCRKC1 | 2.17 | 2.86 | 0.003 | 0.032 | WCRKC thioredoxin 1 [Source:UniProtKB/TrEMBL;Acc:F4K3Y1] |
| AT5G11250 | - | 1.65 | 1.62 | 0.018 | 0.009 | Disease resistance protein (TIR-NBS-LRR class) [Source:UniProtKB/TrEMBL;Acc:Q9LFN1] |

| | | | | | | |
|---|---|---|---|---|---|---|
| AT5G11260 | HY5 | 1.85 | 2.85 | 0.003 | 0.011 | Basic-leucine zipper (bZIP) transcription factor family protein [Source:TAIR;Acc:AT5G11260] |
| AT5G12940 | - | 2.59 | 1.79 | 0.000 | 0.019 | Leucine-rich repeat (LRR) family protein [Source:UniProtKB/TrEMBL;Acc:Q9LXU5] |
| AT5G14070 | GRXC8 | 1.67 | 1.04 | 0.036 | 0.037 | Glutaredoxin-C8 [Source:UniProtKB/Swiss-Prot;Acc:Q8LF89] |
| AT5G16820 | HSFA1B | 1.34 | 1.18 | 0.007 | 0.011 | Heat stress transcription factor A-1b [Source:UniProtKB/Swiss-Prot;Acc:O81821] |
| AT5G23810 | AAP7 | 2.26 | 2.47 | 0.002 | 0.001 | Probable amino acid permease 7 [Source:UniProtKB/Swiss-Prot;Acc:Q9FF99] |
| AT5G24460 | - | 1.30 | 1.42 | 0.000 | 0.003 | RING-H2 zinc finger protein [Source:UniProtKB/TrEMBL;Acc:Q9FGE4] |
| AT5G24470 | APRR5 | 6.18 | 6.56 | 0.003 | 0.040 | Two-component response regulator-like APRR5 [Source:UniProtKB/Swiss-Prot;Acc:Q6LA42] |
| AT5G24770 | VSP2 | 1.64 | 1.40 | 0.004 | 0.005 | Vegetative storage protein 2 [Source:UniProtKB/Swiss-Prot;Acc:O82122] |
| AT5G28630 | - | 2.19 | 2.01 | 0.017 | 0.019 | Glycine-rich protein [Source:UniProtKB/TrEMBL;Acc:Q8VXY1] |
| AT5G38510 | RBL9 | 1.02 | 1.12 | 0.001 | 0.002 | RHOMBOID-like protein 9, chloroplastic [Source:UniProtKB/Swiss-Prot;Acc:Q9FFX0] |
| AT5G39020 | - | 3.42 | 1.73 | 0.002 | 0.049 | Probable receptor-like protein kinase At5g39020 [Source:UniProtKB/Swiss-Prot;Acc:Q9FID6] |
| AT5G40500 | - | 1.49 | 1.30 | 0.006 | 0.008 | Uncharacterized protein At5g40500 [Source:UniProtKB/TrEMBL;Acc:Q8VZ30] |
| AT5G41100 | - | 2.52 | 1.04 | 0.000 | 0.042 | At5g41100 [Source:UniProtKB/TrEMBL;Acc:A4FVS2] |
| AT5G43260 | - | 1.05 | 1.19 | 0.013 | 0.011 | Chaperone protein dnaJ-like protein [Source:UniProtKB/TrEMBL;Acc:Q94CB5] |
| AT5G43440 | - | 2.68 | 2.19 | 0.001 | 0.021 | 1-aminocyclopropane-1-carboxylate oxidase homolog 9 [Source:UniProtKB/Swiss-Prot;Acc:Q9LSW7] |
| AT5G44210 | ERF9 | 1.73 | 2.13 | 0.011 | 0.015 | Ethylene-responsive transcription factor 9 [Source:UniProtKB/Swiss-Prot;Acc:Q9FE67] |
| AT5G44585 | - | 5.47 | 3.04 | 0.007 | 0.037 | unknown protein; FUNCTIONS IN: molecular_function unknown; INVOLVED IN: biological_process unknown; LOCATED IN: endomembrane system; Ha. [Source:TAIR;Acc:AT5G44585] |
| AT5G45820 | CIPK20 | 1.70 | 2.06 | 0.017 | 0.010 | CBL-interacting serine/threonine-protein kinase 20 [Source:UniProtKB/Swiss-Prot;Acc:Q9FJ54] |
| AT5G45830 | DOG1 | 3.22 | 3.61 | 0.006 | 0.000 | delay of germination 1 [Source:TAIR;Acc:AT5G45830] |
| AT5G47240 | atnudt8 | 4.50 | 3.83 | 0.000 | 0.017 | nudix hydrolase homolog 8 [Source:TAIR;Acc:AT5G47240] |
| AT5G48540 | CRRSP55 | 3.23 | 1.26 | 0.000 | 0.002 | Cysteine-rich repeat secretory protein 55 [Source:UniProtKB/Swiss-Prot;Acc:Q9LV60] |
| AT5G50010 | BHLH145 | 2.86 | 2.99 | 0.048 | 0.035 | Transcription factor bHLH145 [Source:UniProtKB/Swiss-Prot;Acc:Q9FGB0] |
| AT5G50450 | - | 2.42 | 3.49 | 0.007 | 0.010 | F-box protein At5g50450 [Source:UniProtKB/Swiss-Prot;Acc:Q9FK27] |
| AT5G52250 | RUP1 | 2.36 | 3.36 | 0.004 | 0.007 | WD repeat-containing protein RUP1 [Source:UniProtKB/Swiss-Prot;Acc:Q9LTJ6] |
| AT5G54470 | - | 3.10 | 3.30 | 0.000 | 0.000 | BBX29 [Source:UniProtKB/TrEMBL;Acc:A0A178UJV0] |
| AT5G54710 | - | 2.36 | 1.12 | 0.002 | 0.019 | Ankyrin repeat family protein [Source:TAIR;Acc:AT5G54710] |
| AT5G54960 | PDC2 | 2.39 | 2.33 | 0.008 | 0.032 | Pyruvate decarboxylase 2 [Source:UniProtKB/Swiss-Prot;Acc:Q9FFT4] |
| AT5G55970 | - | 1.20 | 1.44 | 0.021 | 0.009 | RING/U-box superfamily protein [Source:UniProtKB/TrEMBL;Acc:Q8LES9] |
| AT5G57630 | CIPK21 | 1.71 | 2.30 | 0.002 | 0.037 | CBL-interacting serine/threonine-protein kinase 21 [Source:UniProtKB/Swiss-Prot;Acc:Q94CG0] |
| AT5G57640 | - | 2.79 | 2.93 | 0.003 | 0.013 | GCK domain-containing protein [Source:UniProtKB/TrEMBL;Acc:Q9FKK9] |
| AT5G59030 | COPT1 | 1.31 | 1.08 | 0.006 | 0.002 | Copper transporter 1 [Source:UniProtKB/Swiss-Prot;Acc:Q39065] |
| AT5G59070 | - | 4.63 | 3.83 | 0.009 | 0.014 | UDP-Glycosyltransferase superfamily protein [Source:UniProtKB/TrEMBL;Acc:F4KHR9] |
| AT5G60900 | RLK1 | 2.10 | 1.89 | 0.001 | 0.015 | receptor-like protein kinase 1 [Source:TAIR;Acc:AT5G60900] |
| AT5G62350 | - | 1.57 | 1.88 | 0.001 | 0.013 | Plant invertase/pectin methylesterase inhibitor superfamily protein [Source:UniProtKB/TrEMBL;Acc:Q9LVA4] |
| AT5G63160 | BT1 | 1.17 | 1.89 | 0.033 | 0.009 | BTB/POZ and TAZ domain-containing protein 1 [Source:UniProtKB/Swiss-Prot;Acc:Q9FMK7] |
| AT5G63780 | SHA1 | 1.19 | 1.49 | 0.004 | 0.002 | At5g63780 [Source:UniProtKB/TrEMBL;Acc:Q8GUG6] |
| AT5G63790 | ANAC102 | 2.21 | 1.02 | 0.002 | 0.026 | NAC domain containing protein 102 [Source:TAIR;Acc:AT5G63790] |
| AT5G64860 | DPE1 | 1.86 | 1.41 | 0.003 | 0.017 | 4-alpha-glucanotransferase DPE1, chloroplastic/amyloplastic [Source:UniProtKB/Swiss-Prot;Acc:Q9LV91] |

| | | | | | | |
|---|---|---|---|---|---|---|
| AT5G66070 | - | 4.09 | 1.59 | 0.005 | 0.036 | RING/U-box superfamily protein [Source:UniProtKB/TrEMBL;Acc:F4JZ26] |
| **'280-0d and 310-0d'_DEGs down-regulated** | | | | | | |
| AT1G01430 | TBL25 | -2.27 | -1.64 | 0.000 | 0.007 | Mannan O-acetyltransferase 3 [Source:UniProtKB/TrEMBL;Acc:A0A346P850] |
| AT1G01840 | - | -1.53 | -1.32 | 0.005 | 0.005 | AP2-like ethylene-responsive transcription factor SNZ [Source:UniProtKB/TrEMBL;Acc:Q9LQ71] |
| AT1G02850 | BGLU11 | -1.68 | -1.45 | 0.010 | 0.006 | Beta-glucosidase 11 [Source:UniProtKB/Swiss-Prot;Acc:B3H5Q1] |
| AT1G02870 | - | -1.21 | -1.18 | 0.007 | 0.035 | At1g02870 [Source:UniProtKB/TrEMBL;Acc:Q8RWK5] |
| AT1G03110 | TRM82 | -1.77 | -1.27 | 0.010 | 0.013 | tRNA (guanine-N(7)-)-methyltransferase non-catalytic subunit [Source:UniProtKB/TrEMBL;Acc:A0A178VZV6] |
| AT1G03360 | RRP4 | -1.65 | -1.44 | 0.005 | 0.016 | RRP4 [Source:UniProtKB/TrEMBL;Acc:A0A178W3N1] |
| AT1G03530 | ATNAF1 | -1.77 | -1.72 | 0.003 | 0.013 | NAF1 [Source:UniProtKB/TrEMBL;Acc:A0A178WI11] |
| AT1G03930 | CKL9 | -1.28 | -1.08 | 0.008 | 0.022 | Casein kinase 1-like protein 9 [Source:UniProtKB/Swiss-Prot;Acc:Q9ZWB3] |
| AT1G04040 | - | -3.00 | -1.60 | 0.005 | 0.005 | At1g04040/F21M11_2 [Source:UniProtKB/TrEMBL;Acc:Q9ZWC4] |
| AT1G04120 | ABCC5 | -1.44 | -1.38 | 0.014 | 0.001 | MRP5 [Source:UniProtKB/TrEMBL;Acc:A0A178WGC2] |
| AT1G04360 | ATL1 | -1.11 | -1.57 | 0.009 | 0.049 | RING-H2 finger protein ATL1 [Source:UniProtKB/Swiss-Prot;Acc:P93823] |
| AT1G04520 | CRRSP3 | -1.07 | -1.32 | 0.002 | 0.013 | Cysteine-rich repeat secretory protein 3 [Source:UniProtKB/Swiss-Prot;Acc:Q6NM73] |
| AT1G04770 | - | -1.64 | -1.46 | 0.022 | 0.029 | Protein SULFUR DEFICIENCY-INDUCED 2 [Source:UniProtKB/Swiss-Prot;Acc:Q8L730] |
| AT1G05460 | SDE3 | -1.35 | -1.19 | 0.016 | 0.022 | Probable RNA helicase SDE3 [Source:UniProtKB/Swiss-Prot;Acc:Q8GYD9] |
| AT1G05560 | UGT1 | -3.29 | -1.60 | 0.007 | 0.029 | UDP-glucosyltransferase 75B1 [Source:TAIR;Acc:AT1G05560] |
| AT1G06360 | - | -1.70 | -1.02 | 0.003 | 0.007 | Fatty acid desaturase family protein [Source:TAIR;Acc:AT1G06360] |
| AT1G06380 | - | -1.32 | -1.10 | 0.004 | 0.046 | Ribosomal protein L1p/L10e family [Source:UniProtKB/TrEMBL;Acc:Q9LMI1] |
| AT1G06720 | - | -2.35 | -1.39 | 0.037 | 0.024 | P-loop containing nucleoside triphosphate hydrolases superfamily protein [Source:TAIR;Acc:AT1G06720] |
| AT1G07720 | KCS3 | -3.21 | -1.25 | 0.000 | 0.013 | 3-ketoacyl-CoA synthase 3 [Source:TAIR;Acc:AT1G07720] |
| AT1G08230 | GAT1 | -1.31 | -1.28 | 0.001 | 0.001 | GABA transporter 1 [Source:UniProtKB/Swiss-Prot;Acc:F4HW02] |
| AT1G08580 | - | -1.47 | -1.29 | 0.030 | 0.000 | At1g08580 [Source:UniProtKB/TrEMBL;Acc:Q9FRS7] |
| AT1G08610 | - | -2.40 | -2.19 | 0.004 | 0.021 | Pentatricopeptide repeat-containing protein At1g08610 [Source:UniProtKB/Swiss-Prot;Acc:Q9FRS4] |
| AT1G09220 | PCMP-E25 | -1.43 | -1.15 | 0.035 | 0.019 | Pentatricopeptide repeat-containing protein At1g09220, mitochondrial [Source:UniProtKB/Swiss-Prot;Acc:Q680Z7] |
| AT1G10470 | ARR4 | -1.76 | -1.65 | 0.001 | 0.010 | Two-component response regulator ARR4 [Source:UniProtKB/Swiss-Prot;Acc:O82798] |
| AT1G11700 | - | -2.19 | -2.00 | 0.002 | 0.040 | At1g11700 [Source:UniProtKB/TrEMBL;Acc:Q9SAA7] |
| AT1G11710 | - | -2.04 | -2.62 | 0.029 | 0.000 | Pentatricopeptide repeat-containing protein At1g11710, mitochondrial [Source:UniProtKB/Swiss-Prot;Acc:Q9SAA6] |
| AT1G12110 | NPF6.3 | -1.90 | -1.13 | 0.023 | 0.040 | NRT1.1 [Source:UniProtKB/TrEMBL;Acc:A0A178W8F7] |
| AT1G12330 | - | -2.75 | -1.55 | 0.001 | 0.041 | Cyclin-dependent kinase-like protein [Source:UniProtKB/TrEMBL;Acc:Q9LNB2] |
| AT1G13810 | - | -1.54 | -1.51 | 0.035 | 0.038 | Restriction endonuclease, type II-like superfamily protein [Source:UniProtKB/TrEMBL;Acc:Q5XVK9] |
| AT1G14060 | - | -1.37 | -1.62 | 0.001 | 0.010 | F7A19.14 protein [Source:UniProtKB/TrEMBL;Acc:Q9XI82] |
| AT1G14460 | - | -1.79 | -1.11 | 0.004 | 0.010 | Protein STICHEL-like 1 [Source:UniProtKB/Swiss-Prot;Acc:F4HW65] |
| AT1G14600 | - | -2.40 | -1.70 | 0.028 | 0.001 | Putative Myb family transcription factor At1g14600 [Source:UniProtKB/Swiss-Prot;Acc:Q700D9] |
| AT1G14840 | MAP70.4 | -1.73 | -1.13 | 0.045 | 0.033 | Microtubule-associated protein 70-4 [Source:UniProtKB/Swiss-Prot;Acc:Q9LQU7] |
| AT1G15250 | RPL37A | -1.53 | -1.04 | 0.003 | 0.014 | 60S ribosomal protein L37-1 [Source:UniProtKB/Swiss-Prot;Acc:Q8LFH7] |
| AT1G15260 | - | -2.46 | -1.42 | 0.008 | 0.015 | LOW protein: ATP-dependent RNA helicase-like protein [Source:UniProtKB/TrEMBL;Acc:Q8VY60] |
| AT1G15510 | PCMP-H73 | -2.23 | -1.29 | 0.021 | 0.010 | Pentatricopeptide repeat-containing protein At1g15510, chloroplastic [Source:UniProtKB/Swiss-Prot;Acc:Q9M9E2] |

| | | | | | |
|---|---|---|---|---|---|
| AT1G15550 | GA3OX1 | -4.42 | -3.31 | 0.031 | 0.039 Gibberellin 3-beta-dioxygenase 1 [Source:UniProtKB/Swiss-Prot;Acc:Q39103] |
| AT1G16400 | CYP79F2 | -1.31 | -2.00 | 0.038 | 0.012 Hexahomomethionine N-hydroxylase [Source:UniProtKB/Swiss-Prot;Acc:Q9FUY7] |
| AT1G16750 | - | -2.70 | -1.23 | 0.009 | 0.015 At1g16750/F19K19_26 [Source:UniProtKB/TrEMBL;Acc:Q8L7U6] |
| AT1G18090 | - | -1.34 | -1.50 | 0.002 | 0.006 5'-3' exonuclease family protein [Source:UniProtKB/TrEMBL;Acc:Q94A33] |
| AT1G18800 | NRP2 | -1.29 | -1.27 | 0.018 | 0.000 NRP2 [Source:UniProtKB/TrEMBL;Acc:A0A178W8U0] |
| AT1G19350 | BES1 | -1.32 | -1.03 | 0.000 | 0.005 Brassinosteroid signaling positive regulator (BZR1) family protein [Source:UniProtKB/TrEMBL;Acc:F4HP45] |
| AT1G19630 | CYP722A1 | -2.70 | -1.78 | 0.009 | 0.026 Cytochrome P450, family 722, subfamily A, polypeptide 1 [Source:UniProtKB/TrEMBL;Acc:F4HP86] |
| AT1G19640 | JMT | -1.56 | -2.29 | 0.009 | 0.002 Jasmonate O-methyltransferase [Source:UniProtKB/Swiss-Prot;Acc:Q9AR07] |
| AT1G20370 | - | -1.23 | -1.12 | 0.019 | 0.014 F14O10.3 protein [Source:UniProtKB/TrEMBL;Acc:Q9LN29] |
| AT1G21520 | - | -1.43 | -1.16 | 0.041 | 0.016 Uncharacterized protein At1g21520/F24J8_4 [Source:UniProtKB/TrEMBL;Acc:Q8GYE6] |
| AT1G23280 | - | -1.94 | -1.33 | 0.000 | 0.001 Protein MAK16 homolog [Source:UniProtKB/TrEMBL;Acc:F4I4Q1] |
| AT1G23480 | CSLA3 | -2.43 | -1.16 | 0.012 | 0.043 Glycosyltransferase (Fragment) [Source:UniProtKB/TrEMBL;Acc:W8Q7B4] |
| AT1G24020 | MLP423 | -1.16 | -1.37 | 0.020 | 0.025 MLP-like protein 423 [Source:UniProtKB/Swiss-Prot;Acc:Q93VR4] |
| AT1G25425 | CLE43 | -1.75 | -1.85 | 0.009 | 0.041 CLAVATA3/ESR (CLE)-related protein 43 [Source:UniProtKB/Swiss-Prot;Acc:Q6IWB1] |
| AT1G25440 | COL16 | -2.81 | -1.50 | 0.000 | 0.028 Zinc finger protein CONSTANS-LIKE 16 [Source:UniProtKB/Swiss-Prot;Acc:Q8RWD0] |
| AT1G26770 | ATEXPA10 | -2.18 | -1.78 | 0.006 | 0.010 Expansin [Source:UniProtKB/TrEMBL;Acc:F4HPC1] |
| AT1G27050 | ATHB54 | -1.98 | -1.32 | 0.007 | 0.002 Uncharacterized protein At1g27050 [Source:UniProtKB/Swiss-Prot;Acc:P0CJ66] |
| AT1G28395 | - | -1.65 | -1.40 | 0.019 | 0.004 At1g28395 [Source:UniProtKB/TrEMBL;Acc:Q8LEQ7] |
| AT1G28440 | HSL1 | -2.28 | -1.43 | 0.000 | 0.016 Receptor-like protein kinase HSL1 [Source:UniProtKB/Swiss-Prot;Acc:Q9SGP2] |
| AT1G28660 | - | -2.81 | -1.48 | 0.025 | 0.034 GDSL esterase/lipase At1g28660 [Source:UniProtKB/Swiss-Prot;Acc:Q9FPE4] |
| AT1G29660 | - | -2.95 | -1.65 | 0.000 | 0.001 GDSL esterase/lipase At1g29660 [Source:UniProtKB/Swiss-Prot;Acc:Q9C7N5] |
| AT1G29720 | RFK1 | -2.21 | -1.76 | 0.001 | 0.016 Probable LRR receptor-like serine/threonine-protein kinase At1g29720 [Source:UniProtKB/Swiss-Prot;Acc:Q9ASQ6] |
| AT1G29900 | CARB | -1.78 | -1.01 | 0.041 | 0.006 Carbamoyl-phosphate synthase large chain, chloroplastic [Source:UniProtKB/Swiss-Prot;Acc:Q42601] |
| AT1G29965 | RPL18AA | -1.56 | -1.49 | 0.002 | 0.023 60S ribosomal protein L18a-1 [Source:UniProtKB/Swiss-Prot;Acc:Q8L7K0] |
| AT1G30110 | NUDT25 | -1.11 | -1.36 | 0.003 | 0.026 Nudix hydrolase 25 [Source:UniProtKB/Swiss-Prot;Acc:Q9C6Z2] |
| AT1G30160 | - | -1.41 | -1.32 | 0.026 | 0.026 At1g30160 [Source:UniProtKB/TrEMBL;Acc:Q6DBP6] |
| AT1G30230 | - | -1.12 | -1.04 | 0.000 | 0.000 Translation elongation factor EF1B/ribosomal protein S6 family protein [Source:UniProtKB/TrEMBL;Acc:A8MRC4] |
| AT1G30530 | UGT78D1 | -2.11 | -1.84 | 0.004 | 0.006 Glycosyltransferase (Fragment) [Source:UniProtKB/TrEMBL;Acc:W8PVA4] |
| AT1G30960 | ERG | -2.76 | -1.64 | 0.003 | 0.005 GTP-binding protein ERG [Source:UniProtKB/Swiss-Prot;Acc:O82653] |
| AT1G31173 | MIR167D | -1.22 | -1.89 | 0.019 | 0.003 MIR167D; miRNA [Source:TAIR;Acc:AT1G31173] |
| AT1G31835 | - | -1.11 | -1.36 | 0.037 | 0.049 unknown protein; FUNCTIONS IN: molecular_function unknown; INVOLVED IN: biological_process unknown; LOCATED IN: endomembrane system; Ha. [Source:TAIR;Acc:AT1G31835] |
| AT1G31970 | RH5 | -1.75 | -1.44 | 0.002 | 0.001 STRS1 [Source:UniProtKB/TrEMBL;Acc:A0A178W5E1] |
| AT1G34418 | - | -1.53 | -1.55 | 0.002 | 0.036 other RNA [Source:TAIR;Acc:AT1G34418] |
| AT1G34640 | - | -1.87 | -1.66 | 0.011 | 0.007 At1g34640 [Source:UniProtKB/TrEMBL;Acc:Q9LNM6] |
| AT1G36280 | - | -1.53 | -1.17 | 0.022 | 0.001 Adenylosuccinate lyase [Source:UniProtKB/TrEMBL;Acc:Q8GUN7] |
| AT1G47500 | RBP47C' | -1.17 | -1.19 | 0.016 | 0.006 Polyadenylate-binding protein RBP47C' [Source:UniProtKB/Swiss-Prot;Acc:Q9SX80] |
| AT1G48300 | - | -2.83 | -2.55 | 0.006 | 0.001 unknown protein; FUNCTIONS IN: molecular_function unknown; INVOLVED IN: biological_process unknown; LOCATED IN: endomembrane system; EXPRESSED IN: 24 plant structures; EXPRESSED DURING: 15 growth stages; Ha. [Source:TAIR;Acc:AT1G48300] |

| | | | | | | |
|---|---|---|---|---|---|---|
| AT1G48480 | RKL1 | -1.57 | -1.58 | 0.003 | 0.025 | Probable inactive receptor kinase At1g48480 [Source:UniProtKB/Swiss-Prot;Acc:Q9LP77] |
| AT1G48770 | - | -1.35 | -1.15 | 0.001 | 0.036 | Uncharacterized protein F11I4_6 [Source:UniProtKB/TrEMBL;Acc:Q9C745] |
| AT1G50420 | SCL3 | -2.98 | -2.02 | 0.000 | 0.004 | Scarecrow-like protein 3 [Source:UniProtKB/Swiss-Prot;Acc:Q9LPR8] |
| AT1G51060 | HTA10 | -1.16 | -1.15 | 0.043 | 0.000 | Probable histone H2A.1 [Source:UniProtKB/Swiss-Prot;Acc:Q9C681] |
| AT1G52240 | ROPGEF11 | -2.04 | -1.86 | 0.019 | 0.027 | ROPGEF11 [Source:UniProtKB/TrEMBL;Acc:A0A178W8V0] |
| AT1G52905 | - | -1.80 | -1.48 | 0.001 | 0.002 | At1g52905 [Source:UniProtKB/TrEMBL;Acc:Q1EBU8] |
| AT1G52930 | BRIX1-2 | -1.24 | -1.16 | 0.010 | 0.000 | Ribosome biogenesis protein BRX1 homolog 2 [Source:UniProtKB/Swiss-Prot;Acc:Q9C928] |
| AT1G53645 | - | -1.19 | -1.11 | 0.006 | 0.010 | At1g53640/F22G10.8 [Source:UniProtKB/TrEMBL;Acc:Q9C8L9] |
| AT1G54010 | GLL23 | -1.45 | -1.78 | 0.027 | 0.008 | Inactive GDSL esterase/lipase-like protein 23 [Source:UniProtKB/Swiss-Prot;Acc:Q8W4H8] |
| AT1G54120 | - | -1.92 | -2.58 | 0.027 | 0.045 | F15I1.22 [Source:UniProtKB/TrEMBL;Acc:Q9SYH0] |
| AT1G54690 | HTA3 | -1.67 | -1.38 | 0.018 | 0.041 | Probable histone H2AXb [Source:UniProtKB/Swiss-Prot;Acc:Q9S9K7] |
| AT1G54820 | - | -1.12 | -1.02 | 0.010 | 0.031 | Protein kinase superfamily protein [Source:UniProtKB/TrEMBL;Acc:F4HYK7] |
| AT1G56110 | NOP56 | -2.01 | -1.45 | 0.018 | 0.035 | At1g56110/T6H22_9 [Source:UniProtKB/TrEMBL;Acc:Q9SGT7] |
| AT1G56210 | HIPP35 | -1.80 | -1.13 | 0.048 | 0.004 | Heavy metal-associated isoprenylated plant protein 35 [Source:UniProtKB/Swiss-Prot;Acc:Q9C7J6] |
| AT1G56580 | SVB | -1.75 | -1.12 | 0.012 | 0.030 | At1g56580/F25P12_18 [Source:UniProtKB/TrEMBL;Acc:Q9FXB0] |
| AT1G60850 | ATRPAC42 | -1.78 | -1.50 | 0.001 | 0.002 | DNA-directed RNA polymerase family protein [Source:UniProtKB/TrEMBL;Acc:Q9C6C2] |
| AT1G61570 | TIM13 | -1.45 | -1.45 | 0.008 | 0.010 | Mitochondrial import inner membrane translocase subunit TIM13 [Source:UniProtKB/Swiss-Prot;Acc:Q9XH48] |
| AT1G61580 | ARP2 | -1.79 | -1.58 | 0.001 | 0.003 | 60S ribosomal protein L3-2 [Source:UniProtKB/Swiss-Prot;Acc:P22738] |
| AT1G61870 | PPR336 | -1.57 | -1.24 | 0.001 | 0.002 | Pentatricopeptide repeat-containing protein At1g61870, mitochondrial [Source:UniProtKB/Swiss-Prot;Acc:Q8LE47] |
| AT1G62150 | - | -1.91 | -3.17 | 0.000 | 0.030 | Mitochondrial transcription termination factor family protein [Source:UniProtKB/TrEMBL;Acc:Q8GWB0] |
| AT1G62370 | - | -3.14 | -1.66 | 0.027 | 0.026 | F2401.10 [Source:UniProtKB/TrEMBL;Acc:O48801] |
| AT1G62560 | FMOGS-OX3 | -1.53 | -1.74 | 0.023 | 0.000 | Flavin-containing monooxygenase FMO GS-OX3 [Source:UniProtKB/Swiss-Prot;Acc:Q9SXE1] |
| AT1G63780 | IMP4 | -1.37 | -1.16 | 0.004 | 0.009 | IMP4 [Source:UniProtKB/TrEMBL;Acc:A0A178WDJ3] |
| AT1G63810 | - | -3.01 | -1.06 | 0.007 | 0.047 | Nucleolar protein [Source:UniProtKB/TrEMBL;Acc:Q0WVM5] |
| AT1G64670 | BDG1 | -2.22 | -1.20 | 0.000 | 0.000 | alpha/beta-Hydrolases superfamily protein [Source:TAIR;Acc:AT1G64670] |
| AT1G64790 | ILA | -1.08 | -1.23 | 0.025 | 0.001 | Protein ILITYHIA [Source:UniProtKB/Swiss-Prot;Acc:F4I893] |
| AT1G65080 | ALB3L2 | -1.79 | -1.48 | 0.012 | 0.005 | ALBINO3-like protein 2, chloroplastic [Source:UniProtKB/Swiss-Prot;Acc:Q8L718] |
| AT1G66350 | RGL1 | -1.91 | -1.30 | 0.001 | 0.002 | DELLA protein RGL1 [Source:UniProtKB/Swiss-Prot;Acc:Q9C8Y3] |
| AT1G67630 | POLA2 | -1.79 | -1.63 | 0.004 | 0.008 | DNA polymerase alpha subunit B [Source:UniProtKB/TrEMBL;Acc:F4HTP2] |
| AT1G68238 | - | -1.51 | -1.76 | 0.003 | 0.050 | Putative uncharacterized protein [Source:UniProtKB/TrEMBL;Acc:Q1G3X3] |
| AT1G68520 | COL6 | -3.32 | -1.78 | 0.012 | 0.007 | Zinc finger protein CONSTANS-LIKE 6 [Source:UniProtKB/Swiss-Prot;Acc:Q8LG76] |
| AT1G68570 | NPF3.1 | -2.07 | -1.84 | 0.037 | 0.022 | Protein NRT1/ PTR FAMILY 3.1 [Source:UniProtKB/Swiss-Prot;Acc:Q9SX20] |
| AT1G68990 | MGP3 | -2.07 | -1.38 | 0.008 | 0.012 | DNA-directed RNA polymerase [Source:UniProtKB/TrEMBL;Acc:F4I0I3] |
| AT1G69040 | ACR4 | -1.60 | -1.12 | 0.014 | 0.022 | ACT domain repeat 4 [Source:UniProtKB/TrEMBL;Acc:F4I0I7] |
| AT1G69070 | - | -1.78 | -1.54 | 0.003 | 0.023 | FUNCTIONS IN: molecular_function unknown; INVOLVED IN: biological_process unknown; LOCATED IN: cellular_component unknown; EXPRESSED IN: 23 plant structures; EXPRESSED DURING: 13 growth stages; CONTAINS InterPro DOMAIN/s: Nop14-like protein (InterPr /.../07276); Ha. [Source:TAIR;Acc:AT1G69070] |
| AT1G69400 | BUB3.3 | -1.23 | -1.01 | 0.033 | 0.049 | Mitotic checkpoint protein BUB3.3 [Source:UniProtKB/Swiss-Prot;Acc:F4I241] |
| AT1G69490 | NAC029 | -2.22 | -1.53 | 0.001 | 0.002 | NAP [Source:UniProtKB/TrEMBL;Acc:A0A178W8K0] |

| | | | | | | |
|---|---|---|---|---|---|---|
| AT1G70550 | - | -1.69 | -1.07 | 0.014 | 0.007 | Protein of Unknown Function (DUF239) [Source:TAIR;Acc:AT1G70550] |
| AT1G72040 | - | -1.47 | -1.04 | 0.000 | 0.039 | At1g72040 [Source:UniProtKB/TrEMBL;Acc:Q501D4] |
| AT1G72470 | ATEXO70D1 | -2.26 | -1.30 | 0.001 | 0.008 | Exocyst subunit Exo70 family protein [Source:UniProtKB/TrEMBL;Acc:Q9C9E5] |
| AT1G73970 | - | -1.04 | -1.02 | 0.008 | 0.004 | unknown protein; Ha. [Source:TAIR;Acc:AT1G73970] |
| AT1G74870 | - | -1.96 | -1.67 | 0.008 | 0.007 | At1g74870 [Source:UniProtKB/TrEMBL;Acc:Q9S7I7] |
| AT1G74930 | ERF018 | -1.60 | -1.90 | 0.029 | 0.015 | Ethylene-responsive transcription factor ERF018 [Source:UniProtKB/Swiss-Prot;Acc:Q9S7L5] |
| AT1G75710 | - | -2.28 | -1.38 | 0.005 | 0.008 | C2H2-like zinc finger protein [Source:UniProtKB/TrEMBL;Acc:Q9LR10] |
| AT1G77030 | RH29 | -2.08 | -2.34 | 0.016 | 0.014 | Putative DEAD-box ATP-dependent RNA helicase 29 [Source:UniProtKB/Swiss-Prot;Acc:O49289] |
| AT1G77460 | CSI3 | -1.24 | -1.73 | 0.033 | 0.034 | Protein CELLULOSE SYNTHASE INTERACTIVE 3 [Source:UniProtKB/Swiss-Prot;Acc:F4I718] |
| AT1G77690 | LAX3 | -1.18 | -1.18 | 0.031 | 0.011 | LAX3 [Source:UniProtKB/TrEMBL;Acc:A0A178W1L1] |
| AT1G78020 | FLZ6 | -1.08 | -1.10 | 0.002 | 0.013 | FCS-Like Zinc finger 6 [Source:UniProtKB/Swiss-Prot;Acc:Q9SGZ8] |
| AT1G78170 | - | -4.19 | -1.63 | 0.000 | 0.023 | E3 ubiquitin-protein ligase [Source:UniProtKB/TrEMBL;Acc:Q84JP5] |
| AT1G78570 | RHM1 | -1.61 | -1.11 | 0.019 | 0.027 | Trifunctional UDP-glucose 4,6-dehydratase/UDP-4-keto-6-deoxy-D-glucose 3,5-epimerase/UDP-4-keto-L-rhamnose-reductase RHM1 [Source:UniProtKB/Swiss-Prot;Acc:Q9SYM5] |
| AT1G79030 | - | -1.71 | -1.10 | 0.008 | 0.021 | Chaperone DnaJ-domain superfamily protein [Source:UniProtKB/TrEMBL;Acc:F4IDH6] |
| AT1G79110 | BRG2 | -2.83 | -1.59 | 0.040 | 0.001 | Probable BOI-related E3 ubiquitin-protein ligase 2 [Source:UniProtKB/Swiss-Prot;Acc:F4IDI6] |
| AT1G79470 | IMPDH | -1.86 | -1.54 | 0.028 | 0.026 | Inosine-5'-monophosphate dehydrogenase 1 [Source:UniProtKB/Swiss-Prot;Acc:P47996] |
| AT1G80130 | - | -1.65 | -2.01 | 0.001 | 0.004 | F18B13.21 protein [Source:UniProtKB/TrEMBL;Acc:Q9SSC6] |
| AT1G80150 | - | -1.61 | -1.77 | 0.024 | 0.006 | Pentatricopeptide repeat-containing protein At1g80150, mitochondrial [Source:UniProtKB/Swiss-Prot;Acc:Q8GW57] |
| AT1G80520 | - | -1.45 | -1.04 | 0.028 | 0.030 | Sterile alpha motif (SAM) domain-containing protein [Source:UniProtKB/TrEMBL;Acc:Q9M8M0] |
| AT1G80640 | - | -1.67 | -1.18 | 0.018 | 0.003 | Probable receptor-like protein kinase At1g80640 [Source:UniProtKB/Swiss-Prot;Acc:Q0V7T5] |
| AT1G80750 | RPL7A | -1.33 | -1.01 | 0.000 | 0.000 | 60S ribosomal protein L7-1 [Source:UniProtKB/Swiss-Prot;Acc:Q9SAI5] |
| AT1G80760 | NIP6-1 | -1.60 | -1.43 | 0.000 | 0.013 | Aquaporin NIP6-1 [Source:UniProtKB/Swiss-Prot;Acc:Q9SAI4] |
| AT2G01505 | CLE16 | -1.77 | -1.35 | 0.016 | 0.030 | CLAVATA3/ESR (CLE)-related protein 16 [Source:UniProtKB/Swiss-Prot;Acc:Q8S8M2] |
| AT2G01680 | - | -1.31 | -1.04 | 0.001 | 0.001 | Ankyrin repeat-containing protein At2g01680 [Source:UniProtKB/Swiss-Prot;Acc:Q9ZU96] |
| AT2G01740 | - | -2.08 | -1.38 | 0.016 | 0.035 | Pentatricopeptide repeat-containing protein At2g01740 [Source:UniProtKB/Swiss-Prot;Acc:Q9ZUA2] |
| AT2G02780 | - | -2.44 | -1.32 | 0.000 | 0.001 | Probable LRR receptor-like serine/threonine-protein kinase At2g02780 [Source:UniProtKB/Swiss-Prot;Acc:C0LGJ9] |
| AT2G02910 | - | -1.51 | -1.11 | 0.010 | 0.003 | At2g02910 [Source:UniProtKB/TrEMBL;Acc:A4IJ37] |
| AT2G04660 | APC2 | -1.18 | -1.60 | 0.022 | 0.007 | APC2 [Source:UniProtKB/TrEMBL;Acc:A0A178VYA9] |
| AT2G10931 | - | -1.66 | -1.08 | 0.009 | 0.039 | Putative uncharacterized protein [Source:UniProtKB/TrEMBL;Acc:Q1PF83] |
| AT2G12462 | - | -1.83 | -1.58 | 0.012 | 0.002 | Sterile alpha motif (SAM) domain protein [Source:UniProtKB/TrEMBL;Acc:Q1G3Q5] |
| AT2G14460 | - | -1.08 | -1.28 | 0.017 | 0.020 | At2g14460 [Source:UniProtKB/TrEMBL;Acc:Q9ZQQ9] |
| AT2G15000 | - | -1.10 | -1.26 | 0.030 | 0.020 | unknown protein; FUNCTIONS IN: molecular_function unknown; INVOLVED IN: biological_process unknown; LOCATED IN: chloroplast; EXPRESSED IN: 22 plant structures; EXPRESSED DURING: 13 growth stages; BEST Arabidopsis thaliana protein match is: unknown p /.../ (TAIR:AT4G34265.2); Ha. [Source:TAIR;Acc:AT2G15000] |
| AT2G15690 | PCMP-H66 | -1.38 | -1.10 | 0.034 | 0.002 | Pentatricopeptide repeat-containing protein At2g15690, mitochondrial [Source:UniProtKB/Swiss-Prot;Acc:Q9ZQE5] |
| AT2G17670 | - | -2.26 | -1.90 | 0.006 | 0.010 | Pentatricopeptide repeat-containing protein At2g17670 [Source:UniProtKB/Swiss-Prot;Acc:Q84J71] |
| AT2G18328 | RL4 | -1.17 | -1.09 | 0.001 | 0.002 | Protein RADIALIS-like 4 [Source:UniProtKB/Swiss-Prot;Acc:Q1G3C4] |
| AT2G18330 | - | -1.25 | -1.20 | 0.007 | 0.007 | AAA-type ATPase family protein [Source:UniProtKB/TrEMBL;Acc:Q9ZPW5] |

| | | | | | | |
|---|---|---|---|---|---|---|
| AT2G19780 | - | -1.74 | -1.47 | 0.035 | 0.049 | Leucine-rich repeat (LRR) family protein [Source:UniProtKB/TrEMBL;Acc:O82202] |
| AT2G20180 | PIF1 | -2.14 | -1.51 | 0.004 | 0.002 | Transcription factor PIF1 [Source:UniProtKB/Swiss-Prot;Acc:Q8GZM7] |
| AT2G20490 | NOP10 | -1.58 | -1.29 | 0.001 | 0.016 | H/ACA ribonucleoprotein complex subunit 3-like protein [Source:UniProtKB/Swiss-Prot;Acc:Q93XX8] |
| AT2G20570 | GPRI1 | -1.72 | -1.35 | 0.000 | 0.016 | GBF's pro-rich region-interacting factor 1 [Source:UniProtKB/TrEMBL;Acc:F4IVF9] |
| AT2G21440 | - | -1.82 | -1.21 | 0.008 | 0.002 | Expressed protein [Source:UniProtKB/TrEMBL;Acc:Q9SJT4] |
| AT2G21790 | RNR1 | -1.76 | -1.35 | 0.024 | 0.034 | Ribonucleoside-diphosphate reductase large subunit [Source:UniProtKB/Swiss-Prot;Acc:Q9SJ20] |
| AT2G22810 | ACS4 | -9.99 | -1.57 | 0.000 | 0.031 | 1-aminocyclopropane-1-carboxylate synthase 4 [Source:UniProtKB/Swiss-Prot;Acc:Q43309] |
| AT2G23430 | KRP1 | -2.28 | -1.81 | 0.018 | 0.004 | Cyclin-dependent kinase inhibitor 1 [Source:UniProtKB/Swiss-Prot;Acc:Q67Y93] |
| AT2G23470 | RUS4 | -1.57 | -1.95 | 0.013 | 0.044 | Protein root UVB sensitive 4 [Source:UniProtKB/Swiss-Prot;Acc:Q67YT8] |
| AT2G23560 | MES7 | -3.32 | -2.22 | 0.010 | 0.012 | MES7 [Source:UniProtKB/TrEMBL;Acc:A0A178VX48] |
| AT2G23690 | - | -3.89 | -1.81 | 0.011 | 0.035 | HTH-type transcriptional regulator [Source:UniProtKB/TrEMBL;Acc:Q680G2] |
| AT2G25580 | PCMP-H75 | -1.75 | -1.13 | 0.012 | 0.017 | MEF8 [Source:UniProtKB/TrEMBL;Acc:A0A178VVM1] |
| AT2G26310 | FAP2 | -3.69 | -1.70 | 0.028 | 0.006 | FAP2 [Source:UniProtKB/TrEMBL;Acc:A0A178VYJ7] |
| AT2G26690 | NPF6.2 | -1.62 | -1.03 | 0.006 | 0.047 | Protein NRT1/ PTR FAMILY 6.2 [Source:UniProtKB/Swiss-Prot;Acc:Q9SZY4] |
| AT2G26860 | - | -1.23 | -1.53 | 0.038 | 0.010 | FBD-associated F-box protein At2g26860 [Source:UniProtKB/Swiss-Prot;Acc:Q84X02] |
| AT2G27060 | - | -2.62 | -2.27 | 0.023 | 0.037 | Leucine-rich repeat protein kinase family protein [Source:UniProtKB/TrEMBL;Acc:F4IVP3] |
| AT2G27230 | LHW | -1.47 | -1.55 | 0.013 | 0.005 | LHW [Source:UniProtKB/TrEMBL;Acc:A0A178VXD3] |
| AT2G27840 | HDT4 | -2.14 | -1.64 | 0.010 | 0.015 | Histone deacetylase HDT4 [Source:UniProtKB/Swiss-Prot;Acc:Q9M4T3] |
| AT2G28150 | - | -2.52 | -1.09 | 0.000 | 0.026 | UPSTREAM OF FLC protein (DUF966) [Source:UniProtKB/TrEMBL;Acc:Q8GYT8] |
| AT2G28330 | SMR11 | -2.19 | -1.20 | 0.013 | 0.005 | Cyclin-dependent protein kinase inhibitor SMR11 [Source:UniProtKB/Swiss-Prot;Acc:Q9SKN7] |
| AT2G28350 | ARF10 | -3.58 | -1.41 | 0.011 | 0.029 | Auxin response factor (Fragment) [Source:UniProtKB/TrEMBL;Acc:C0SV66] |
| AT2G29200 | APUM1 | -1.75 | -1.60 | 0.027 | 0.008 | Pumilio homolog 1 [Source:UniProtKB/Swiss-Prot;Acc:Q9ZW07] |
| AT2G30070 | POT1 | -1.34 | -1.22 | 0.003 | 0.006 | Potassium transporter 1 [Source:UniProtKB/Swiss-Prot;Acc:O22397] |
| AT2G31060 | - | -1.90 | -1.08 | 0.024 | 0.026 | Elongation factor family protein [Source:UniProtKB/TrEMBL;Acc:F4IPW5] |
| AT2G31160 | LSH3 | -1.25 | -1.72 | 0.032 | 0.013 | Protein LIGHT-DEPENDENT SHORT HYPOCOTYLS 3 [Source:UniProtKB/Swiss-Prot;Acc:O82268] |
| AT2G31730 | BHLH154 | -1.33 | -1.04 | 0.038 | 0.040 | Transcription factor bHLH154 [Source:UniProtKB/Swiss-Prot;Acc:Q7XJU1] |
| AT2G32010 | IP5P7 | -3.50 | -1.86 | 0.024 | 0.037 | CVL1 [Source:UniProtKB/TrEMBL;Acc:A0A178VNT8] |
| AT2G32220 | RPL27A | -1.46 | -1.82 | 0.036 | 0.040 | 60S ribosomal protein L27-1 [Source:UniProtKB/Swiss-Prot;Acc:Q9SKX8] |
| AT2G32487 | - | -1.76 | -1.40 | 0.006 | 0.012 | unknown protein; FUNCTIONS IN: molecular_function unknown; INVOLVED IN: biological_process unknown; LOCATED IN: endomembrane system; Ha. [Source:TAIR;Acc:AT2G32487] |
| AT2G33050 | AtRLP26 | -2.47 | -1.15 | 0.023 | 0.012 | Receptor like protein 26 [Source:UniProtKB/Swiss-Prot;Acc:O49328] |
| AT2G33170 | - | -1.56 | -1.58 | 0.010 | 0.005 | Probable leucine-rich repeat receptor-like protein kinase At2g33170 [Source:UniProtKB/Swiss-Prot;Acc:O49318] |
| AT2G33210 | HSP60-2 | -1.44 | -1.22 | 0.007 | 0.020 | Chaperonin CPN60-like 1, mitochondrial [Source:UniProtKB/Swiss-Prot;Acc:Q8L7B5] |
| AT2G34490 | CYP710A2 | -1.91 | -1.64 | 0.027 | 0.016 | Cytochrome P450 710A2 [Source:UniProtKB/Swiss-Prot;Acc:O64698] |
| AT2G34925 | CLE42 | -1.91 | -1.12 | 0.011 | 0.031 | CLAVATA3/ESR (CLE)-related protein 42 [Source:UniProtKB/Swiss-Prot;Acc:Q6IWB2] |
| AT2G35750 | - | -1.22 | -1.20 | 0.037 | 0.036 | Transmembrane protein [Source:UniProtKB/TrEMBL;Acc:Q9ZQP7] |
| AT2G36880 | METK3 | -1.02 | -1.16 | 0.024 | 0.030 | S-adenosylmethionine synthase [Source:UniProtKB/TrEMBL;Acc:A0A178VXF8] |
| AT2G37600 | RPL36A | -1.26 | -1.10 | 0.004 | 0.032 | 60S ribosomal protein L36-1 [Source:UniProtKB/Swiss-Prot;Acc:O80929] |
| AT2G37690 | - | -2.33 | -1.33 | 0.035 | 0.014 | Phosphoribosylaminoimidazole carboxylase like protein [Source:UniProtKB/TrEMBL;Acc:Q84TI2] |

| | | | | | | |
|---|---|---|---|---|---|---|
| AT2G37770 | AKR4C9 | -1.66 | -1.49 | 0.016 | 0.009 | NADPH-dependent aldo-keto reductase, chloroplastic [Source:UniProtKB/Swiss-Prot;Acc:Q0PGJ6] |
| AT2G37890 | - | -1.27 | -1.76 | 0.027 | 0.031 | Mitochondrial carrier like protein [Source:UniProtKB/TrEMBL;Acc:Q8L7R0] |
| AT2G37990 | - | -1.16 | -1.08 | 0.000 | 0.006 | Ribosome biogenesis regulatory protein homolog [Source:UniProtKB/Swiss-Prot;Acc:Q9SH88] |
| AT2G38110 | GPAT6 | -1.62 | -1.31 | 0.002 | 0.032 | Glycerol-3-phosphate 2-O-acyltransferase 6 [Source:UniProtKB/Swiss-Prot;Acc:O80437] |
| AT2G38760 | ANN3 | -2.03 | -1.51 | 0.029 | 0.035 | Annexin [Source:UniProtKB/TrEMBL;Acc:A0A178W040] |
| AT2G39675 | TAS1C | -2.14 | -1.10 | 0.047 | 0.025 | TAS1C; other RNA [Source:TAIR;Acc:AT2G39675] |
| AT2G39681 | TAS2 | -2.33 | -1.57 | 0.023 | 0.024 | TAS2; other RNA [Source:TAIR;Acc:AT2G39681] |
| AT2G39705 | RTFL8 | -1.89 | -1.01 | 0.000 | 0.005 | RTFL8 [Source:UniProtKB/TrEMBL;Acc:A0A178VQG8] |
| AT2G40230 | - | -3.14 | -1.36 | 0.000 | 0.002 | HXXXD-type acyl-transferase family protein [Source:UniProtKB/TrEMBL;Acc:Q9XEF2] |
| AT2G40360 | BOP1 | -1.93 | -1.62 | 0.009 | 0.002 | Ribosome biogenesis protein BOP1 homolog [Source:UniProtKB/Swiss-Prot;Acc:F4IH25] |
| AT2G40670 | ARR16 | -2.38 | -2.71 | 0.002 | 0.028 | Response regulator 16 [Source:UniProtKB/TrEMBL;Acc:F4II22] |
| AT2G40700 | - | -2.48 | -1.56 | 0.016 | 0.027 | P-loop containing nucleoside triphosphate hydrolases superfamily protein [Source:TAIR;Acc:AT2G40700] |
| AT2G40820 | - | -1.68 | -1.24 | 0.005 | 0.007 | FUNCTIONS IN: molecular_function unknown; INVOLVED IN: biological_process unknown; LOCATED IN: plasma membrane; EXPRESSED IN: 21 plant structures; EXPRESSED DURING: 13 growth stages; BEST Arabidopsis thaliana protein match is: myosin heavy chain-rel /.../TAIR:AT3G56480.1); Ha. [Source:TAIR;Acc:AT2G40820] |
| AT2G41312 | - | -2.65 | -1.27 | 0.015 | 0.034 | other RNA [Source:TAIR;Acc:AT2G41312] |
| AT2G41650 | - | -1.02 | -1.13 | 0.007 | 0.001 | At2g41650 [Source:UniProtKB/TrEMBL;Acc:O22226] |
| AT2G42300 | BHLH48 | -1.40 | -1.09 | 0.003 | 0.022 | Transcription factor bHLH48 [Source:UniProtKB/Swiss-Prot;Acc:Q8VZ02] |
| AT2G42395 | - | -1.01 | -2.29 | 0.015 | 0.028 | At2g42395 [Source:UniProtKB/TrEMBL;Acc:Q84TF7] |
| AT2G42610 | LSH10 | -1.08 | -1.50 | 0.025 | 0.012 | LSH10 [Source:UniProtKB/TrEMBL;Acc:A0A178VLX9] |
| AT2G43110 | - | -2.26 | -2.96 | 0.004 | 0.002 | At2g43110 [Source:UniProtKB/TrEMBL;Acc:Q8GWB4] |
| AT2G43190 | - | -1.59 | -1.39 | 0.049 | 0.011 | POP4 [Source:UniProtKB/TrEMBL;Acc:A0A384KZC1] |
| AT2G43650 | EMB2777 | -2.16 | -1.69 | 0.005 | 0.001 | Sas10/U3 ribonucleoprotein (Utp) family protein [Source:UniProtKB/TrEMBL;Acc:Q8L3P4] |
| AT2G44230 | - | -2.00 | -1.28 | 0.007 | 0.008 | At2g44230/F4I1.4 [Source:UniProtKB/TrEMBL;Acc:O64858] |
| AT2G44670 | FLZ3 | -1.76 | -1.51 | 0.005 | 0.009 | FCS-Like Zinc finger 3 [Source:UniProtKB/Swiss-Prot;Acc:O80506] |
| AT2G44740 | CYCU4-1 | -3.43 | -2.15 | 0.016 | 0.004 | Cyclin-U4-1 [Source:UniProtKB/Swiss-Prot;Acc:O80513] |
| AT2G44860 | - | -1.16 | -1.17 | 0.002 | 0.000 | Probable ribosome biogenesis protein RLP24 [Source:UniProtKB/Swiss-Prot;Acc:O22165] |
| AT2G45290 | TKL-2 | -1.00 | -1.13 | 0.009 | 0.029 | Transketolase-2, chloroplastic [Source:UniProtKB/Swiss-Prot;Acc:F4IW47] |
| AT2G47490 | NDT1 | -1.03 | -1.09 | 0.000 | 0.026 | Nicotinamide adenine dinucleotide transporter 1, chloroplastic [Source:UniProtKB/Swiss-Prot;Acc:O22261] |
| AT2G47800 | ABCC4 | -2.59 | -2.15 | 0.002 | 0.001 | ABC transporter C family member 4 [Source:UniProtKB/Swiss-Prot;Acc:Q7DM58] |
| AT3G01160 | - | -1.53 | -1.13 | 0.026 | 0.048 | Pre-rRNA-processing ESF1-like protein [Source:UniProtKB/TrEMBL;Acc:Q9MAC6] |
| AT3G01345 | - | -8.59 | -8.59 | 0.013 | 0.013 | Expressed protein [Source:UniProtKB/TrEMBL;Acc:Q3EBD7] |
| AT3G02020 | AK3 | -1.28 | -1.53 | 0.005 | 0.002 | Aspartokinase 3, chloroplastic [Source:UniProtKB/Swiss-Prot;Acc:Q9S702] |
| AT3G02650 | - | -1.71 | -1.03 | 0.016 | 0.001 | Pentatricopeptide repeat-containing protein At3g02650, mitochondrial [Source:UniProtKB/Swiss-Prot;Acc:P0C896] |
| AT3G02790 | MBS1 | -1.60 | -2.64 | 0.033 | 0.008 | Protein METHYLENE BLUE SENSITIVITY 1 [Source:UniProtKB/Swiss-Prot;Acc:Q9M8S0] |
| AT3G03920 | - | -1.49 | -1.04 | 0.002 | 0.003 | Putative H/ACA ribonucleoprotein complex subunit 1-like protein 1 [Source:UniProtKB/Swiss-Prot;Acc:Q8VZT0] |
| AT3G04770 | RPSAb | -1.06 | -1.59 | 0.010 | 0.018 | 40S ribosomal protein SA [Source:UniProtKB/TrEMBL;Acc:F4J4W3] |
| AT3G04820 | - | -1.56 | -1.07 | 0.040 | 0.026 | Pseudouridine synthase family protein [Source:UniProtKB/TrEMBL;Acc:Q5XVC2] |
| AT3G05060 | NOP5-2 | -1.83 | -1.58 | 0.001 | 0.001 | Probable nucleolar protein 5-2 [Source:UniProtKB/Swiss-Prot;Acc:Q9MAB3] |

| | | | | | | |
|---|---|---|---|---|---|---|
| AT3G05270 | FPP3 | -1.28 | -1.33 | 0.009 | 0.040 | Filament-like plant protein 3 [Source:UniProtKB/Swiss-Prot;Acc:Q9MA92] |
| AT3G06680 | - | -1.41 | -1.42 | 0.013 | 0.021 | 60S ribosomal protein L29 [Source:UniProtKB/TrEMBL;Acc:F4JC32] |
| AT3G06868 | - | -1.70 | -1.21 | 0.037 | 0.006 | Vitellogenin-like protein [Source:UniProtKB/TrEMBL;Acc:Q0WM46] |
| AT3G06930 | PRMT13 | -1.13 | -1.02 | 0.050 | 0.001 | Probable histone-arginine methyltransferase 1.3 [Source:UniProtKB/Swiss-Prot;Acc:Q84W92] |
| AT3G07050 | NSN1 | -1.83 | -1.17 | 0.033 | 0.008 | Guanine nucleotide-binding protein-like NSN1 [Source:UniProtKB/Swiss-Prot;Acc:Q9M8Z5] |
| AT3G07215 | - | -1.71 | -1.01 | 0.007 | 0.045 | other RNA [Source:TAIR;Acc:AT3G07215] |
| AT3G07590 | SMD1A | -1.27 | -1.18 | 0.002 | 0.010 | Small nuclear ribonucleoprotein SmD1a [Source:UniProtKB/Swiss-Prot;Acc:Q9SSF1] |
| AT3G07750 | - | -1.26 | -1.23 | 0.010 | 0.011 | 3'-5'-exoribonuclease family protein [Source:UniProtKB/TrEMBL;Acc:Q9FPI0] |
| AT3G07860 | SNRNP25 | -2.18 | -1.35 | 0.017 | 0.007 | Ubiquitin-like superfamily protein [Source:UniProtKB/TrEMBL;Acc:A0A178V8D0] |
| AT3G08600 | - | -1.36 | -1.06 | 0.008 | 0.011 | AT3g08600/F17O14_7 [Source:UniProtKB/TrEMBL;Acc:Q9C9Z6] |
| AT3G10530 | - | -2.07 | -1.57 | 0.043 | 0.007 | F18K10.11 protein [Source:UniProtKB/TrEMBL;Acc:Q9LPP3] |
| AT3G10610 | RPS17C | -1.39 | -1.20 | 0.002 | 0.028 | 40S ribosomal protein S17-3 [Source:UniProtKB/Swiss-Prot;Acc:Q9SQZ1] |
| AT3G11670 | DGD1 | -1.95 | -1.38 | 0.009 | 0.026 | DGD1 [Source:UniProtKB/TrEMBL;Acc:A0A178VKL1] |
| AT3G11964 | RRP5 | -2.39 | -2.10 | 0.014 | 0.001 | rRNA biogenesis protein RRP5 [Source:UniProtKB/Swiss-Prot;Acc:F4J8K6] |
| AT3G12110 | ACT11 | -1.71 | -1.05 | 0.002 | 0.049 | Actin-11 [Source:UniProtKB/Swiss-Prot;Acc:P53496] |
| AT3G12860 | - | -2.37 | -1.98 | 0.005 | 0.006 | NOP56-like pre RNA processing ribonucleoprotein [Source:UniProtKB/TrEMBL;Acc:Q9LTV0] |
| AT3G12920 | BRG3 | -2.65 | -1.20 | 0.001 | 0.011 | Probable BOI-related E3 ubiquitin-protein ligase 3 [Source:UniProtKB/Swiss-Prot;Acc:Q9LDD1] |
| AT3G13065 | SRF4 | -1.76 | -2.02 | 0.004 | 0.000 | Protein STRUBBELIG-RECEPTOR FAMILY 4 [Source:UniProtKB/Swiss-Prot;Acc:Q6R2K2] |
| AT3G13230 | - | -1.91 | -1.57 | 0.012 | 0.000 | At3g13230 [Source:UniProtKB/TrEMBL;Acc:Q9LTU6] |
| AT3G13810 | AtIDD11 | -2.31 | -1.43 | 0.028 | 0.002 | indeterminate(ID)-domain 11 [Source:TAIR;Acc:AT3G13810] |
| AT3G14210 | ESM1 | -1.69 | -1.82 | 0.042 | 0.000 | GDSL esterase/lipase ESM1 [Source:UniProtKB/Swiss-Prot;Acc:Q9LJG3] |
| AT3G15030 | TCP4 | -1.49 | -1.43 | 0.005 | 0.006 | TCP4 [Source:UniProtKB/TrEMBL;Acc:A0A384KY84] |
| AT3G15460 | BRIX1-1 | -2.41 | -2.15 | 0.046 | 0.002 | Ribosome biogenesis protein BRX1 homolog 1 [Source:UniProtKB/Swiss-Prot;Acc:Q9LE16] |
| AT3G15510 | NAC056 | -1.40 | -1.20 | 0.008 | 0.010 | NAC transcription factor 56 [Source:UniProtKB/Swiss-Prot;Acc:Q9LD44] |
| AT3G16870 | GATA17 | -2.62 | -1.96 | 0.019 | 0.020 | Uncharacterized protein At3g16870 [Source:UniProtKB/TrEMBL;Acc:Q0WPA1] |
| AT3G17185 | TASIR-ARF | -1.92 | -1.21 | 0.006 | 0.021 | TAS3/TASIR-ARF (TRANS-ACTING SIRNA3); other RNA [Source:TAIR;Acc:AT3G17185] |
| AT3G18130 | RACK1C | -1.53 | -1.61 | 0.001 | 0.033 | Receptor for activated C kinase 1C [Source:UniProtKB/Swiss-Prot;Acc:Q9LV28] |
| AT3G18600 | RH51 | -2.30 | -1.74 | 0.012 | 0.005 | DEAD-box ATP-dependent RNA helicase 51 [Source:UniProtKB/Swiss-Prot;Acc:Q9LIH9] |
| AT3G21420 | - | -1.68 | -1.35 | 0.004 | 0.004 | 2-oxoglutarate (2OG) and Fe(II)-dependent oxygenase superfamily protein [Source:UniProtKB/TrEMBL;Acc:Q9LIF4] |
| AT3G22310 | RH9 | -2.18 | -1.54 | 0.013 | 0.022 | DEAD-box ATP-dependent RNA helicase 9, mitochondrial [Source:UniProtKB/Swiss-Prot;Acc:Q9LUW6] |
| AT3G22660 | EBP2 | -1.54 | -1.31 | 0.001 | 0.000 | Probable rRNA-processing protein EBP2 homolog [Source:UniProtKB/Swiss-Prot;Acc:Q9LUJ5] |
| AT3G23000 | CIPK7 | -1.47 | -1.60 | 0.004 | 0.011 | CBL-interacting serine/threonine-protein kinase 7 [Source:UniProtKB/Swiss-Prot;Acc:Q9XIW0] |
| AT3G23590 | MED33A | -1.57 | -1.35 | 0.002 | 0.006 | Mediator of RNA polymerase II transcription subunit 33A [Source:UniProtKB/Swiss-Prot;Acc:Q9LUG9] |
| AT3G23620 | - | -1.34 | -1.12 | 0.001 | 0.014 | Ribosome production factor 2 homolog [Source:UniProtKB/Swiss-Prot;Acc:Q9LUG5] |
| AT3G23990 | CPN60 | -1.33 | -1.14 | 0.005 | 0.021 | Chaperonin CPN60, mitochondrial [Source:UniProtKB/Swiss-Prot;Acc:P29197] |
| AT3G24000 | PCMP-H87 | -1.38 | -1.40 | 0.034 | 0.039 | Pentatricopeptide repeat-containing protein At3g24000, mitochondrial [Source:UniProtKB/Swiss-Prot;Acc:Q9LIQ7] |
| AT3G24080 | - | -1.41 | -1.31 | 0.012 | 0.000 | KRR1 family protein [Source:UniProtKB/TrEMBL;Acc:F4J5D3] |
| AT3G25290 | - | -1.38 | -1.25 | 0.022 | 0.027 | Cytochrome b561 and DOMON domain-containing protein At3g25290 [Source:UniProtKB/Swiss-Prot;Acc:Q9LSE7] |

| | | | | | | |
|---|---|---|---|---|---|---|
| AT3G25585 | AAPT2 | -1.58 | -1.45 | 0.000 | 0.007 | Choline/ethanolaminephosphotransferase 2 [Source:UniProtKB/Swiss-Prot;Acc:O82568] |
| AT3G25590 | - | -3.21 | -2.73 | 0.020 | 0.042 | At3g25590 [Source:UniProtKB/TrEMBL;Acc:Q9LI86] |
| AT3G26450 | - | -1.50 | -1.03 | 0.004 | 0.036 | Major latex protein, putative [Source:UniProtKB/TrEMBL;Acc:Q9LIN0] |
| AT3G26922 | - | -1.50 | -1.30 | 0.004 | 0.040 | F-box/LRR-repeat protein At3g26922 [Source:UniProtKB/Swiss-Prot;Acc:Q9LJF8] |
| AT3G26932 | DRB3 | -2.02 | -1.62 | 0.012 | 0.015 | Double-stranded RNA-binding protein 3 [Source:UniProtKB/Swiss-Prot;Acc:Q9LJF5] |
| AT3G26960 | - | -2.39 | -1.91 | 0.050 | 0.009 | Pollen Ole e 1 allergen and extensin family protein [Source:UniProtKB/TrEMBL;Acc:F4JEV5] |
| AT3G27325 | - | -1.77 | -1.09 | 0.036 | 0.001 | Hydrolases, acting on ester bond [Source:UniProtKB/TrEMBL;Acc:F4IWG2] |
| AT3G28070 | - | -1.84 | -2.20 | 0.006 | 0.021 | WAT1-related protein At3g28070 [Source:UniProtKB/Swiss-Prot;Acc:Q8VYZ7] |
| AT3G28100 | - | -1.56 | -1.06 | 0.019 | 0.029 | WAT1-related protein At3g28100 [Source:UniProtKB/Swiss-Prot;Acc:Q9LRS5] |
| AT3G28920 | ZHD9 | -1.77 | -1.57 | 0.033 | 0.033 | Zinc-finger homeodomain protein 9 [Source:UniProtKB/Swiss-Prot;Acc:Q9LHF0] |
| AT3G30180 | CYP85A2 | -4.27 | -2.01 | 0.016 | 0.018 | Cytochrome P450 85A2 [Source:UniProtKB/Swiss-Prot;Acc:Q940V4] |
| AT3G43800 | GSTU27 | -1.10 | -1.07 | 0.007 | 0.004 | Glutathione S-transferase U27 [Source:UniProtKB/Swiss-Prot;Acc:Q9LZG7] |
| AT3G44430 | - | -1.16 | -1.11 | 0.028 | 0.008 | Transmembrane protein [Source:UniProtKB/TrEMBL;Acc:Q9M282] |
| AT3G44435 | - | -1.01 | -1.15 | 0.008 | 0.040 | - |
| AT3G44750 | HDT1 | -1.94 | -1.57 | 0.013 | 0.004 | Histone deacetylase HDT1 [Source:UniProtKB/Swiss-Prot;Acc:Q9FVE6] |
| AT3G45030 | RPS20C | -1.11 | -1.01 | 0.020 | 0.044 | 40S ribosomal protein S20-1 [Source:UniProtKB/Swiss-Prot;Acc:P49200] |
| AT3G45230 | - | -1.06 | -1.05 | 0.023 | 0.039 | Hydroxyproline-rich glycoprotein family protein [Source:UniProtKB/TrEMBL;Acc:Q9M1T6] |
| AT3G45680 | NPF2.3 | -1.79 | -2.24 | 0.022 | 0.017 | Protein NRT1/ PTR FAMILY 2.3 [Source:UniProtKB/Swiss-Prot;Acc:Q9M175] |
| AT3G46210 | RRP46 | -1.68 | -1.02 | 0.018 | 0.008 | Exosome complex exonuclease RRP46 homolog [Source:UniProtKB/Swiss-Prot;Acc:Q9LX74] |
| AT3G46990 | - | -1.70 | -1.32 | 0.019 | 0.022 | DUF740 family protein, putative (DUF740) [Source:UniProtKB/TrEMBL;Acc:Q9SD74] |
| AT3G47420 | ATPS3 | -2.15 | -1.37 | 0.013 | 0.038 | Putative glycerol-3-phosphate transporter 1 [Source:UniProtKB/Swiss-Prot;Acc:Q9C5L3] |
| AT3G47750 | ABCA4 | -2.91 | -1.13 | 0.007 | 0.036 | ABC transporter A family member 4 [Source:UniProtKB/Swiss-Prot;Acc:Q9STT8] |
| AT3G48490 | - | -1.18 | -1.80 | 0.037 | 0.009 | At3g48490 [Source:UniProtKB/TrEMBL;Acc:Q8LD28] |
| AT3G48740 | SWEET11 | -1.16 | -1.17 | 0.034 | 0.000 | Bidirectional sugar transporter SWEET11 [Source:UniProtKB/Swiss-Prot;Acc:Q9SMM5] |
| AT3G49660 | WDR5A | -1.51 | -1.45 | 0.013 | 0.020 | WDR5a [Source:UniProtKB/TrEMBL;Acc:A0A178VK59] |
| AT3G49990 | - | -1.17 | -1.03 | 0.009 | 0.001 | At3g49990 [Source:UniProtKB/TrEMBL;Acc:Q9SN19] |
| AT3G50340 | - | -1.52 | -1.02 | 0.001 | 0.018 | At3g50340 [Source:UniProtKB/TrEMBL;Acc:Q9SND3] |
| AT3G50630 | KRP2 | -2.72 | -1.00 | 0.001 | 0.010 | Cyclin-dependent kinase inhibitor 2 [Source:UniProtKB/Swiss-Prot;Acc:Q9SCR2] |
| AT3G51060 | SRS1 | -2.58 | -1.27 | 0.018 | 0.018 | Protein SHI RELATED SEQUENCE 1 [Source:UniProtKB/Swiss-Prot;Acc:Q9SD40] |
| AT3G51950 | - | -1.55 | -1.50 | 0.001 | 0.006 | Zinc finger CCCH domain-containing protein 46 [Source:UniProtKB/Swiss-Prot;Acc:Q9SV09] |
| AT3G52040 | - | -1.10 | -1.06 | 0.017 | 0.029 | 2,3-bisphosphoglycerate-dependent phosphoglycerate mutase [Source:UniProtKB/TrEMBL;Acc:Q9SV00] |
| AT3G55280 | RPL23AB | -1.10 | -1.13 | 0.021 | 0.029 | 60S ribosomal protein L23a-2 [Source:UniProtKB/Swiss-Prot;Acc:Q9M3C3] |
| AT3G55340 | PHIP1 | -2.51 | -1.64 | 0.023 | 0.027 | PHIP1 [Source:UniProtKB/TrEMBL;Acc:A0A384KSV0] |
| AT3G55605 | - | -1.74 | -1.59 | 0.026 | 0.004 | At3g55605 [Source:UniProtKB/TrEMBL;Acc:Q8LCT2] |
| AT3G56070 | CYP19-3 | -1.80 | -1.78 | 0.038 | 0.001 | Peptidyl-prolyl cis-trans isomerase CYP19-3 [Source:UniProtKB/Swiss-Prot;Acc:Q38867] |
| AT3G56370 | IRK | -2.17 | -1.47 | 0.008 | 0.004 | Probable LRR receptor-like serine/threonine-protein kinase IRK [Source:UniProtKB/Swiss-Prot;Acc:Q9LY03] |
| AT3G56990 | EDA7 | -2.78 | -2.66 | 0.011 | 0.035 | AT3g56990/F24I3_70 [Source:UniProtKB/TrEMBL;Acc:Q9M1J9] |
| AT3G57150 | CBF5 | -2.11 | -1.88 | 0.009 | 0.000 | Uncharacterized protein At3g57150 (Fragment) [Source:UniProtKB/TrEMBL;Acc:C0SVF3] |

| | | | | | | |
|---|---|---|---|---|---|---|
| AT3G57490 | RPS2D | -1.06 | -1.21 | 0.020 | 0.011 | 40S ribosomal protein S2-4 [Source:UniProtKB/Swiss-Prot;Acc:Q9SCM3] |
| AT3G57660 | NRPA1 | -3.27 | -2.25 | 0.022 | 0.024 | DNA-directed RNA polymerase I subunit 1 [Source:UniProtKB/Swiss-Prot;Acc:Q9SVY0] |
| AT3G57940 | - | -1.92 | -1.97 | 0.012 | 0.017 | RNA cytidine acetyltransferase 2 [Source:UniProtKB/Swiss-Prot;Acc:Q9M2Q4] |
| AT3G58660 | - | -1.48 | -1.43 | 0.023 | 0.008 | Ribosomal protein L1p/L10e family [Source:UniProtKB/TrEMBL;Acc:Q9LXT5] |
| AT3G58990 | IPMI1 | -1.66 | -1.81 | 0.048 | 0.000 | 3-isopropylmalate dehydratase small subunit 2 [Source:UniProtKB/Swiss-Prot;Acc:Q9LYT7] |
| AT3G59630 | - | -2.49 | -1.26 | 0.001 | 0.022 | 2-(3-amino-3-carboxypropyl)histidine synthase subunit 2 [Source:UniProtKB/TrEMBL;Acc:Q9M1A5] |
| AT3G59670 | - | -2.70 | -1.93 | 0.009 | 0.006 | Elongation factor [Source:UniProtKB/TrEMBL;Acc:Q56XZ5] |
| AT3G60400 | MTERF18 | -1.18 | -1.04 | 0.011 | 0.020 | Transcription termination factor MTEF18, mitochondrial [Source:UniProtKB/Swiss-Prot;Acc:Q9M219] |
| AT3G60440 | - | -1.05 | -1.21 | 0.005 | 0.012 | Phosphoglycerate mutase family protein [Source:UniProtKB/TrEMBL;Acc:F4JBU0] |
| AT3G61100 | - | -1.87 | -1.53 | 0.002 | 0.010 | Putative endonuclease or glycosyl hydrolase [Source:UniProtKB/TrEMBL;Acc:Q9LEW6] |
| AT3G61780 | emb1703 | -1.50 | -1.10 | 0.024 | 0.031 | Embryo defective 1703 [Source:UniProtKB/TrEMBL;Acc:Q9M360] |
| AT3G62050 | - | -1.43 | -1.50 | 0.004 | 0.049 | At3g62050 [Source:UniProtKB/TrEMBL;Acc:A6QR85] |
| AT3G63130 | RANGAP1 | -1.32 | -1.11 | 0.002 | 0.020 | RAN GTPase-activating protein 1 [Source:UniProtKB/Swiss-Prot;Acc:Q9LE82] |
| AT3G66652 | FIPS3 | -2.48 | -1.74 | 0.034 | 0.010 | FIP1[III]-like protein [Source:UniProtKB/Swiss-Prot;Acc:F4JC20] |
| AT4G00310 | MEE46 | -1.40 | -1.04 | 0.033 | 0.041 | AT4G00310 protein [Source:UniProtKB/TrEMBL;Acc:O23071] |
| AT4G00930 | CIP4.1 | -1.18 | -1.22 | 0.032 | 0.002 | COP1-interacting protein 4.1 [Source:UniProtKB/TrEMBL;Acc:F4JHQ0] |
| AT4G00950 | MEE47 | -2.45 | -1.42 | 0.013 | 0.009 | Uncharacterized protein At4g00950 [Source:UniProtKB/Swiss-Prot;Acc:Q9M160] |
| AT4G01070 | UGT72B1 | -1.92 | -1.59 | 0.017 | 0.038 | UDP-glycosyltransferase 72B1 [Source:UniProtKB/Swiss-Prot;Acc:Q9M156] |
| AT4G01950 | ATGPAT3 | -1.90 | -1.85 | 0.008 | 0.015 | glycerol-3-phosphate acyltransferase 3 [Source:TAIR;Acc:AT4G01950] |
| AT4G01990 | - | -1.78 | -1.43 | 0.018 | 0.045 | Pentatricopeptide repeat-containing protein At4g01990, mitochondrial [Source:UniProtKB/Swiss-Prot;Acc:Q93WC5] |
| AT4G02075 | PIT1 | -1.33 | -1.25 | 0.022 | 0.033 | At4g02075 [Source:UniProtKB/TrEMBL;Acc:Q9XF50] |
| AT4G02630 | - | -1.55 | -1.49 | 0.001 | 0.046 | Protein kinase superfamily protein [Source:UniProtKB/TrEMBL;Acc:O22764] |
| AT4G02930 | TUFA | -1.39 | -1.17 | 0.034 | 0.004 | Elongation factor Tu, mitochondrial [Source:UniProtKB/Swiss-Prot;Acc:Q9ZT91] |
| AT4G03010 | - | -1.79 | -1.72 | 0.008 | 0.015 | Leucine-rich repeat family protein [Source:UniProtKB/TrEMBL;Acc:Q9ZT98] |
| AT4G04670 | - | -3.31 | -1.61 | 0.010 | 0.036 | tRNA wybutosine-synthesizing protein 2/3/4 [Source:UniProtKB/Swiss-Prot;Acc:Q8W4K1] |
| AT4G07825 | - | -1.98 | -1.21 | 0.006 | 0.014 | At4g07825 [Source:UniProtKB/TrEMBL;Acc:Q8GZ72] |
| AT4G09890 | - | -2.21 | -1.69 | 0.003 | 0.018 | At4g09890 [Source:UniProtKB/TrEMBL;Acc:Q9T0E8] |
| AT4G09970 | - | -2.03 | -1.49 | 0.011 | 0.027 | At4g09970 [Source:UniProtKB/TrEMBL;Acc:Q6DBH2] |
| AT4G10450 | RPL9D | -1.55 | -1.20 | 0.001 | 0.007 | 60S ribosomal protein L9-2 [Source:UniProtKB/Swiss-Prot;Acc:Q9SZX9] |
| AT4G10550 | - | -1.53 | -1.41 | 0.037 | 0.045 | Subtilase family protein [Source:TAIR;Acc:AT4G10550] |
| AT4G10620 | - | -2.62 | -1.29 | 0.048 | 0.034 | F3H7.11 protein [Source:UniProtKB/TrEMBL;Acc:Q9ZSB8] |
| AT4G12030 | BASS5 | -1.26 | -1.50 | 0.022 | 0.000 | Probable sodium/metabolite cotransporter BASS5, chloroplastic [Source:UniProtKB/Swiss-Prot;Acc:F4JPW1] |
| AT4G12430 | TPPF | -2.08 | -1.38 | 0.001 | 0.003 | Probable trehalose-phosphate phosphatase F [Source:UniProtKB/Swiss-Prot;Acc:Q9SU39] |
| AT4G12600 | - | -1.83 | -1.35 | 0.002 | 0.033 | Ribosomal protein L7Ae/L30e/S12e/Gadd45 family protein [Source:UniProtKB/TrEMBL;Acc:F4JRD3] |
| AT4G13575 | - | -2.62 | -2.03 | 0.020 | 0.012 | unknown protein; FUNCTIONS IN: molecular_function unknown; INVOLVED IN: biological_process unknown; LOCATED IN: cellular_component unknown; Ha. [Source:TAIR;Acc:AT4G13575] |
| AT4G13840 | CER26 | -1.76 | -1.26 | 0.011 | 0.044 | Protein ECERIFERUM 26 [Source:UniProtKB/Swiss-Prot;Acc:Q9SVM9] |
| AT4G14740 | - | -2.10 | -1.21 | 0.007 | 0.001 | Auxin canalization protein (DUF828) [Source:UniProtKB/TrEMBL;Acc:Q8VY20] |
| AT4G14750 | IQD19 | -2.27 | -1.32 | 0.017 | 0.030 | IQ-domain 19 [Source:TAIR;Acc:AT4G14750] |

| | | | | | | |
|---|---|---|---|---|---|---|
| AT4G14760 | NET1B | -1.70 | -1.24 | 0.020 | 0.016 | Protein NETWORKED 1B [Source:UniProtKB/Swiss-Prot;Acc:F4JIF4] |
| AT4G15440 | CYP74B2 | -1.97 | -1.14 | 0.046 | 0.007 | Probable inactive linolenate hydroperoxide lyase [Source:UniProtKB/Swiss-Prot;Acc:B3LF83] |
| AT4G15770 | - | -1.68 | -1.31 | 0.015 | 0.000 | 60S ribosome subunit biogenesis protein NIP7 homolog [Source:UniProtKB/TrEMBL;Acc:Q6NM52] |
| AT4G16141 | - | -1.80 | -1.07 | 0.000 | 0.000 | GATA type zinc finger transcription factor family protein [Source:UniProtKB/TrEMBL;Acc:Q8GW81] |
| AT4G16610 | - | -8.30 | -1.31 | 0.047 | 0.047 | C2H2-like zinc finger protein [Source:UniProtKB/TrEMBL;Acc:O23504] |
| AT4G16630 | RH28 | -1.40 | -1.22 | 0.018 | 0.001 | DEAD-box ATP-dependent RNA helicase 28 [Source:UniProtKB/Swiss-Prot;Acc:Q9ZRZ8] |
| AT4G16670 | - | -2.45 | -1.36 | 0.034 | 0.030 | At4g16670 [Source:UniProtKB/TrEMBL;Acc:Q5HZ31] |
| AT4G17770 | TPS5 | -1.11 | -1.11 | 0.039 | 0.026 | Alpha,alpha-trehalose-phosphate synthase [UDP-forming] 5 [Source:UniProtKB/Swiss-Prot;Acc:O23617] |
| AT4G17880 | MYC4 | -2.31 | -1.47 | 0.004 | 0.037 | Transcription factor MYC4 [Source:UniProtKB/Swiss-Prot;Acc:O49687] |
| AT4G18570 | - | -2.02 | -1.59 | 0.003 | 0.002 | AT4g18560/F28J12_220 [Source:UniProtKB/TrEMBL;Acc:Q8L7S5] |
| AT4G21140 | - | -1.38 | -1.22 | 0.027 | 0.008 | BEST Arabidopsis thaliana protein match is: copper ion binding (TAIR:AT4G05400.2); Ha. [Source:TAIR;Acc:AT4G21140] |
| AT4G21220 | LPXD2 | -2.15 | -1.57 | 0.013 | 0.032 | Probable UDP-3-O-acylglucosamine N-acyltransferase 2, mitochondrial [Source:UniProtKB/Swiss-Prot;Acc:F4JIP6] |
| AT4G22380 | - | -1.40 | -1.08 | 0.008 | 0.021 | At4g22380 [Source:UniProtKB/TrEMBL;Acc:Q8LCC7] |
| AT4G24710 | - | -1.64 | -1.36 | 0.022 | 0.009 | P-loop containing nucleoside triphosphate hydrolases superfamily protein [Source:TAIR;Acc:AT4G24710] |
| AT4G25340 | FKBP53 | -1.83 | -1.77 | 0.003 | 0.000 | Peptidyl-prolyl cis-trans isomerase FKBP53 [Source:UniProtKB/Swiss-Prot;Acc:Q93ZG9] |
| AT4G25730 | - | -2.02 | -1.68 | 0.000 | 0.000 | Putative rRNA methyltransferase [Source:UniProtKB/TrEMBL;Acc:F4JTD2] |
| AT4G26540 | RGI3 | -1.97 | -1.58 | 0.025 | 0.044 | LRR receptor-like serine/threonine-protein kinase [Source:UniProtKB/Swiss-Prot;Acc:C0LGR3] |
| AT4G26960 | - | -1.67 | -1.22 | 0.021 | 0.012 | unknown protein; FUNCTIONS IN: molecular_function unknown; INVOLVED IN: biological_process unknown; LOCATED IN: chloroplast; BEST Arabidopsis thaliana protein match is: unknown protein (TAIR:AT5G54970.1); Ha. [Source:TAIR;Acc:AT4G26960] |
| AT4G27340 | - | -1.71 | -1.17 | 0.045 | 0.008 | tRNA (guanine(37)-N1)-methyltransferase 2 [Source:UniProtKB/Swiss-Prot;Acc:Q6NQ64] |
| AT4G28010 | - | -2.89 | -2.02 | 0.005 | 0.012 | RPF5 [Source:UniProtKB/TrEMBL;Acc:A0A178UVK0] |
| AT4G28450 | - | -1.69 | -1.48 | 0.003 | 0.009 | AT4g28450/F20O9_130 [Source:UniProtKB/TrEMBL;Acc:Q93VK1] |
| AT4G30020 | SBT2.6 | -1.00 | -1.42 | 0.010 | 0.019 | Subtilisin-like protease SBT2.6 [Source:UniProtKB/Swiss-Prot;Acc:Q9SZV5] |
| AT4G30400 | ATL13 | -4.15 | -2.40 | 0.001 | 0.003 | RING-H2 finger protein ATL13 [Source:UniProtKB/Swiss-Prot;Acc:Q940Q4] |
| AT4G30800 | RPS11B | -1.44 | -1.46 | 0.031 | 0.004 | 40S ribosomal protein S11-2 [Source:UniProtKB/Swiss-Prot;Acc:O65569] |
| AT4G30960 | CIPK6 | -1.37 | -1.05 | 0.011 | 0.026 | CBL-interacting serine/threonine-protein kinase 6 [Source:UniProtKB/Swiss-Prot;Acc:O65554] |
| AT4G31330 | - | -2.55 | -1.59 | 0.012 | 0.046 | Uncharacterized protein At4g31330 [Source:UniProtKB/TrEMBL;Acc:Q9C5C1] |
| AT4G31590 | CSLC5 | -1.95 | -1.45 | 0.000 | 0.037 | Glycosyltransferase (Fragment) [Source:UniProtKB/TrEMBL;Acc:W8PV45] |
| AT4G32920 | - | -1.05 | -1.87 | 0.015 | 0.002 | Glycine-rich protein [Source:UniProtKB/TrEMBL;Acc:F4JV81] |
| AT4G33040 | GRXC6 | -1.53 | -1.77 | 0.007 | 0.032 | Glutaredoxin-C6 [Source:UniProtKB/Swiss-Prot;Acc:Q8L9S3] |
| AT4G34555 | RPS25D | -1.14 | -1.17 | 0.037 | 0.048 | 40S ribosomal protein S25-3 [Source:UniProtKB/Swiss-Prot;Acc:Q8GYL5] |
| AT4G34650 | SQS2 | -1.69 | -1.08 | 0.007 | 0.024 | Inactive squalene synthase 2 [Source:UniProtKB/Swiss-Prot;Acc:O65688] |
| AT4G36240 | GATA7 | -1.95 | -1.39 | 0.040 | 0.005 | GATA transcription factor 7 [Source:UniProtKB/Swiss-Prot;Acc:O65515] |
| AT4G36380 | CYP90C1 | -2.43 | -1.12 | 0.017 | 0.000 | ROT3 [Source:UniProtKB/TrEMBL;Acc:A0A178V4B0] |
| AT4G37080 | - | -2.31 | -1.53 | 0.006 | 0.020 | Protein of unknown function, DUF547 [Source:TAIR;Acc:AT4G37080] |
| AT4G38100 | CURT1D | -1.04 | -1.10 | 0.016 | 0.013 | Protein CURVATURE THYLAKOID 1D, chloroplastic [Source:UniProtKB/Swiss-Prot;Acc:Q8LDD3] |
| AT4G38150 | - | -1.97 | -1.30 | 0.015 | 0.006 | Pentatricopeptide repeat-containing protein At4g38150 [Source:UniProtKB/Swiss-Prot;Acc:Q9SZL5] |
| AT4G38950 | KIN7F | -1.72 | -1.08 | 0.019 | 0.025 | Kinesin-like protein KIN-7F [Source:UniProtKB/Swiss-Prot;Acc:F4JUI9] |
| AT4G39070 | BBX20 | -1.44 | -1.83 | 0.018 | 0.018 | B-box zinc finger protein 20 [Source:UniProtKB/Swiss-Prot;Acc:Q0IGM7] |

| | | | | | | |
|---|---|---|---|---|---|---|
| AT4G39510 | CYP96A12 | -1.66 | -1.02 | 0.006 | 0.008 | CYP96A12 [Source:UniProtKB/TrEMBL;Acc:A0A178V036] |
| AT4G39740 | HCC2 | -1.64 | -1.00 | 0.001 | 0.048 | HCC2 [Source:UniProtKB/TrEMBL;Acc:A0A178V000] |
| AT4G39770 | TPPH | -2.36 | -1.90 | 0.005 | 0.027 | Probable trehalose-phosphate phosphatase H [Source:UniProtKB/Swiss-Prot;Acc:Q8GWG2] |
| AT4G39800 | IPS1 | -1.18 | -1.63 | 0.006 | 0.042 | Inositol-3-phosphate synthase isozyme 1 [Source:UniProtKB/Swiss-Prot;Acc:P42801] |
| AT4G40060 | ATHB-16 | -1.57 | -1.37 | 0.016 | 0.009 | HB16 [Source:UniProtKB/TrEMBL;Acc:A0A178V0P3] |
| AT5G01075 | - | -1.79 | -1.16 | 0.002 | 0.011 | At5g01075 [Source:UniProtKB/TrEMBL;Acc:Q8LEW6] |
| AT5G01170 | OPSL1 | -1.37 | -2.06 | 0.006 | 0.002 | Protein OCTOPUS-like [Source:UniProtKB/Swiss-Prot;Acc:Q9LFB9] |
| AT5G01740 | - | -3.62 | -1.70 | 0.000 | 0.017 | AT5g01740/T20L15_10 [Source:UniProtKB/TrEMBL;Acc:Q9LZX2] |
| AT5G01810 | CIPK15 | -1.25 | -1.16 | 0.033 | 0.003 | CBL-interacting serine/threonine-protein kinase 15 [Source:UniProtKB/Swiss-Prot;Acc:P92937] |
| AT5G02050 | - | -1.51 | -1.24 | 0.000 | 0.014 | At5g02050 [Source:UniProtKB/TrEMBL;Acc:Q9LZM6] |
| AT5G02760 | - | -3.01 | -1.60 | 0.022 | 0.001 | Probable protein phosphatase 2C 67 [Source:UniProtKB/Swiss-Prot;Acc:Q501F9] |
| AT5G03120 | - | -3.16 | -1.08 | 0.005 | 0.004 | unknown protein; FUNCTIONS IN: molecular_function unknown; INVOLVED IN: biological_process unknown; LOCATED IN: endomembrane system; EXPRESSED IN: 21 plant structures; EXPRESSED DURING: 13 growth stages; Ha. [Source:TAIR;Acc:AT5G03120] |
| AT5G03140 | LECRK82 | -2.23 | -1.98 | 0.020 | 0.001 | L-type lectin-domain containing receptor kinase VIII.2 [Source:UniProtKB/Swiss-Prot;Acc:Q9LYX1] |
| AT5G03760 | CSLA9 | -2.37 | -1.34 | 0.002 | 0.040 | Glucomannan 4-beta-mannosyltransferase 9 [Source:UniProtKB/Swiss-Prot;Acc:Q9LZR3] |
| AT5G04190 | PKS4 | -1.99 | -1.35 | 0.001 | 0.015 | Protein PHYTOCHROME KINASE SUBSTRATE 4 [Source:UniProtKB/Swiss-Prot;Acc:Q9FYE2] |
| AT5G04530 | KCS19 | -1.95 | -2.08 | 0.007 | 0.019 | 3-ketoacyl-CoA synthase 19 [Source:UniProtKB/Swiss-Prot;Acc:Q9LZ72] |
| AT5G04950 | NAS1 | -2.06 | -2.25 | 0.029 | 0.017 | Nicotianamine synthase 1 [Source:UniProtKB/Swiss-Prot;Acc:Q9FF79] |
| AT5G05680 | NUP88 | -1.01 | -1.18 | 0.001 | 0.039 | Nuclear pore complex protein NUP88 [Source:UniProtKB/Swiss-Prot;Acc:Q9FFK6] |
| AT5G05860 | UGT76C2 | -1.31 | -1.37 | 0.016 | 0.025 | Glycosyltransferase (Fragment) [Source:UniProtKB/TrEMBL;Acc:W8QNB5] |
| AT5G05990 | - | -1.74 | -1.31 | 0.032 | 0.000 | AT5g05990/K18J17_19 [Source:UniProtKB/TrEMBL;Acc:Q9FI87] |
| AT5G06270 | - | -2.71 | -1.39 | 0.000 | 0.030 | Gb [Source:UniProtKB/TrEMBL;Acc:Q9FNI1] |
| AT5G06530 | ABCG22 | -2.68 | -1.18 | 0.001 | 0.032 | ABC transporter G family member 22 [Source:UniProtKB/Swiss-Prot;Acc:Q93YS4] |
| AT5G07000 | SOT14 | -2.01 | -1.06 | 0.001 | 0.047 | Cytosolic sulfotransferase 14 [Source:UniProtKB/Swiss-Prot;Acc:Q8GZ53] |
| AT5G07240 | IQD24 | -2.25 | -1.31 | 0.000 | 0.017 | IQ-domain 24 [Source:TAIR;Acc:AT5G07240] |
| AT5G07580 | ERF106 | -1.41 | -1.43 | 0.013 | 0.005 | Ethylene-responsive transcription factor ERF106 [Source:UniProtKB/Swiss-Prot;Acc:Q9LY05] |
| AT5G07690 | MYB29 | -1.26 | -1.71 | 0.000 | 0.008 | Transcription factor MYB29 [Source:UniProtKB/Swiss-Prot;Acc:Q9FLR1] |
| AT5G08180 | - | -1.98 | -1.76 | 0.000 | 0.000 | H/ACA ribonucleoprotein complex subunit 2-like protein [Source:UniProtKB/Swiss-Prot;Acc:Q9LEY9] |
| AT5G08260 | SCPL35 | -1.61 | -1.52 | 0.049 | 0.007 | Serine carboxypeptidase-like 35 [Source:UniProtKB/Swiss-Prot;Acc:Q9LEY1] |
| AT5G08620 | RH25 | -2.45 | -1.54 | 0.011 | 0.007 | DEAD-box ATP-dependent RNA helicase 25 [Source:UniProtKB/Swiss-Prot;Acc:Q94C75] |
| AT5G10965 | - | -1.87 | -1.21 | 0.034 | 0.003 | - |
| AT5G11240 | - | -1.98 | -1.35 | 0.000 | 0.035 | At5g11240 [Source:UniProtKB/TrEMBL;Acc:B5X503] |
| AT5G11550 | - | -1.31 | -1.33 | 0.006 | 0.007 | ARM repeat superfamily protein [Source:UniProtKB/TrEMBL;Acc:Q9LYD7] |
| AT5G11790 | NDL2 | -1.38 | -1.31 | 0.003 | 0.030 | Protein NDL2 [Source:UniProtKB/Swiss-Prot;Acc:Q9ASU8] |
| AT5G12220 | - | -1.65 | -1.29 | 0.020 | 0.018 | Las1-like family protein [Source:UniProtKB/TrEMBL;Acc:F4JZH9] |
| AT5G12420 | - | -1.59 | -1.10 | 0.003 | 0.028 | O-acyltransferase (WSD1-like) family protein [Source:UniProtKB/TrEMBL;Acc:Q94CK0] |
| AT5G13400 | NPF6.1 | -2.44 | -1.04 | 0.000 | 0.002 | Protein NRT1/ PTR FAMILY 6.1 [Source:UniProtKB/Swiss-Prot;Acc:Q9LYR6] |
| AT5G14050 | - | -1.62 | -1.23 | 0.004 | 0.006 | U3 small nucleolar RNA-associated protein 18 homolog [Source:UniProtKB/Swiss-Prot;Acc:Q9FMU5] |
| AT5G14210 | - | -1.36 | -1.03 | 0.001 | 0.038 | Leucine-rich repeat protein kinase family protein [Source:TAIR;Acc:AT5G14210] |

| | | | | | | |
|---|---|---|---|---|---|---|
| AT5G14520 | PES | -1.56 | -1.01 | 0.014 | 0.004 | Pescadillo homolog [Source:UniProtKB/Swiss-Prot;Acc:Q9LYK7] |
| AT5G14800 | PROC1 | -1.07 | -1.09 | 0.024 | 0.009 | Pyrroline-5-carboxylate reductase [Source:UniProtKB/Swiss-Prot;Acc:P54904] |
| AT5G14880 | POT8 | -1.74 | -1.07 | 0.001 | 0.013 | Potassium transporter [Source:UniProtKB/TrEMBL;Acc:A0A178UBD9] |
| AT5G15700 | - | -1.97 | -1.38 | 0.001 | 0.035 | DNA-directed RNA polymerase [Source:UniProtKB/TrEMBL;Acc:F4KB75] |
| AT5G15740 | OFUT34 | -2.07 | -1.68 | 0.023 | 0.008 | O-fucosyltransferase 34 [Source:UniProtKB/Swiss-Prot;Acc:Q4V398] |
| AT5G15750 | - | -1.28 | -1.33 | 0.019 | 0.037 | Alpha-L RNA-binding motif/Ribosomal protein S4 family protein [Source:UniProtKB/TrEMBL;Acc:Q683D4] |
| AT5G15980 | - | -1.13 | -1.08 | 0.038 | 0.019 | Pentatricopeptide repeat-containing protein At5g15980, mitochondrial [Source:UniProtKB/Swiss-Prot;Acc:Q8LPF1] |
| AT5G16000 | NIK1 | -2.03 | -1.62 | 0.007 | 0.000 | Protein NSP-INTERACTING KINASE 1 [Source:UniProtKB/Swiss-Prot;Acc:Q9LFS4] |
| AT5G17580 | - | -2.01 | -1.04 | 0.004 | 0.014 | Phototropic-responsive NPH3 family protein [Source:TAIR;Acc:AT5G17580] |
| AT5G17790 | VAR3 | -1.05 | -1.02 | 0.050 | 0.031 | Zinc finger protein VAR3, chloroplastic [Source:UniProtKB/Swiss-Prot;Acc:Q8S9K3] |
| AT5G18440 | - | -1.73 | -1.06 | 0.030 | 0.045 | NUFIP [Source:UniProtKB/TrEMBL;Acc:A0A178UB40] |
| AT5G18770 | - | -1.30 | -1.09 | 0.033 | 0.046 | F-box/FBD/LRR-repeat protein At5g18770 [Source:UniProtKB/Swiss-Prot;Acc:Q93ZK9] |
| AT5G20160 | - | -1.37 | -1.10 | 0.001 | 0.001 | Ribosomal protein L7Ae/L30e/S12e/Gadd45 family protein [Source:UniProtKB/TrEMBL;Acc:F4K455] |
| AT5G20270 | HHP1 | -2.06 | -1.51 | 0.005 | 0.006 | Heptahelical transmembrane protein 1 [Source:UniProtKB/Swiss-Prot;Acc:Q93ZH9] |
| AT5G20700 | FLZ14 | -1.46 | -1.35 | 0.004 | 0.002 | FCS-Like Zinc finger 14 [Source:UniProtKB/Swiss-Prot;Acc:Q8GYX2] |
| AT5G21100 | - | -1.51 | -1.94 | 0.000 | 0.039 | Ascorbate oxidase-like protein [Source:UniProtKB/TrEMBL;Acc:Q8LPL3] |
| AT5G21482 | CKX7 | -1.09 | -1.10 | 0.000 | 0.001 | Cytokinin dehydrogenase 7 [Source:UniProtKB/Swiss-Prot;Acc:Q9FUJ1] |
| AT5G21960 | ERF016 | -1.54 | -3.07 | 0.043 | 0.019 | Ethylene-responsive transcription factor ERF016 [Source:UniProtKB/Swiss-Prot;Acc:Q9C591] |
| AT5G22020 | - | -2.02 | -1.53 | 0.000 | 0.004 | At5g22020 [Source:UniProtKB/TrEMBL;Acc:Q9C586] |
| AT5G22460 | - | -1.90 | -2.25 | 0.003 | 0.009 | Alpha/beta-Hydrolases superfamily protein [Source:UniProtKB/TrEMBL;Acc:Q9FMQ5] |
| AT5G22650 | HDT2 | -1.17 | -1.03 | 0.003 | 0.002 | Histone deacetylase HDT2 [Source:UniProtKB/Swiss-Prot;Acc:Q56WH4] |
| AT5G23100 | - | -1.57 | -1.29 | 0.004 | 0.025 | At5g23100 [Source:UniProtKB/TrEMBL;Acc:Q9FN45] |
| AT5G23280 | TCP7 | -3.32 | -1.48 | 0.005 | 0.001 | Transcription factor TCP7 [Source:UniProtKB/Swiss-Prot;Acc:Q9FMX2] |
| AT5G24660 | LSU2 | -2.59 | -2.12 | 0.049 | 0.025 | Protein RESPONSE TO LOW SULFUR 2 [Source:UniProtKB/Swiss-Prot;Acc:Q9FIR9] |
| AT5G25220 | KNAT3 | -1.45 | -1.01 | 0.004 | 0.001 | KNAT3 [Source:UniProtKB/TrEMBL;Acc:A0A178U9D1] |
| AT5G27120 | NOP5-1 | -2.27 | -1.60 | 0.018 | 0.026 | Probable nucleolar protein 5-1 [Source:UniProtKB/Swiss-Prot;Acc:O04658] |
| AT5G35480 | - | -1.11 | -1.05 | 0.034 | 0.029 | At5g35480 [Source:UniProtKB/TrEMBL;Acc:Q9FJB1] |
| AT5G38040 | UGT76E7 | -1.93 | -2.05 | 0.018 | 0.026 | Glycosyltransferase (Fragment) [Source:UniProtKB/TrEMBL;Acc:W8PUC5] |
| AT5G38720 | - | -1.80 | -1.21 | 0.000 | 0.025 | unknown protein; Ha. [Source:TAIR;Acc:AT5G38720] |
| AT5G39850 | RPS9C | -1.44 | -1.64 | 0.029 | 0.011 | 40S ribosomal protein S9-2 [Source:UniProtKB/Swiss-Prot;Acc:Q9FLF0] |
| AT5G40380 | CRK42 | -4.04 | -2.13 | 0.012 | 0.007 | Cysteine-rich receptor-like protein kinase 42 [Source:UniProtKB/Swiss-Prot;Acc:Q9FNE1] |
| AT5G40490 | - | -1.18 | -1.01 | 0.001 | 0.001 | RNA-binding (RRM/RBD/RNP motifs) family protein [Source:UniProtKB/TrEMBL;Acc:Q9FM47] |
| AT5G42860 | - | -2.06 | -2.12 | 0.018 | 0.001 | Emb [Source:UniProtKB/TrEMBL;Acc:Q9FMN3] |
| AT5G43310 | - | -1.28 | -1.14 | 0.017 | 0.021 | COP1-interacting protein-like protein [Source:UniProtKB/TrEMBL;Acc:F4K5Y5] |
| AT5G44680 | - | -2.20 | -1.23 | 0.006 | 0.009 | DNA glycosylase superfamily protein [Source:UniProtKB/TrEMBL;Acc:Q9FIZ5] |
| AT5G46740 | UBP21 | -1.46 | -2.14 | 0.035 | 0.049 | Ubiquitin carboxyl-terminal hydrolase 21 [Source:UniProtKB/Swiss-Prot;Acc:Q9FIQ1] |
| AT5G48800 | - | -1.24 | -1.53 | 0.045 | 0.002 | BTB/POZ domain-containing protein At5g48800 [Source:UniProtKB/Swiss-Prot;Acc:Q9FKB6] |
| AT5G48850 | SDI1 | -3.91 | -2.51 | 0.009 | 0.021 | ATSDI1 [Source:UniProtKB/TrEMBL;Acc:A0A178UKA2] |

| | | | | | | |
|---|---|---|---|---|---|---|
| AT5G48930 | HST | -1.28 | -1.15 | 0.012 | 0.005 | Shikimate O-hydroxycinnamoyltransferase [Source:UniProtKB/Swiss-Prot;Acc:Q9FI78] |
| AT5G49215 | - | -1.67 | -1.61 | 0.025 | 0.003 | Pectin lyase-like superfamily protein [Source:UniProtKB/TrEMBL;Acc:Q8RX04] |
| AT5G50810 | TIM8 | -1.32 | -1.11 | 0.001 | 0.000 | Mitochondrial import inner membrane translocase subunit TIM8 [Source:UniProtKB/Swiss-Prot;Acc:Q9XGY4] |
| AT5G52470 | MED36B | -1.49 | -1.01 | 0.000 | 0.040 | Probable mediator of RNA polymerase II transcription subunit 36b [Source:UniProtKB/Swiss-Prot;Acc:Q9FEF8] |
| AT5G53370 | ATPMEPCRF | -1.43 | -1.19 | 0.029 | 0.016 | pectin methylesterase PCR fragment F [Source:TAIR;Acc:AT5G53370] |
| AT5G53880 | - | -1.71 | -1.21 | 0.002 | 0.010 | Putative uncharacterized protein [Source:UniProtKB/TrEMBL;Acc:Q9FN38] |
| AT5G53900 | - | -1.59 | -1.06 | 0.003 | 0.048 | Gb [Source:UniProtKB/TrEMBL;Acc:Q9FN36] |
| AT5G54250 | CNGC4 | -1.97 | -1.59 | 0.000 | 0.010 | Cyclic nucleotide-gated ion channel 4 [Source:UniProtKB/Swiss-Prot;Acc:Q94AS9] |
| AT5G55400 | FIM3 | -1.42 | -1.29 | 0.003 | 0.005 | Fimbrin-3 [Source:UniProtKB/Swiss-Prot;Acc:Q9FJ70] |
| AT5G55920 | OLI2 | -2.03 | -1.64 | 0.020 | 0.004 | Nucleolar protein-like [Source:UniProtKB/TrEMBL;Acc:Q9FG73] |
| AT5G56040 | - | -4.33 | -2.23 | 0.030 | 0.012 | Leucine-rich receptor-like protein kinase family protein [Source:UniProtKB/TrEMBL;Acc:F4K6B8] |
| AT5G57120 | - | -1.20 | -1.14 | 0.018 | 0.008 | AT5g57120/MUL3_6 [Source:UniProtKB/TrEMBL;Acc:Q9LU74] |
| AT5G57280 | RID2 | -1.70 | -1.51 | 0.045 | 0.004 | RID2 [Source:UniProtKB/TrEMBL;Acc:A0A178UNY0] |
| AT5G58570 | - | -1.08 | -1.28 | 0.025 | 0.025 | Transmembrane protein [Source:UniProtKB/TrEMBL;Acc:Q94AT2] |
| AT5G59590 | UGT76E2 | -8.76 | -8.76 | 0.016 | 0.016 | Glycosyltransferase (Fragment) [Source:UniProtKB/TrEMBL;Acc:W8PUA4] |
| AT5G60670 | RPL12C | -1.22 | -1.09 | 0.006 | 0.024 | 60S ribosomal protein L12-3 [Source:UniProtKB/Swiss-Prot;Acc:Q9FF52] |
| AT5G61030 | RBG3 | -1.39 | -1.47 | 0.049 | 0.022 | GR-RBP3 [Source:UniProtKB/TrEMBL;Acc:A0A178UBT5] |
| AT5G61310 | - | -1.81 | -1.88 | 0.010 | 0.014 | Cytochrome c oxidase subunit 5C [Source:UniProtKB/TrEMBL;Acc:A0A178UCG7] |
| AT5G61340 | - | -1.77 | -1.54 | 0.001 | 0.001 | Transmembrane protein [Source:UniProtKB/TrEMBL;Acc:Q9FLJ9] |
| AT5G61420 | MYB28 | -1.87 | -1.34 | 0.004 | 0.003 | PMG1 [Source:UniProtKB/TrEMBL;Acc:A0A178UN88] |
| AT5G61460 | SMC6B | -1.27 | -1.27 | 0.031 | 0.001 | Structural maintenance of chromosomes protein 6B [Source:UniProtKB/Swiss-Prot;Acc:Q9FII7] |
| AT5G61770 | PPAN | -1.19 | -1.26 | 0.008 | 0.004 | PETER PAN-like protein [Source:UniProtKB/TrEMBL;Acc:F4K3M1] |
| AT5G62190 | RH7 | -1.87 | -1.27 | 0.000 | 0.000 | DEAD-box ATP-dependent RNA helicase 7 [Source:UniProtKB/Swiss-Prot;Acc:Q39189] |
| AT5G62370 | - | -5.58 | -1.89 | 0.014 | 0.008 | Pentatricopeptide repeat-containing protein At5g62370 [Source:UniProtKB/Swiss-Prot;Acc:Q9LVA2] |
| AT5G62440 | EMB514 | -1.42 | -1.01 | 0.024 | 0.000 | Protein EMBRYO DEFECTIVE 514 [Source:UniProtKB/Swiss-Prot;Acc:Q8L557] |
| AT5G62920 | ARR6 | -2.56 | -2.44 | 0.016 | 0.007 | Response regulator 6 [Source:UniProtKB/TrEMBL;Acc:Q0WSS6] |
| AT5G63180 | - | -1.59 | -1.19 | 0.005 | 0.023 | Probable pectate lyase 22 [Source:UniProtKB/Swiss-Prot;Acc:Q93Z25] |
| AT5G63530 | HIPP07 | -2.02 | -1.41 | 0.012 | 0.029 | Heavy metal-associated isoprenylated plant protein 7 [Source:UniProtKB/Swiss-Prot;Acc:Q9C5D3] |
| AT5G64420 | - | -1.94 | -1.75 | 0.015 | 0.018 | DNA polymerase V family [Source:UniProtKB/TrEMBL;Acc:Q9FGF4] |
| AT5G65010 | ASN2 | -1.90 | -1.49 | 0.011 | 0.029 | asparagine synthetase 2 [Source:TAIR;Acc:AT5G65010] |
| AT5G65730 | XTH6 | -3.23 | -1.27 | 0.000 | 0.031 | Xyloglucan endotransglucosylase/hydrolase [Source:UniProtKB/TrEMBL;Acc:Q0WUU2] |
| AT5G65860 | - | -2.55 | -1.98 | 0.018 | 0.011 | Ankyrin repeat family protein [Source:UniProtKB/TrEMBL;Acc:Q2V2U9] |
| AT5G65880 | - | -1.03 | -1.30 | 0.021 | 0.017 | At5g65880 [Source:UniProtKB/TrEMBL;Acc:Q9FHP2] |
| AT5G66540 | - | -1.69 | -1.32 | 0.032 | 0.000 | U3 small nucleolar ribonucleoprotein protein MPP10 [Source:UniProtKB/TrEMBL;Acc:Q9FJY5] |
| AT5G66770 | SCL4 | -1.77 | -1.49 | 0.038 | 0.005 | Scarecrow-like protein 4 [Source:UniProtKB/Swiss-Prot;Acc:Q9FL03] |
| AT5G67160 | EPS1 | -2.66 | -2.10 | 0.028 | 0.007 | Protein ENHANCED PSEUDOMONAS SUSCEPTIBILTY 1 [Source:UniProtKB/Swiss-Prot;Acc:Q9FH97] |
| AT5G67200 | - | -3.47 | -1.72 | 0.040 | 0.001 | Probable inactive receptor kinase At5g67200 [Source:UniProtKB/Swiss-Prot;Acc:Q93Y06] |
| AT5G67280 | RLK | -1.97 | -1.55 | 0.018 | 0.047 | At5g67280/K3G17_4 [Source:UniProtKB/TrEMBL;Acc:Q9FGQ5] |
