## Supplemental Table S2 for "Transcriptome and metabolome analyses revealed that narrowband 280 and 310 nm UV-B induce distinctive responses in Arabidopsis"

Common DEGs in 280-2d and 310-2d

| ID | Gene name | $Log_2$ FC | | *P* -value | | Description |
|---|---|---|---|---|---|---|
| | | 280-2d | 310-2d | 280-2d | 310-2d | |
| **'280-2d and 310-2d'_DEGs up-regulated** | | | | | | |
| AT4G37420 | - | 11.37 | 8.67 | 0.000 | 0.007 | Glycosyltransferase family protein (DUF23) [Source:UniProtKB/TrEMBL;Acc:Q9SZU2] |
| AT2G01810 | - | 9.27 | 8.64 | 0.001 | 0.017 | PHD finger protein At2g01810 [Source:UniProtKB/Swiss-Prot;Acc:Q9ZUA9] |
| AT1G10480 | ZFP5 | 9.01 | 8.78 | 0.001 | 0.010 | Zinc finger protein 5 [Source:UniProtKB/Swiss-Prot;Acc:Q39264] |
| AT4G25820 | XTH14 | 8.82 | 7.33 | 0.041 | 0.041 | Xyloglucan endotransglucosylase/hydrolase [Source:UniProtKB/TrEMBL;Acc:A0A178UTG4] |
| AT4G31870 | GPX7 | 5.07 | 3.42 | 0.005 | 0.005 | Putative glutathione peroxidase 7, chloroplastic [Source:UniProtKB/Swiss-Prot;Acc:Q9SZ54] |
| AT2G04515 | - | 4.79 | 2.63 | 0.001 | 0.000 | Transmembrane protein [Source:UniProtKB/TrEMBL;Acc:Q5S4Y8] |
| AT3G11090 | LBD21 | 3.50 | 1.84 | 0.000 | 0.009 | LBD21 [Source:UniProtKB/TrEMBL;Acc:A0A178V6L2] |
| AT2G39410 | - | 3.25 | 1.57 | 0.005 | 0.033 | Alpha/beta-Hydrolases superfamily protein [Source:UniProtKB/TrEMBL;Acc:O80628] |
| AT2G46400 | WRKY46 | 3.19 | 1.24 | 0.003 | 0.046 | Probable WRKY transcription factor 46 [Source:UniProtKB/Swiss-Prot;Acc:Q9SKD9] |
| AT3G04510 | LSH2 | 3.09 | 1.04 | 0.010 | 0.007 | Protein LIGHT-DEPENDENT SHORT HYPOCOTYLS 2 [Source:UniProtKB/Swiss-Prot;Acc:Q9M836] |
| AT3G55515 | DVL8 | 2.89 | 1.61 | 0.009 | 0.019 | DVL8 [Source:UniProtKB/TrEMBL;Acc:Q6IM93] |
| AT4G38000 | DOF4.7 | 2.88 | 1.17 | 0.004 | 0.007 | DOF4.7 [Source:UniProtKB/TrEMBL;Acc:A0A384KID3] |
| AT1G15002 | - | 2.25 | 2.03 | 0.021 | 0.029 | Potential natural antisense gene, locus overlaps with AT1G15000 [Source:TAIR;Acc:AT1G15002] |
| AT3G50850 | - | 1.80 | 1.69 | 0.036 | 0.026 | At3g50850 [Source:UniProtKB/TrEMBL;Acc:Q9SVL4] |
| AT5G09976 | - | 1.66 | 1.63 | 0.022 | 0.006 | FUNCTIONS IN: molecular_function unknown; INVOLVED IN: biological_process unknown; LOCATED IN: cellular_component unknown; BEST Arabidopsis thaliana protein match is: precursor of peptide 1 (TAIR:AT5G64900.1); Ha. [Source:TAIR;Acc:AT5G09976] |
| **'280-2d and 310-2d'_DEGs down-regulated** | | | | | | |
| AT2G02540 | ZHD3 | -1.94 | -1.35 | 0.004 | 0.002 | Zinc-finger homeodomain protein 3 [Source:UniProtKB/Swiss-Prot;Acc:O64722] |
