## Supplemental Table S3 for "Transcriptome and metabolome analyses revealed that narrowband 280 and 310 nm UV-B induce distinctive responses in Arabidopsis"

Enriched terms of KEGG pathway and gene ontology (GO) biological process for sample-specific DEGs.
Data analysis were performed in DAVID (http://david.abcc.ncifcrf.gov/home.jsp) and considered significant *P* -values were < 0.05.

| Term | Count | % | *P* -value |
|---|---|---|---|
| **Kegg pathway** | | | |
| **280-0d_DEGs up-regulated** | | | |
| Plant-pathogen interaction | 31 | 3.2 | 3.00E-19 |
| alpha-Linolenic acid metabolism | 5 | 0.5 | 1.20E-02 |
| Plant hormone signal transduction | 13 | 1.3 | 3.20E-02 |
| **280-0d_DEGs down-regulated** | | | |
| Plant hormone signal transduction | 32 | 2 | 9.60E-04 |
| Fatty acid elongation | 9 | 0.6 | 1.30E-03 |
| Cutin, suberine and wax biosynthesis | 7 | 0.4 | 6.90E-03 |
| Biosynthesis of amino acids | 28 | 1.8 | 7.70E-03 |
| Monobactam biosynthesis | 5 | 0.3 | 1.10E-02 |
| Glycine, serine and threonine metabolism | 11 | 0.7 | 1.80E-02 |
| Biosynthesis of secondary metabolites | 83 | 5.2 | 2.00E-02 |
| Biosynthesis of antibiotics | 40 | 2.5 | 2.80E-02 |
| RNA degradation | 14 | 0.9 | 2.90E-02 |
| **310-0d_DEGs up-regulated** | | | |
| – | | | |
| **310-0d_DEGs down-regulated** | | | |
| Ribosome biogenesis in eukaryotes | 7 | 1.8 | 3.80E-03 |
| Sulfur metabolism | 4 | 1 | 2.00E-02 |
| **280-2d_DEGs up-regulated** | | | |
| Biosynthesis of secondary metabolites | 140 | 7.1 | 3.10E-10 |
| Phenylpropanoid biosynthesis | 37 | 1.9 | 1.90E-08 |
| Metabolic pathways | 202 | 10.3 | 1.80E-06 |
| Fatty acid degradation | 14 | 0.7 | 2.00E-05 |
| alpha-Linolenic acid metabolism | 12 | 0.6 | 1.30E-04 |
| Phenylalanine, tyrosine and tryptophan biosynthesis | 15 | 0.8 | 2.20E-04 |
| Biosynthesis of antibiotics | 59 | 3 | 3.60E-04 |
| beta-Alanine metabolism | 12 | 0.6 | 3.70E-04 |
| Biosynthesis of amino acids | 38 | 1.9 | 7.40E-04 |
| Starch and sucrose metabolism | 22 | 1.1 | 1.80E-03 |
| Plant-pathogen interaction | 26 | 1.3 | 1.90E-03 |
| Phenylalanine metabolism | 11 | 0.6 | 2.30E-03 |
| Carbon metabolism | 37 | 1.9 | 2.50E-03 |
| Cyanoamino acid metabolism | 13 | 0.7 | 4.10E-03 |
| Regulation of autophagy | 8 | 0.4 | 6.20E-03 |
| Carotenoid biosynthesis | 8 | 0.4 | 9.40E-03 |
| Glutathione metabolism | 16 | 0.8 | 1.20E-02 |
| Citrate cycle (TCA cycle) | 12 | 0.6 | 1.60E-02 |
| Peroxisome | 14 | 0.7 | 3.20E-02 |
| Arginine and proline metabolism | 10 | 0.5 | 3.40E-02 |
| **280-2d_DEGs down-regulated** | | | |
| Photosynthesis | 39 | 2.1 | 2.30E-24 |
| Metabolic pathways | 171 | 9.4 | 1.60E-06 |
| Glyoxylate and dicarboxylate metabolism | 18 | 1 | 9.70E-06 |
| Carbon metabolism | 39 | 2.1 | 1.00E-05 |
| Carbon fixation in photosynthetic organisms | 16 | 0.9 | 6.50E-05 |
| Plant hormone signal transduction | 34 | 1.9 | 7.80E-04 |

| | | | |
|---|---|---|---|
| Pyruvate metabolism | 15 | 0.8 | 1.90E-03 |
| Fatty acid elongation | 9 | 0.5 | 2.00E-03 |
| Photosynthesis - antenna proteins | 7 | 0.4 | 3.30E-03 |
| Fructose and mannose metabolism | 11 | 0.6 | 1.10E-02 |
| Brassinosteroid biosynthesis | 4 | 0.2 | 1.50E-02 |
| Glycolysis / Gluconeogenesis | 15 | 0.8 | 2.60E-02 |
| Oxidative phosphorylation | 19 | 1 | 3.50E-02 |
| Phagosome | 12 | 0.7 | 3.50E-02 |
| **310-2d_DEGs up-regulated** | | | |
| – | | | |
| **310-2d_DEGs down-regulated** | | | |
| – | | | |
| **Biological process** | | | |
| **280-0d_DEGs up-regulated** | | | |
| response to chitin | 68 | 7 | 2.1E-60 |
| defense response | 90 | 9.2 | 2.8E-27 |
| plant-type hypersensitive response | 23 | 2.4 | 1.2E-14 |
| signal transduction | 54 | 5.5 | 6.0E-14 |
| protein phosphorylation | 80 | 8.2 | 7.3E-14 |
| response to wounding | 34 | 3.5 | 3.3E-13 |
| defense response to bacterium | 40 | 4.1 | 4.7E-13 |
| response to bacterium | 23 | 2.4 | 1.4E-11 |
| response to fungus | 15 | 1.5 | 2.9E-07 |
| response to water deprivation | 30 | 3.1 | 4.9E-07 |
| response to abscisic acid | 37 | 3.8 | 5.1E-07 |
| response to oxidative stress | 30 | 3.1 | 1.2E-06 |
| ethylene-activated signaling pathway | 22 | 2.3 | 2.7E-06 |
| response to cold | 29 | 3 | 5.9E-06 |
| response to salicylic acid | 19 | 1.9 | 1.6E-05 |
| defense response, incompatible interaction | 8 | 0.8 | 2.3E-05 |
| defense response to bacterium, incompatible interaction | 10 | 1 | 2.6E-05 |
| cellular response to hypoxia | 8 | 0.8 | 3.1E-05 |
| defense response to fungus | 35 | 3.6 | 1.1E-04 |
| protein autophosphorylation | 16 | 1.6 | 1.3E-04 |
| negative regulation of defense response | 7 | 0.7 | 2.3E-04 |
| vasculature development | 7 | 0.7 | 2.9E-04 |
| response to molecule of bacterial origin | 6 | 0.6 | 5.0E-04 |
| response to salt stress | 34 | 3.5 | 5.0E-04 |
| response to jasmonic acid | 16 | 1.6 | 6.4E-04 |
| abscisic acid-activated signaling pathway | 18 | 1.8 | 8.8E-04 |
| regulation of salicylic acid biosynthetic process | 4 | 0.4 | 9.1E-04 |
| cell death | 8 | 0.8 | 1.0E-03 |
| protein autoubiquitination | 5 | 0.5 | 1.3E-03 |
| regulation of jasmonic acid mediated signaling pathway | 6 | 0.6 | 1.6E-03 |
| response to osmotic stress | 13 | 1.3 | 1.8E-03 |
| cell surface receptor signaling pathway | 8 | 0.8 | 2.0E-03 |
| defense response by callose deposition in cell wall | 5 | 0.5 | 2.3E-03 |
| positive regulation of cell death | 4 | 0.4 | 2.4E-03 |
| regulation of defense response | 9 | 0.9 | 2.5E-03 |
| response to karrikin | 13 | 1.3 | 2.7E-03 |
| response to hydrogen peroxide | 8 | 0.8 | 3.4E-03 |
| response to other organism | 8 | 0.8 | 3.8E-03 |
| response to high light intensity | 8 | 0.8 | 3.8E-03 |

| | | | |
|---|---|---|---|
| cellular transition metal ion homeostasis | 9 | 0.9 | 4.0E-03 |
| response to hypoxia | 7 | 0.7 | 4.1E-03 |
| camalexin biosynthetic process | 4 | 0.4 | 4.9E-03 |
| cellular heat acclimation | 4 | 0.4 | 4.9E-03 |
| leaf senescence | 10 | 1 | 6.6E-03 |
| response to ozone | 6 | 0.6 | 6.6E-03 |
| defense response to oomycetes | 6 | 0.6 | 7.6E-03 |
| indole glucosinolate catabolic process | 3 | 0.3 | 7.7E-03 |
| cellular response to chitin | 3 | 0.3 | 7.7E-03 |
| circadian rhythm | 10 | 1 | 9.3E-03 |
| cold acclimation | 7 | 0.7 | 1.1E-02 |
| jasmonic acid biosynthetic process | 5 | 0.5 | 1.2E-02 |
| systemic acquired resistance, salicylic acid mediated signaling pathway | 4 | 0.4 | 1.3E-02 |
| cell wall macromolecule catabolic process | 5 | 0.5 | 1.4E-02 |
| cellular response to heat | 5 | 0.5 | 1.4E-02 |
| salicylic acid mediated signaling pathway | 5 | 0.5 | 1.4E-02 |
| response to heat | 13 | 1.3 | 1.5E-02 |
| systemic acquired resistance | 6 | 0.6 | 1.7E-02 |
| metal ion transport | 10 | 1 | 1.8E-02 |
| protein ubiquitination | 27 | 2.8 | 1.8E-02 |
| regulation of stomatal complex patterning | 3 | 0.3 | 1.8E-02 |
| defense response to fungus, incompatible interaction | 6 | 0.6 | 2.2E-02 |
| starch catabolic process | 4 | 0.4 | 2.3E-02 |
| response to UV-B | 7 | 0.7 | 2.4E-02 |
| response to molecule of fungal origin | 3 | 0.3 | 2.5E-02 |
| cellular response to molecule of bacterial origin | 3 | 0.3 | 2.5E-02 |
| induced systemic resistance | 4 | 0.4 | 2.7E-02 |
| regulation of salicylic acid mediated signaling pathway | 4 | 0.4 | 2.7E-02 |
| negative regulation of cell death | 3 | 0.3 | 3.3E-02 |
| regulation of salicylic acid metabolic process | 3 | 0.3 | 3.3E-02 |
| positive regulation of abscisic acid-activated signaling pathway | 5 | 0.5 | 3.5E-02 |
| calcium ion transmembrane transport | 4 | 0.4 | 3.5E-02 |
| protein dephosphorylation | 9 | 0.9 | 3.6E-02 |
| innate immune response | 7 | 0.7 | 3.6E-02 |
| transcription, DNA-templated | 85 | 8.7 | 3.6E-02 |
| response to ethylene | 10 | 1 | 3.9E-02 |
| response to stress | 7 | 0.7 | 4.1E-02 |
| response to mechanical stimulus | 3 | 0.3 | 4.1E-02 |
| **280-0d_DEGs down-regulated** | | | |
| auxin-activated signaling pathway | 38 | 2.4 | 5.9E-10 |
| response to auxin | 49 | 3.1 | 6.2E-10 |
| transcription, DNA-templated | 180 | 11.3 | 7.1E-09 |
| cuticle development | 13 | 0.8 | 6.8E-08 |
| regulation of transcription, DNA-templated | 185 | 11.6 | 2.2E-06 |
| positive regulation of auxin mediated signaling pathway | 6 | 0.4 | 1.8E-05 |
| fatty acid metabolic process | 14 | 0.9 | 6.0E-05 |
| cell cycle | 21 | 1.3 | 7.1E-05 |
| response to abscisic acid | 45 | 2.8 | 1.5E-04 |
| response to brassinosteroid | 10 | 0.6 | 1.8E-04 |
| response to light stimulus | 26 | 1.6 | 3.1E-04 |
| water transport | 6 | 0.4 | 3.3E-04 |
| embryo development ending in seed dormancy | 49 | 3.1 | 3.7E-04 |
| wax biosynthetic process | 8 | 0.5 | 3.8E-04 |

| | | | |
|---|---|---|---|
| response to gibberellin | 17 | 1.1 | 4.3E-04 |
| mucilage biosynthetic process | 6 | 0.4 | 5.3E-04 |
| unidimensional cell growth | 18 | 1.1 | 7.5E-04 |
| brassinosteroid mediated signaling pathway | 12 | 0.8 | 9.9E-04 |
| fatty acid biosynthetic process | 19 | 1.2 | 1.1E-03 |
| regulation of intracellular pH | 8 | 0.5 | 1.4E-03 |
| syncytium formation | 6 | 0.4 | 1.7E-03 |
| stomatal movement | 8 | 0.5 | 1.7E-03 |
| regulation of timing of transition from vegetative to reproductive phase | 9 | 0.6 | 2.1E-03 |
| regulation of monopolar cell growth | 4 | 0.3 | 2.2E-03 |
| auxin polar transport | 11 | 0.7 | 2.3E-03 |
| very long-chain fatty acid metabolic process | 6 | 0.4 | 2.4E-03 |
| circadian rhythm | 15 | 0.9 | 2.9E-03 |
| basipetal auxin transport | 8 | 0.5 | 3.2E-03 |
| cellular amino acid biosynthetic process | 12 | 0.8 | 3.6E-03 |
| xyloglucan metabolic process | 9 | 0.6 | 4.1E-03 |
| flower development | 24 | 1.5 | 4.2E-03 |
| purine nucleotide biosynthetic process | 4 | 0.3 | 4.3E-03 |
| response to ethylene | 17 | 1.1 | 4.7E-03 |
| regulation of organ growth | 5 | 0.3 | 5.1E-03 |
| potassium ion transmembrane transport | 8 | 0.5 | 5.4E-03 |
| plant-type primary cell wall biogenesis | 6 | 0.4 | 6.8E-03 |
| ion transmembrane transport | 7 | 0.4 | 6.9E-03 |
| cellular water homeostasis | 8 | 0.5 | 7.4E-03 |
| asymmetric cell division | 5 | 0.3 | 9.3E-03 |
| negative regulation of transcription, DNA-templated | 17 | 1.1 | 9.9E-03 |
| DNA-dependent DNA replication | 6 | 0.4 | 1.0E-02 |
| epidermal cell differentiation | 4 | 0.3 | 1.1E-02 |
| shade avoidance | 5 | 0.3 | 1.2E-02 |
| trichome differentiation | 6 | 0.4 | 1.3E-02 |
| cell wall organization | 31 | 1.9 | 1.5E-02 |
| phototropism | 5 | 0.3 | 1.9E-02 |
| RNA processing | 13 | 0.8 | 2.1E-02 |
| lateral root formation | 8 | 0.5 | 2.1E-02 |
| threonine biosynthetic process | 4 | 0.3 | 2.1E-02 |
| very long-chain fatty acid biosynthetic process | 5 | 0.3 | 2.3E-02 |
| leaf development | 15 | 0.9 | 2.5E-02 |
| chloroplast organization | 15 | 0.9 | 2.5E-02 |
| gravitropism | 7 | 0.4 | 2.6E-02 |
| lipid catabolic process | 18 | 1.1 | 2.8E-02 |
| negative regulation of photomorphogenesis | 4 | 0.3 | 2.8E-02 |
| fatty acid elongation | 4 | 0.3 | 2.8E-02 |
| lipid biosynthetic process | 4 | 0.3 | 2.8E-02 |
| shoot system development | 10 | 0.6 | 2.8E-02 |
| trichome branching | 6 | 0.4 | 2.8E-02 |
| protein ubiquitination | 40 | 2.5 | 3.1E-02 |
| auxin efflux | 5 | 0.3 | 3.3E-02 |
| cutin biosynthetic process | 5 | 0.3 | 3.3E-02 |
| transmembrane receptor protein tyrosine kinase signaling pathway | 15 | 0.9 | 3.4E-02 |
| response to salicylic acid | 17 | 1.1 | 3.5E-02 |
| pattern specification process | 4 | 0.3 | 3.5E-02 |
| protein phosphorylation | 69 | 4.3 | 3.6E-02 |
| cell adhesion | 4 | 0.3 | 4.4E-02 |

| | | | |
|---|---|---|---|
| auxin transport | 4 | 0.3 | 4.4E-02 |
| cell fate specification | 4 | 0.3 | 4.4E-02 |
| methylation | 21 | 1.3 | 4.5E-02 |
| regulation of transcription from RNA polymerase II promoter | 21 | 1.3 | 4.7E-02 |
| cotyledon development | 6 | 0.4 | 4.8E-02 |
| **310-0d_DEGs up-regulated** | | | |
| response to karrikin | 6 | 4.7 | 4.0E-04 |
| glucose import | 4 | 3.1 | 1.5E-03 |
| negative regulation of seed germination | 3 | 2.3 | 4.6E-03 |
| response to light stimulus | 5 | 3.9 | 1.3E-02 |
| hexose transmembrane transport | 3 | 2.3 | 1.7E-02 |
| flavonoid glucuronidation | 4 | 3.1 | 1.7E-02 |
| flavonoid biosynthetic process | 4 | 3.1 | 3.4E-02 |
| response to water deprivation | 5 | 3.9 | 4.7E-02 |
| regulation of transcription, DNA-templated | 17 | 13.3 | 4.9E-02 |
| **310-0d_DEGs down-regulated** | | | |
| rRNA processing | 11 | 2.9 | 1.4E-07 |
| glucosinolate biosynthetic process | 5 | 1.3 | 1.6E-03 |
| response to gibberellin | 7 | 1.8 | 2.9E-03 |
| embryo sac development | 6 | 1.6 | 3.8E-03 |
| response to salicylic acid | 8 | 2.1 | 6.8E-03 |
| protein folding | 10 | 2.6 | 2.2E-02 |
| cysteine biosynthetic process | 3 | 0.8 | 2.4E-02 |
| response to wounding | 8 | 2.1 | 2.4E-02 |
| regulation of transcription from RNA polymerase II promoter | 8 | 2.1 | 3.3E-02 |
| regulation of transcription, DNA-templated | 41 | 10.7 | 3.8E-02 |
| glucosinolate biosynthetic process from homomethionine | 2 | 0.5 | 4.2E-02 |
| **280-2d_DEGs up-regulated** | | | |
| defense response to bacterium | 71 | 3.6 | 1.2E-19 |
| response to bacterium | 37 | 1.9 | 1.6E-15 |
| defense response | 108 | 5.5 | 7.0E-14 |
| response to salicylic acid | 43 | 2.2 | 2.7E-13 |
| response to wounding | 48 | 2.4 | 2.5E-12 |
| oxidation-reduction process | 168 | 8.5 | 8.6E-11 |
| plant-type hypersensitive response | 25 | 1.3 | 6.2E-10 |
| systemic acquired resistance | 18 | 0.9 | 2.7E-09 |
| response to salt stress | 76 | 3.9 | 3.9E-09 |
| leaf senescence | 25 | 1.3 | 7.5E-08 |
| response to oxidative stress | 50 | 2.5 | 1.6E-07 |
| response to jasmonic acid | 33 | 1.7 | 2.6E-07 |
| response to fungus | 21 | 1.1 | 3.0E-07 |
| response to other organism | 18 | 0.9 | 4.3E-07 |
| amino sugar metabolic process | 8 | 0.4 | 5.8E-05 |
| defense response to bacterium, incompatible interaction | 13 | 0.7 | 8.5E-05 |
| response to abscisic acid | 53 | 2.7 | 9.1E-05 |
| response to karrikin | 24 | 1.2 | 1.1E-04 |
| tricarboxylic acid cycle | 14 | 0.7 | 1.4E-04 |
| autophagy | 13 | 0.7 | 1.7E-04 |
| response to water deprivation | 40 | 2 | 2.0E-04 |
| response to molecule of bacterial origin | 8 | 0.4 | 3.5E-04 |
| cellular response to nitrogen starvation | 9 | 0.5 | 4.0E-04 |
| signal transduction | 56 | 2.8 | 4.2E-04 |
| response to chitin | 23 | 1.2 | 5.0E-04 |

| | | | |
|---|---|---|---|
| response to ozone | 10 | 0.5 | 6.5E-04 |
| regulation of systemic acquired resistance | 6 | 0.3 | 1.3E-03 |
| nitrate assimilation | 11 | 0.6 | 1.6E-03 |
| defense response to fungus, incompatible interaction | 11 | 0.6 | 1.6E-03 |
| oligopeptide transport | 14 | 0.7 | 1.6E-03 |
| response to cadmium ion | 43 | 2.2 | 1.8E-03 |
| lignin biosynthetic process | 14 | 0.7 | 1.8E-03 |
| amino acid homeostasis | 5 | 0.3 | 1.9E-03 |
| response to toxic substance | 13 | 0.7 | 2.1E-03 |
| defense response, incompatible interaction | 8 | 0.4 | 2.2E-03 |
| protein phosphorylation | 91 | 4.6 | 2.4E-03 |
| systemic acquired resistance, salicylic acid mediated signaling pathway | 6 | 0.3 | 3.0E-03 |
| pathogen-associated molecular pattern dependent induction by symbiont of host i | 5 | 0.3 | 3.2E-03 |
| secondary metabolic process | 8 | 0.4 | 3.5E-03 |
| defense response to fungus | 53 | 2.7 | 4.0E-03 |
| response to hypoxia | 10 | 0.5 | 4.0E-03 |
| phenylpropanoid biosynthetic process | 9 | 0.5 | 4.3E-03 |
| response to herbivore | 5 | 0.3 | 5.0E-03 |
| protein transport | 40 | 2 | 7.6E-03 |
| regulation of salicylic acid biosynthetic process | 4 | 0.2 | 7.6E-03 |
| amino acid import | 4 | 0.2 | 7.6E-03 |
| regulation of defense response | 12 | 0.6 | 1.0E-02 |
| cell wall macromolecule catabolic process | 7 | 0.4 | 1.2E-02 |
| salicylic acid mediated signaling pathway | 7 | 0.4 | 1.2E-02 |
| response to molecule of fungal origin | 4 | 0.2 | 1.3E-02 |
| defense response to oomycetes | 8 | 0.4 | 1.3E-02 |
| defense response to other organism | 9 | 0.5 | 1.3E-02 |
| protein autophosphorylation | 19 | 1 | 1.5E-02 |
| nitrate transport | 6 | 0.3 | 1.6E-02 |
| negative regulation of response to water deprivation | 3 | 0.2 | 1.7E-02 |
| drug transmembrane transport | 12 | 0.6 | 1.7E-02 |
| cell death | 9 | 0.5 | 1.8E-02 |
| mitophagy | 5 | 0.3 | 1.9E-02 |
| glutathione metabolic process | 11 | 0.6 | 1.9E-02 |
| negative regulation of programmed cell death | 5 | 0.3 | 2.4E-02 |
| abscisic acid biosynthetic process | 5 | 0.3 | 2.4E-02 |
| response to nematode | 12 | 0.6 | 2.5E-02 |
| proteolysis involved in cellular protein catabolic process | 15 | 0.8 | 2.6E-02 |
| flavonol biosynthetic process | 4 | 0.2 | 2.7E-02 |
| metabolic process | 34 | 1.7 | 2.7E-02 |
| recognition of pollen | 8 | 0.4 | 2.7E-02 |
| reactive oxygen species metabolic process | 5 | 0.3 | 3.0E-02 |
| phosphatidylinositol phosphorylation | 5 | 0.3 | 3.0E-02 |
| innate immune response | 11 | 0.6 | 3.1E-02 |
| amino acid export | 3 | 0.2 | 3.2E-02 |
| polyamine catabolic process | 3 | 0.2 | 3.2E-02 |
| salicylic acid catabolic process | 3 | 0.2 | 3.2E-02 |
| oxylipin biosynthetic process | 6 | 0.3 | 3.3E-02 |
| regulation of jasmonic acid mediated signaling pathway | 6 | 0.3 | 3.3E-02 |
| fatty acid beta-oxidation | 8 | 0.4 | 3.5E-02 |
| protein tetramerization | 4 | 0.2 | 3.6E-02 |
| phenylpropanoid metabolic process | 5 | 0.3 | 3.7E-02 |
| aromatic amino acid family biosynthetic process | 5 | 0.3 | 3.7E-02 |

| | | | |
|---|---|---|---|
| response to starvation | 5 | 0.3 | 3.7E-02 |
| cellular amino acid biosynthetic process | 11 | 0.6 | 3.8E-02 |
| jasmonic acid biosynthetic process | 6 | 0.3 | 3.9E-02 |
| chitin catabolic process | 6 | 0.3 | 3.9E-02 |
| negative regulation of defense response | 6 | 0.3 | 3.9E-02 |
| hydrogen peroxide catabolic process | 13 | 0.7 | 4.0E-02 |
| positive regulation of abscisic acid-activated signaling pathway | 7 | 0.4 | 4.3E-02 |
| induced systemic resistance | 5 | 0.3 | 4.5E-02 |
| floral organ abscission | 6 | 0.3 | 4.5E-02 |
| response to insect | 6 | 0.3 | 4.5E-02 |
| multicellular organism development | 40 | 2 | 4.6E-02 |
| negative regulation of leaf senescence | 4 | 0.2 | 4.7E-02 |
| galactolipid biosynthetic process | 4 | 0.2 | 4.7E-02 |
| flavonoid glucuronidation | 15 | 0.8 | 4.8E-02 |
| **280-2d_DEGs down-regulated** | | | |
| photosynthesis | 37 | 2 | 2.8E-12 |
| response to light stimulus | 40 | 2.2 | 6.3E-10 |
| circadian rhythm | 27 | 1.5 | 1.6E-09 |
| auxin-activated signaling pathway | 39 | 2.1 | 3.1E-09 |
| response to far red light | 18 | 1 | 1.5E-08 |
| protein-chromophore linkage | 16 | 0.9 | 2.0E-07 |
| photosynthesis, light reaction | 11 | 0.6 | 2.6E-07 |
| response to blue light | 17 | 0.9 | 2.7E-07 |
| transmembrane receptor protein tyrosine kinase signaling pathway | 27 | 1.5 | 9.2E-07 |
| brassinosteroid mediated signaling pathway | 17 | 0.9 | 1.8E-06 |
| reductive pentose-phosphate cycle | 10 | 0.5 | 3.2E-06 |
| response to cytokinin | 32 | 1.8 | 3.7E-06 |
| response to auxin | 43 | 2.4 | 5.3E-06 |
| response to brassinosteroid | 12 | 0.7 | 1.2E-05 |
| photosystem II assembly | 9 | 0.5 | 1.3E-05 |
| photosynthetic electron transport chain | 8 | 0.4 | 5.2E-05 |
| photosynthetic electron transport in photosystem I | 8 | 0.4 | 5.2E-05 |
| positive gravitropism | 11 | 0.6 | 5.6E-05 |
| cuticle development | 10 | 0.5 | 1.1E-04 |
| unidimensional cell growth | 21 | 1.2 | 1.1E-04 |
| fatty acid biosynthetic process | 22 | 1.2 | 2.1E-04 |
| cotyledon development | 10 | 0.5 | 2.5E-04 |
| response to cold | 39 | 2.1 | 2.5E-04 |
| microtubule-based process | 9 | 0.5 | 2.8E-04 |
| water transport | 6 | 0.3 | 5.3E-04 |
| phototropism | 7 | 0.4 | 7.3E-04 |
| plant-type secondary cell wall biogenesis | 11 | 0.6 | 7.4E-04 |
| cellular water homeostasis | 10 | 0.5 | 8.0E-04 |
| cell wall organization | 38 | 2.1 | 1.2E-03 |
| carbon fixation | 5 | 0.3 | 1.3E-03 |
| response to karrikin | 20 | 1.1 | 1.4E-03 |
| response to red light | 12 | 0.7 | 2.0E-03 |
| nonphotochemical quenching | 5 | 0.3 | 2.2E-03 |
| ion transmembrane transport | 8 | 0.4 | 2.6E-03 |
| cell redox homeostasis | 22 | 1.2 | 2.6E-03 |
| fructose metabolic process | 4 | 0.2 | 3.1E-03 |
| photosynthesis, light harvesting in photosystem I | 7 | 0.4 | 3.2E-03 |
| gravitropism | 9 | 0.5 | 3.5E-03 |
| response to gibberellin | 16 | 0.9 | 3.5E-03 |

| | | | |
|---|---|---|---|
| response to abscisic acid | 43 | 2.4 | 4.1E-03 |
| wax biosynthetic process | 7 | 0.4 | 4.1E-03 |
| regulation of growth | 16 | 0.9 | 4.3E-03 |
| gibberellic acid mediated signaling pathway | 13 | 0.7 | 4.4E-03 |
| hyperosmotic salinity response | 11 | 0.6 | 5.7E-03 |
| chloroplast relocation | 4 | 0.2 | 5.8E-03 |
| regulation of transcription, DNA-templated | 177 | 9.7 | 7.1E-03 |
| response to low light intensity stimulus | 4 | 0.2 | 9.6E-03 |
| photosynthesis, light harvesting | 5 | 0.3 | 1.0E-02 |
| transport | 37 | 2 | 1.2E-02 |
| response to ethylene | 17 | 0.9 | 1.3E-02 |
| cellular response to light stimulus | 5 | 0.3 | 1.3E-02 |
| transcription, DNA-templated | 157 | 8.6 | 1.4E-02 |
| epidermal cell differentiation | 4 | 0.2 | 1.5E-02 |
| auxin polar transport | 10 | 0.5 | 1.5E-02 |
| cell differentiation | 32 | 1.8 | 1.6E-02 |
| ATP synthesis coupled proton transport | 8 | 0.4 | 1.7E-02 |
| shade avoidance | 5 | 0.3 | 1.7E-02 |
| regulation of seed dormancy process | 4 | 0.2 | 2.1E-02 |
| photosynthetic electron transport in photosystem II | 4 | 0.2 | 2.1E-02 |
| photorespiration | 9 | 0.5 | 2.2E-02 |
| very long-chain fatty acid metabolic process | 5 | 0.3 | 2.2E-02 |
| fructose 6-phosphate metabolic process | 5 | 0.3 | 2.2E-02 |
| plant-type cell wall organization | 13 | 0.7 | 2.4E-02 |
| response to jasmonic acid | 19 | 1 | 2.4E-02 |
| glycolytic process | 11 | 0.6 | 2.5E-02 |
| fructose 1,6-bisphosphate metabolic process | 3 | 0.2 | 2.7E-02 |
| chloroplast organization | 16 | 0.9 | 2.8E-02 |
| brassinosteroid biosynthetic process | 7 | 0.4 | 2.9E-02 |
| lipid transport | 18 | 1 | 3.3E-02 |
| regulation of root meristem growth | 6 | 0.3 | 4.2E-02 |
| glycine catabolic process | 3 | 0.2 | 4.2E-02 |
| regulation of auxin biosynthetic process | 3 | 0.2 | 4.2E-02 |
| programmed cell death involved in cell development | 3 | 0.2 | 4.2E-02 |
| chloroplast ribulose bisphosphate carboxylase complex biogenesis | 3 | 0.2 | 4.2E-02 |
| regulation of starch metabolic process | 3 | 0.2 | 4.2E-02 |
| oxidation-reduction process | 110 | 6 | 4.3E-02 |
| cellulose biosynthetic process | 8 | 0.4 | 4.3E-02 |
| plant-type cell wall modification | 4 | 0.2 | 4.6E-02 |
| cutin biosynthetic process | 5 | 0.3 | 4.7E-02 |
| **310-2d_DEGs up-regulated** | | | |
| – | | | |
| **310-2d_DEGs down-regulated** | | | |
| – | | | |
