## Supplemental Table S4 for "Transcriptome and metabolome analyses revealed that narrowband 280 and 310 nm UV-B induce distinctive responses in Arabidopsis"

List of metabolites whose amounts were significantly changed by UV-LED irradiation.
Metabolites were annotated by GC-MS analysis. Value is mean of tripricated measurements and significant difference ($P < 0.05$) was evaluated between peark areas of each metabolites in C-2d and 280/310-2d.

| | | Peak Area | | | *P* -value | |
|---|---|---|---|---|---|---|
| ID | Annotation | C-2d | 280-2d | 310-2d | 280-2d | 310-2d |
| **Hydrophilic metabolites** | | | | | | |
| 3 | Pyruvate | 6907 | 11573 | 5517 | 0.034 | 0.278 |
| 20 | 3-Hydroxypropanoate | 16582 | 44668 | 16596 | 0.021 | 0.997 |
| 44 | Urea | 272447 | 899258 | 183748 | 0.045 | 0.183 |
| 56 | Ethanolamine | 2241503 | 3216703 | 2175413 | 0.024 | 0.790 |
| 71 | Maleic acid | 125321 | 278151 | 206837 | 0.011 | 0.068 |
| 73 | Succinate | 39319 | 213586 | 41952 | 0.006 | 0.886 |
| 78 | Fumarate | 1298448 | 3888598 | 1302608 | 0.003 | 0.990 |
| 115 | Malate | 298492 | 2846827 | 313624 | 0.000 | 0.911 |
| 126 | Aspartate | 83191 | 121301 | 84673 | 0.037 | 0.897 |
| 134 | γ-Aminobutyric acid | 219306 | 460207 | 185952 | 0.025 | 0.676 |
| 143 | Cysteine | 99651 | 67243 | 83672 | 0.042 | 0.285 |
| 150 | 2-Oxoglutarate | 15368 | 95187 | 12702 | 0.000 | 0.497 |
| 151 | 2-Hydroxyglutarate | 9987 | 83453 | 9107 | 0.001 | 0.808 |
| 227 | Aconitate | 15483 | 45355 | 9873 | 0.033 | 0.301 |
| 236 | Glycerol 3-phosphate | 31489 | 20891 | 22700 | 0.031 | 0.159 |
| 246 | Shikimate | 39257 | 105195 | 45372 | 0.004 | 0.592 |
| 256 | Citrate | 373346 | 985698 | 347596 | 0.006 | 0.800 |
| 283 | Adenine | 15163 | 35909 | 15842 | 0.000 | 0.631 |
| 313 | Lysine | 554851 | 227402 | 534773 | 0.001 | 0.680 |
| 314 | Histidine | 212491 | 377490 | 158834 | 0.014 | 0.315 |
| 350 | Hexadecanoic acid | 14515 | 25594 | 14089 | 0.016 | 0.898 |
| **Fatty acid** | | | | | | |
| 3 | Octanoic acid (caprylate;8:0) | -957 | 3818 | 4152 | 0.037 | 0.030 |
| 4 | Decanoic acid (caprate;10:0) | 401 | 3090 | 848 | 0.006 | 0.018 |
| 6 | Dodecanoic acid (laurate;12:0) | 485 | 3039 | 659 | 0.008 | 0.229 |
| 9 | Tetradecanoic acid (myristate;14:0) | 7393 | 12658 | 8555 | 0.002 | 0.355 |
| 11 | Pentadecanoate;15:0 | 1711 | 3307 | 2063 | 0.002 | 0.379 |
| 21 | Octadecenoic acid (oleate;(*Z*)18:1n-9) | 4096 | 6001 | 4028 | 0.018 | 0.852 |
| 40 | Docosanoic acid (behenate;22:0) | 12538 | 16826 | 13259 | 0.023 | 0.673 |
