## Supplemental Table S5 for "Transcriptome and metabolome analyses revealed that narrowband 280 and 310 nm UV-B induce distinctive responses in Arabidopsis"

DEGs specifically up-regulated in 310-0d

| ID | Gene name | $Log_2$ FC | | *P* -value | | Description |
|---|---|---|---|---|---|---|
| | | 280-0d | 310-0d | 280-0d | 310-0d | |
| AT3G24750 | - | 5.38 | 10.76 | 0.185 | 0.014 | unknown protein; Ha. [Source:TAIR;Acc:AT3G24750] |
| AT1G22810 | ERF019 | 9.69 | 10.08 | 0.003 | 0.003 | Ethylene-responsive transcription factor ERF019 [Source:UniProtKB/Swiss-Prot;Acc:O80542] |
| AT1G65330 | PHE1 | 5.42 | 8.89 | 0.196 | 0.038 | MADS-box transcription factor PHERES 1 [Source:UniProtKB/Swiss-Prot;Acc:O80805] |
| AT2G29090 | CYP707A2 | 1.52 | 3.43 | 0.084 | 0.008 | Abscisic acid 8'-hydroxylase 2 [Source:UniProtKB/Swiss-Prot;Acc:O81077] |
| AT2G40130 | SMXL8 | 1.11 | 3.19 | 0.089 | 0.007 | Protein SMAX1-LIKE 8 [Source:UniProtKB/Swiss-Prot;Acc:F4IGZ2] |
| AT1G06980 | - | 1.68 | 3.05 | 0.100 | 0.034 | 6,7-dimethyl-8-ribityllumazine synthase [Source:UniProtKB/TrEMBL;Acc:Q9LMJ2] |
| AT3G30460 | - | 0.94 | 2.96 | 0.174 | 0.032 | RING/U-box superfamily protein [Source:UniProtKB/TrEMBL;Acc:A0A1I9LPW5] |
| AT5G48050 | - | 2.66 | 2.68 | 0.020 | 0.044 | Copia-like polyprotein/retrotransposon [Source:UniProtKB/TrEMBL;Acc:Q9FI33] |
| AT1G02340 | HFR1 | 0.40 | 2.67 | 0.341 | 0.007 | Transcription factor HFR1 [Source:UniProtKB/Swiss-Prot;Acc:Q9FE22] |
| AT4G26150 | GATA22 | 0.39 | 2.64 | 0.198 | 0.021 | Putative GATA transcription factor 22 [Source:UniProtKB/Swiss-Prot;Acc:Q9SZI6] |
| AT4G36450 | MPK14 | 1.67 | 2.38 | 0.192 | 0.003 | Mitogen-activated protein kinase 14 [Source:UniProtKB/Swiss-Prot;Acc:O23236] |
| AT3G29370 | - | 0.95 | 2.35 | 0.224 | 0.022 | Uncharacterized protein At3g29370 [Source:UniProtKB/TrEMBL;Acc:Q8LD48] |
| AT2G28630 | KCS12 | 0.58 | 2.12 | 0.099 | 0.010 | 3-ketoacyl-CoA synthase 12 [Source:UniProtKB/Swiss-Prot;Acc:Q9SIB2] |
| AT1G11260 | STP1 | 1.23 | 2.11 | 0.083 | 0.028 | STP1 [Source:UniProtKB/TrEMBL;Acc:A0A178WJ63] |
| AT5G66400 | RAB18 | 2.22 | 2.08 | 0.127 | 0.040 | RAB18 [Source:UniProtKB/TrEMBL;Acc:A0A178UN24] |
| AT5G67480 | BT4 | 0.77 | 2.01 | 0.029 | 0.038 | BTB and TAZ domain protein 4 [Source:TAIR;Acc:AT5G67480] |
| AT3G18550 | BRC1 | -0.32 | 1.99 | 0.633 | 0.013 | TCP family transcription factor [Source:UniProtKB/TrEMBL;Acc:A0A1I9LSP1] |
| AT4G15500 | UGT84A4 | -0.71 | 1.95 | 0.243 | 0.042 | UDP-glycosyltransferase 84A4 [Source:UniProtKB/Swiss-Prot;Acc:O23402] |
| AT1G75960 | AAE8 | 0.17 | 1.90 | 0.704 | 0.014 | Probable acyl-activating enzyme 8 [Source:UniProtKB/Swiss-Prot;Acc:Q9LQS1] |
| AT1G08570 | ACHT4 | 0.87 | 1.89 | 0.005 | 0.006 | Thioredoxin-like 1-1, chloroplastic [Source:UniProtKB/Swiss-Prot;Acc:O64654] |
| AT3G14460 | - | -1.73 | 1.85 | 0.594 | 0.016 | Putative disease resistance protein At3g14460 [Source:UniProtKB/Swiss-Prot;Acc:Q9LRR5] |
| AT4G00970 | CRK41 | 1.00 | 1.85 | 0.006 | 0.019 | Cysteine-rich receptor-like protein kinase 41 [Source:UniProtKB/Swiss-Prot;Acc:O23081] |
| AT3G57520 | RFS2 | 0.16 | 1.81 | 0.609 | 0.002 | Probable galactinol--sucrose galactosyltransferase 2 [Source:UniProtKB/Swiss-Prot;Acc:Q94A08] |
| AT3G09162 | - | 1.19 | 1.80 | 0.063 | 0.038 | At3g09162 [Source:UniProtKB/TrEMBL;Acc:Q8GYI0] |
| AT1G19370 | - | 1.00 | 1.78 | 0.076 | 0.038 | F18O14.9 [Source:UniProtKB/TrEMBL;Acc:Q9LN61] |
| AT1G52770 | - | -1.44 | 1.78 | 0.342 | 0.036 | At1g52770 [Source:UniProtKB/TrEMBL;Acc:Q9C941] |
| AT2G39400 | - | 0.82 | 1.75 | 0.001 | 0.024 | Alpha/beta-Hydrolases superfamily protein [Source:UniProtKB/TrEMBL;Acc:O80627] |
| AT3G46370 | - | 1.58 | 1.71 | 0.055 | 0.047 | Leucine-rich repeat protein kinase family protein [Source:UniProtKB/TrEMBL;Acc:Q9SNA0] |
| AT5G03700 | - | 0.88 | 1.68 | 0.097 | 0.001 | PAN domain-containing protein At5g03700 [Source:UniProtKB/Swiss-Prot;Acc:Q9LZR8] |

| | | | | | | |
|---|---|---|---|---|---|---|
| AT1G52245 | - | -0.26 | 1.68 | 0.807 | 0.038 | Dynein light chain [Source:UniProtKB/TrEMBL;Acc:Q9M810] |
| AT2G02410 | - | 0.52 | 1.68 | 0.246 | 0.019 | unknown protein; CONTAINS InterPro DOMAIN/s: Protein of unknown function DUF901 (InterPro:IPR010298); Ha. [Source:TAIR;Acc:AT2G02410] |
| AT3G56260 | - | -0.60 | 1.67 | 0.346 | 0.046 | At3g56260 [Source:UniProtKB/TrEMBL;Acc:Q6NLT4] |
| AT5G02950 | - | 1.12 | 1.67 | 0.053 | 0.022 | Tudor/PWWP/MBT superfamily protein [Source:UniProtKB/TrEMBL;Acc:Q9LYZ0] |
| AT1G78820 | - | 0.89 | 1.65 | 0.033 | 0.049 | EP1-like glycoprotein 1 [Source:UniProtKB/Swiss-Prot;Acc:Q9ZVA1] |
| AT1G75190 | - | 0.80 | 1.62 | 0.069 | 0.013 | AT1G75190 protein [Source:UniProtKB/TrEMBL;Acc:Q9FRL0] |
| AT2G28720 | - | 0.69 | 1.61 | 0.009 | 0.018 | Histone H2B.3 [Source:UniProtKB/Swiss-Prot;Acc:Q9SI96] |
| AT1G07250 | UGT71C4 | 0.67 | 1.60 | 0.028 | 0.009 | Flavonol 3-O-glucosyltransferase UGT71C4 [Source:UniProtKB/Swiss-Prot;Acc:Q9LML6] |
| AT1G05020 | AP180 | 1.12 | 1.57 | 0.076 | 0.032 | Clathrin coat assembly protein AP180 [Source:UniProtKB/Swiss-Prot;Acc:Q9ZVN6] |
| AT5G20250 | DIN10 | 0.37 | 1.57 | 0.469 | 0.041 | Raffinose synthase family protein [Source:UniProtKB/TrEMBL;Acc:F4K470] |
| AT3G03440 | - | 1.06 | 1.55 | 0.050 | 0.001 | ARM repeat superfamily protein [Source:UniProtKB/TrEMBL;Acc:F4J139] |
| AT4G03400 | DFL2 | 0.47 | 1.54 | 0.187 | 0.011 | Indole-3-acetic acid-amido synthetase GH3.10 [Source:UniProtKB/Swiss-Prot;Acc:Q9ZNS2] |
| AT2G25200 | - | 0.68 | 1.53 | 0.130 | 0.016 | At2g25200 [Source:UniProtKB/TrEMBL;Acc:Q9SIS2] |
| AT1G77210 | STP14 | 0.68 | 1.52 | 0.198 | 0.039 | Sugar transport protein 14 [Source:UniProtKB/Swiss-Prot;Acc:Q8GW61] |
| AT1G75800 | - | 0.51 | 1.52 | 0.054 | 0.016 | At1g75800/T4O12_2 [Source:UniProtKB/TrEMBL;Acc:Q9LQT4] |
| AT1G18400 | BEE1 | -0.68 | 1.51 | 0.255 | 0.010 | Transcription factor BEE 1 [Source:UniProtKB/Swiss-Prot;Acc:Q8GZ13] |
| AT3G59510 | - | 0.70 | 1.49 | 0.114 | 0.021 | Leucine-rich repeat (LRR) family protein [Source:UniProtKB/TrEMBL;Acc:Q9M1B6] |
| AT1G07420 | SMO2-2 | 0.78 | 1.48 | 0.053 | 0.048 | Methylsterol monooxygenase 2-2 [Source:UniProtKB/Swiss-Prot;Acc:Q8VWZ8] |
| AT3G52070 | - | 0.81 | 1.47 | 0.044 | 0.009 | At3g52070 [Source:UniProtKB/TrEMBL;Acc:Q9SUZ7] |
| AT1G13740 | AFP2 | 0.65 | 1.45 | 0.103 | 0.003 | AFP2 [Source:UniProtKB/TrEMBL;Acc:A0A178WBB1] |
| AT5G40780 | LHT1 | 1.15 | 1.44 | 0.052 | 0.049 | Lysine histidine transporter 1 [Source:UniProtKB/Swiss-Prot;Acc:Q9FKS8] |
| AT1G79700 | - | 0.47 | 1.44 | 0.225 | 0.012 | Integrase-type DNA-binding superfamily protein [Source:UniProtKB/TrEMBL;Acc:A8MQS2] |
| AT4G21970 | - | 1.38 | 1.43 | 0.089 | 0.032 | Protein of unknown function, DUF584 [Source:TAIR;Acc:AT4G21970] |
| AT1G54740 | - | 0.11 | 1.43 | 0.710 | 0.008 | Protein of unknown function (DUF3049) [Source:TAIR;Acc:AT1G54740] |
| AT4G27410 | RD26 | 0.97 | 1.42 | 0.017 | 0.010 | NAC (No Apical Meristem) domain transcriptional regulator superfamily protein [Source:UniProtKB/TrEMBL;Acc:F4JIU9] |
| AT1G70300 | POT6 | 0.92 | 1.41 | 0.223 | 0.012 | Potassium transporter 6 [Source:UniProtKB/Swiss-Prot;Acc:Q8W4I4] |
| AT2G27385 | - | 0.98 | 1.40 | 0.030 | 0.017 | Pollen Ole e 1 allergen and extensin family protein [Source:UniProtKB/TrEMBL;Acc:F4IFT4] |
| AT1G52830 | IAA6 | 0.41 | 1.40 | 0.282 | 0.005 | SHY1 [Source:UniProtKB/TrEMBL;Acc:A0A384LEJ2] |
| AT4G36790 | - | 0.92 | 1.40 | 0.019 | 0.027 | Major facilitator superfamily protein [Source:UniProtKB/TrEMBL;Acc:O23203] |
| AT1G76410 | ATL8 | -0.07 | 1.39 | 0.770 | 0.017 | ATL8 [Source:UniProtKB/TrEMBL;Acc:A0A178WF21] |
| AT1G23010 | LPR1 | 0.64 | 1.39 | 0.084 | 0.008 | Multicopper oxidase LPR1 [Source:UniProtKB/Swiss-Prot;Acc:F4I4K5] |
| AT5G43860 | CLH2 | 1.10 | 1.39 | 0.099 | 0.026 | Chlorophyllase-2, chloroplastic [Source:UniProtKB/Swiss-Prot;Acc:Q9M7I7] |
| AT1G23390 | - | 0.38 | 1.37 | 0.352 | 0.037 | F-box/Kelch repeat-containing F-box family protein [Source:UniProtKB/TrEMBL;Acc:C4PVQ8] |
| AT2G37025 | TRFL8 | -0.94 | 1.37 | 0.092 | 0.005 | TRF-like 8 [Source:UniProtKB/TrEMBL;Acc:F4IPY7] |

| | | | | | | |
|---|---|---|---|---|---|---|
| AT3G45300 | IVD | 0.38 | 1.36 | 0.250 | 0.015 | Isovaleryl-CoA dehydrogenase, mitochondrial [Source:UniProtKB/Swiss-Prot;Acc:Q9SWG0] |
| AT5G56870 | BGAL4 | 0.15 | 1.36 | 0.684 | 0.008 | Beta-galactosidase 4 [Source:UniProtKB/Swiss-Prot;Acc:Q9SCV8] |
| AT4G36040 | ATJ11 | 0.81 | 1.35 | 0.015 | 0.015 | At4g36040 [Source:UniProtKB/TrEMBL;Acc:Q1H544] |
| AT4G21480 | STP12 | 0.51 | 1.34 | 0.385 | 0.038 | STP12 [Source:UniProtKB/TrEMBL;Acc:A0A178V2V7] |
| AT1G15100 | RHA2A | 0.49 | 1.33 | 0.108 | 0.016 | E3 ubiquitin-protein ligase RHA2A [Source:UniProtKB/Swiss-Prot;Acc:Q9ZT50] |
| AT5G24490 | - | 0.75 | 1.31 | 0.002 | 0.042 | 30S ribosomal protein [Source:UniProtKB/TrEMBL;Acc:Q94K97] |
| AT1G78590 | NADK3 | 0.67 | 1.30 | 0.048 | 0.007 | NAD(H) kinase 3 [Source:TAIR;Acc:AT1G78590] |
| AT5G67390 | - | 0.73 | 1.30 | 0.105 | 0.004 | At5g67390 [Source:UniProtKB/TrEMBL;Acc:Q9FN13] |
| AT1G32690 | - | 0.49 | 1.30 | 0.189 | 0.014 | At1g32690 [Source:UniProtKB/TrEMBL;Acc:Q6NNH5] |
| AT1G28230 | PUP1 | 0.93 | 1.30 | 0.025 | 0.035 | Purine permease 1 [Source:UniProtKB/Swiss-Prot;Acc:Q9FZ96] |
| AT1G13360 | - | 0.21 | 1.29 | 0.361 | 0.007 | T6J4.11 protein [Source:UniProtKB/TrEMBL;Acc:Q9FX61] |
| AT5G57180 | CIA2 | -0.77 | 1.28 | 0.041 | 0.035 | CIA2 [Source:UniProtKB/TrEMBL;Acc:A0A178UDY1] |
| AT5G50950 | FUM2 | 0.84 | 1.28 | 0.007 | 0.002 | AT5G50950 protein [Source:UniProtKB/TrEMBL;Acc:B9DFR5] |
| AT4G03420 | - | 0.73 | 1.28 | 0.023 | 0.041 | AT4g03420/F9H3_4 [Source:UniProtKB/TrEMBL;Acc:Q8S9K9] |
| AT1G78970 | LUP1 | -0.21 | 1.27 | 0.326 | 0.004 | Lupeol synthase 1 [Source:UniProtKB/Swiss-Prot;Acc:Q9C5M3] |
| AT1G12120 | - | 0.87 | 1.26 | 0.011 | 0.001 | T28K15.14 protein [Source:UniProtKB/TrEMBL;Acc:Q9FWW2] |
| AT2G45350 | CRR4 | 0.28 | 1.24 | 0.550 | 0.049 | Pentatricopeptide repeat-containing protein At2g45350, chloroplastic [Source:UniProtKB/Swiss-Prot;Acc:O22137] |
| AT5G44260 | TZF5 | -0.03 | 1.23 | 0.902 | 0.012 | Zinc finger CCCH domain-containing protein 61 [Source:UniProtKB/Swiss-Prot;Acc:Q9FKW2] |
| AT1G30360 | ERD4 | -0.30 | 1.21 | 0.067 | 0.001 | Hyperosmolality-gated Ca2+ permeable channel 3.1 [Source:UniProtKB/TrEMBL;Acc:A0A097NUQ9] |
| AT1G53163 | - | 0.75 | 1.21 | 0.318 | 0.041 | F12M16.3 [Source:UniProtKB/TrEMBL;Acc:Q9MAI8] |
| AT3G52240 | - | 0.67 | 1.20 | 0.035 | 0.010 | Transcriptional regulator ATRX [Source:UniProtKB/TrEMBL;Acc:Q8GYC9] |
| AT2G42620 | MAX2 | -0.17 | 1.19 | 0.355 | 0.020 | F-box protein MAX2 [Source:UniProtKB/Swiss-Prot;Acc:Q9SIM9] |
| AT3G44798 | - | 0.88 | 1.18 | 0.070 | 0.032 | other RNA [Source:TAIR;Acc:AT3G44798] |
| AT1G15175 | - | 0.62 | 1.17 | 0.067 | 0.012 | other RNA [Source:TAIR;Acc:AT1G15175] |
| AT3G18320 | - | 0.06 | 1.16 | 0.830 | 0.024 | F-box and associated interaction domains-containing protein [Source:TAIR;Acc:AT3G18320] |
| AT3G21760 | UGT71B2 | 0.35 | 1.16 | 0.010 | 0.017 | Glycosyltransferase (Fragment) [Source:UniProtKB/TrEMBL;Acc:W8PVD4] |
| AT5G50290 | - | 0.96 | 1.16 | 0.023 | 0.009 | Wall-associated receptor kinase galacturonan-binding protein [Source:UniProtKB/TrEMBL;Acc:Q5XV04] |
| AT5G01520 | AIRP2 | 0.34 | 1.15 | 0.253 | 0.000 | AtAIRP2 [Source:UniProtKB/TrEMBL;Acc:A0A178U9T4] |
| AT4G36530 | - | 0.74 | 1.15 | 0.029 | 0.024 | Alpha/beta-Hydrolases superfamily protein [Source:UniProtKB/TrEMBL;Acc:O23227] |
| AT3G57680 | CTPA3 | 1.11 | 1.15 | 0.292 | 0.011 | Carboxyl-terminal-processing peptidase 3, chloroplastic [Source:UniProtKB/Swiss-Prot;Acc:F4J3G5] |
| AT1G80310 | MOT2 | 0.99 | 1.14 | 0.005 | 0.014 | MOT2 [Source:UniProtKB/TrEMBL;Acc:A0A178WGR3] |
| AT1G53450 | - | 0.81 | 1.14 | 0.066 | 0.037 | unknown protein; BEST Arabidopsis thaliana protein match is: unknown protein (TAIR:AT3G14830.2); Ha. [Source:TAIR;Acc:AT1G53450] |
| AT1G51805 | - | 0.85 | 1.13 | 0.024 | 0.010 | Leucine-rich repeat protein kinase family protein [Source:UniProtKB/TrEMBL;Acc:F4IB63] |
| AT1G13245 | RTFL17 | 0.65 | 1.13 | 0.064 | 0.013 | At1g13245 [Source:UniProtKB/TrEMBL;Acc:Q9SAF8] |

| | | | | | | |
|---|---|---|---|---|---|---|
| AT3G23880 | - | 0.12 | 1.13 | 0.549 | 0.042 | F-box/kelch-repeat protein At3g23880 [Source:UniProtKB/Swiss-Prot;Acc:Q9LIR8] |
| AT1G24440 | - | 0.49 | 1.12 | 0.052 | 0.011 | F21J9.10 [Source:UniProtKB/TrEMBL;Acc:Q9FYL9] |
| AT3G19290 | ABF4 | 0.08 | 1.12 | 0.684 | 0.008 | ABRE binding factor 4 [Source:UniProtKB/TrEMBL;Acc:F4JB53] |
| AT2G37950 | - | -0.12 | 1.11 | 0.887 | 0.021 | At2g37950 [Source:UniProtKB/TrEMBL;Acc:Q5XEN4] |
| AT1G19400 | - | 0.44 | 1.10 | 0.160 | 0.039 | Erythronate-4-phosphate dehydrogenase family protein [Source:UniProtKB/TrEMBL;Acc:Q8VYC6] |
| AT2G18700 | TPS11 | 0.37 | 1.10 | 0.160 | 0.012 | Probable alpha,alpha-trehalose-phosphate synthase [UDP-forming] 11 [Source:UniProtKB/Swiss-Prot;Acc:Q9ZV48] |
| AT4G37180 | HHO5 | 0.94 | 1.10 | 0.002 | 0.045 | Transcription factor HHO5 [Source:UniProtKB/Swiss-Prot;Acc:F4JRB0] |
| AT4G36850 | - | 0.21 | 1.10 | 0.485 | 0.032 | PQ-loop repeat family protein / transmembrane family protein [Source:UniProtKB/TrEMBL;Acc:Q94AH7] |
| AT5G56180 | ARP8 | 0.45 | 1.09 | 0.056 | 0.005 | Actin-related protein 8 [Source:UniProtKB/Swiss-Prot;Acc:Q9FKT0] |
| AT4G37470 | KAI2 | 0.45 | 1.08 | 0.007 | 0.000 | KAI2 [Source:UniProtKB/TrEMBL;Acc:A0A178V1E5] |
| AT1G80420 | XRCC1 | 0.82 | 1.08 | 0.010 | 0.007 | DNA-repair protein XRCC1 [Source:UniProtKB/Swiss-Prot;Acc:Q24JK4] |
| AT3G19930 | STP4 | 0.56 | 1.07 | 0.061 | 0.010 | STP4 [Source:UniProtKB/TrEMBL;Acc:A0A178V5Z0] |
| AT1G76240 | - | -1.11 | 1.07 | 0.056 | 0.035 | At1g76240 [Source:UniProtKB/TrEMBL;Acc:Q501A3] |
| AT2G16365 | - | 0.39 | 1.07 | 0.131 | 0.016 | F-box protein At2g16365 [Source:UniProtKB/Swiss-Prot;Acc:Q84V03] |
| AT5G13730 | SIGD | 0.18 | 1.06 | 0.109 | 0.001 | RNA polymerase sigma factor sigD, chloroplastic [Source:UniProtKB/Swiss-Prot;Acc:Q9ZSL6] |
| AT2G45660 | SOC1 | 0.84 | 1.06 | 0.005 | 0.002 | SOC1 [Source:UniProtKB/TrEMBL;Acc:A0A178VZL4] |
| AT1G29670 | - | 0.22 | 1.05 | 0.193 | 0.007 | GDSL-like Lipase/Acylhydrolase superfamily protein [Source:TAIR;Acc:AT1G29670] |
| AT4G27560 | UGT79B2 | 0.28 | 1.05 | 0.333 | 0.039 | UDP-glycosyltransferase 79B2 [Source:UniProtKB/Swiss-Prot;Acc:Q9T080] |
| AT1G78995 | - | 0.67 | 1.05 | 0.075 | 0.026 | At1g78995 [Source:UniProtKB/TrEMBL;Acc:Q8L5A1] |
| AT1G67660 | - | 0.83 | 1.05 | 0.053 | 0.033 | Restriction endonuclease, type II-like superfamily protein [Source:UniProtKB/TrEMBL;Acc:Q8GW93] |
| AT1G43670 | CYFBP | 0.85 | 1.04 | 0.018 | 0.004 | Fructose-1,6-bisphosphatase, cytosolic [Source:UniProtKB/Swiss-Prot;Acc:Q9MA79] |
| AT5G18680 | TULP11 | 0.34 | 1.04 | 0.319 | 0.009 | Uncharacterized protein At5g18680 (Fragment) [Source:UniProtKB/TrEMBL;Acc:C0SVQ0] |
| AT5G54280 | VIII-2 | 0.11 | 1.03 | 0.740 | 0.042 | Myosin-2 [Source:UniProtKB/Swiss-Prot;Acc:F4K0A6] |
| AT5G19140 | ATAILP1 | 0.97 | 1.03 | 0.002 | 0.046 | Aluminum induced protein with YGL and LRDR motifs [Source:UniProtKB/TrEMBL;Acc:Q94BR2] |
| AT5G05140 | MED26B | 0.94 | 1.03 | 0.009 | 0.004 | Probable mediator of RNA polymerase II transcription subunit 26b [Source:UniProtKB/Swiss-Prot;Acc:Q9FHK9] |
| AT3G23480 | - | 0.81 | 1.03 | 0.025 | 0.031 | Cyclopropane-fatty-acyl-phospholipid synthase [Source:UniProtKB/TrEMBL;Acc:F4J434] |
| AT1G68490 | - | -0.17 | 1.03 | 0.133 | 0.003 | At1g68490 [Source:UniProtKB/TrEMBL;Acc:Q9CA32] |
| AT5G62030 | - | 0.65 | 1.03 | 0.010 | 0.027 | Diphthamide synthesis DPH2 family protein [Source:UniProtKB/TrEMBL;Acc:Q8RWW3] |
| AT5G19530 | ACL5 | 0.72 | 1.02 | 0.022 | 0.016 | Thermospermine synthase ACAULIS5 [Source:UniProtKB/Swiss-Prot;Acc:Q9S7X6] |
| AT2G02710 | TLP1 | 0.62 | 1.02 | 0.036 | 0.009 | Protein TWIN LOV 1 [Source:UniProtKB/Swiss-Prot;Acc:O64511] |
| AT3G10720 | PME25 | 0.41 | 1.01 | 0.097 | 0.009 | Probable pectinesterase/pectinesterase inhibitor 25 [Source:UniProtKB/Swiss-Prot;Acc:Q94CB1] |
| AT5G56860 | GATA21 | -0.34 | 1.01 | 0.113 | 0.034 | GATA transcription factor 21 [Source:UniProtKB/Swiss-Prot;Acc:Q5HZ36] |
| AT3G26290 | CYP71B26 | 0.84 | 1.01 | 0.088 | 0.021 | Cytochrome P450, family 71, subfamily B, polypeptide 26 [Source:UniProtKB/TrEMBL;Acc:A0A1I9LRI3] |
| AT5G06410 | - | 0.36 | 1.01 | 0.377 | 0.027 | DNAJ heat shock N-terminal domain-containing protein [Source:UniProtKB/TrEMBL;Acc:Q8L7K4] |
